# *Pseudomonas* volatiles shape the root transcriptome and microbiome to promote plant growth under drought

**DOI:** 10.64898/2026.01.27.701981

**Authors:** Zulema Carracedo Lorenzo, Muhammad Syamsu Rizaludin, Jielin Wang, Roland Berdaguer, Cristina Brito-López, Carlos Sánchez Arcos, Paolina Garbeva, Corné M.J. Pieterse, Marcel Dicke, Christa Testerink, Karen J. Kloth, Rumyana Karlova

## Abstract

- Volatile organic compounds (VOCs) emitted by soil bacteria influence interactions with other soil microbes and with plant roots. While their potential as plant-growth promoters is well recognized, their role in promoting plant resilience to abiotic stress and the underlying molecular mechanisms remains poorly understood. Here, we investigate the role of *Pseudomonas* VOCs in enhancing plant resilience to drought stress.
- *Arabidopsis thaliana* plants were exposed to VOCs emitted by *Pseudomonas* strains under both control and osmotic-stress conditions. VOC exposure generally enhanced plant growth, and this effect was even more pronounced under both drought and salt stress. Transcriptomic analysis revealed that VOC exposure modulates key stress-responsive pathways, including those related to abscisic acid biosynthesis and signalling, sugar transport, iron uptake, aliphatic glucosinolate biosynthesis, and plant defences. Using Arabidopsis mutants, we identified abscisic acid and aliphatic glucosinolates as important components in mediating the plant response to VOCs. SWEET11/12 sugar transporters and ABA signaling genes were downregulated by VOCs exposure, in order to allow for a positive regulation of lateral root numbers (in case of SWEET genes) and plant growth in general under drought stress. In summary, using metabolomics, transcriptomics and functional analysis, we showed a negative cross-talk between the effects of VOCs on plant growth and glucosinolate production, whereas a positive interaction was observed between the biosynthesis of coumarins and VOCs.
- Notably, VOCs also improved drought tolerance in soil-grown *Brassica oleracea* plants. We showed that VOC treatment altered the root-associated microbiome under drought, leading to a community composition more similar to that of well-watered plants.
- Our results show that *Pseudomonas* emitted VOCs can promote plant growth under drought conditions, linked to root transcriptional reprogramming and direct or indirect microbiome modulation.

## Introduction

Plants are exposed to a variety of biotic and abiotic stresses, such as pathogens, insect pests and drought. They are anchored by their roots in the soil, a place for water and nutrients uptake, but also a rich niche for plant-microbe interactions. Beneficial soil microorganisms have been shown to improve plant growth (plant-growth promoting rhizobacteria, PGPR) and to contribute to plant health when plants are subjected to (a)biotic stresses (Porter et al., 2020; Rubin et al., 2017). Drought is one of the most severe abiotic stresses for plants and represents a threat to food security (Gupta et al., 6488).

To cope with water deficit, plants have developed complex adaptative mechanisms that involve morpho-anatomical, biochemical and molecular responses to reduce water loss and preserve their hydraulic status (Zahedi et al., 2025). Fine-tuning of phytohormone signalling is key for overcoming water deficit, with abscisic acid (ABA) playing a central role. In *Arabidopsis thaliana*, drought stress triggers increased ABA levels, particularly in the stomata and vasculature tissue (Karlova et al., 2021; Mo et al., 2024). Through activation of *SNF1-related protein kinases 2* (*SnRK2*), ABA activates a cascade of genes leading to stomatal closure and an improved plant water balance (Gupta et al., 6488). In addition, it has been shown that phosphorylation of the sugar transporters *SWEET11* and *SWEET12* by ABA-activated *SnRK2*s improves the transport of sucrose over long distances to the roots, a vital process for building a resilient root system when facing drought conditions (Chen et al., 2022).

The interplay of ABA and other phytohormones also determine the accumulation and conversion of secondary metabolites that play a key role in resilience to drought. Glucosinolates are a well-known example of this in the Brassicaceae family (Abuelsoud et al., 2016). In Arabidopsis, aliphatic glucosinolate levels are maintained by auxin-sensitive Aux/IAA repressors when plants are under drought conditions, and regulate stomatal closure, presumably via isothiocyanate-mediated ROS generation in or near the stomata (Salehin et al., 2019). Other studies also showed that upregulation of the aliphatic glucosinolate biosynthesis genes *branched-chain aminotransferase 4* (*BCAT4*) and *methylthioalkylmalate synthase* (*MAM*) 1 and 3, increase tolerance to drought and elevated CO_2_ (AbdElgawad et al., 2023). Additionally, aliphatic glucosinolates have been shown to accumulate in plant roots and be exuded into the rhizosphere, where they play a role in shaping the rhizosphere microbiome (Bressan et al., 2009; Chroston et al., 2024). However, it is not clear in the rhizosphere if the exuded glucosinolates are intact or processed in possible breakdown products already in the roots or these breakdown products are generated due to enzymatic activity of the rhizosphere microbiome in the soil leading to changes in the microbiome composition at the rhizosphere.

In recent decades, the holobiont perspective has gained increasing recognition, based on the concept that plant responses to (a)biotic stresses result from the complex and intertwined interactions between the plant and its microbiome (Ali et al., 2024; Trivedi et al., 2022). This perspective has led to new insights into the role of root microbiome in the alleviation of drought stress. Drought stress responses cause strong alterations in the root microbiome, either directly, through the proliferation of drought-resilient microbial taxa (Metze et al., 2023), or indirectly, via drought-induced changes in root exudate patterns (De Vries et al., 2019). Secondary metabolites in these exudates, such as glucosinolates, act as modulators of the root microbiome (Bressan et al., 2009; Chroston et al., 2024). Typical drought-induced alterations in the composition of the root microbiome include the enrichment of Actinobacteria and depletion of Proteobacteria (Fitzpatrick et al., 2018; Naylor et al., 2017; Santos-Medellín et al., 2021; Berdaguer et al., 2024). These shifts in the microbial community are believed to contribute to enhanced plant drought resilience, for example by maintaining plant iron homeostasis (Xu et al., 2021). This is particularly important as iron availability, along with nitrogen, is among the nutrients most affected by drought. Its uptake depends on soil water content and the expression of plant transporters that are downregulated during drought conditions (Kanwar et al., 2021; Xu et al., 2021).

Members of the root-associated microbiota, including PGPR, produce a wide range of volatile organic compounds (VOCs), some of which have been shown to positively impact plant health (Blom et al., 2011; Garbeva & Weisskopf, 2020). VOCs can modulate plant growth by inducing different molecular pathways within the plant or by inhibiting the proliferation of pests or pathogens that are harmful to the plant (Fincheira & Quiroz, 2018; Garbeva & Weisskopf, 2020). To date, only for a few bacterial VOCs, the molecular mechanism of action in the plant have been elucidated. For example, indole, a VOC reported to be produced by many rhizobacteria and growth promoting fungi, promotes plant growth by modulating auxin signalling (Bailly et al., 2014; Contreras-Cornejo et al., 2009). Similarly, 2R,3R-butanediol, a VOC produced by *Pseudomonas chlororaphis* O6, enhances tolerance to drought in *Arabidopsis thaliana* (Cho et al., 2008).

One of the most extensively studied plant-growth-promoting rhizobacteria (PGPR) is *Pseudomonas simiae* WCS417 (*Ps*WCS417) (Pieterse et al., 2021). Since its first discovery in 1988, when it was isolated from wheat roots, *Ps*WCS417 has served as a model to unravel the molecular mechanisms underlying plant-growth promotion (Pieterse et al., 2021). Under axenic conditions, *Ps*WCS417 increases plant biomass and promotes the development of lateral roots and root hairs, enhancing the capacity of the plants to absorb water and nutrients (Zamioudis et al., 2013). This growth-promoting effect has been attributed to enhanced auxin signalling in *Arabidopsis thaliana*, as evidenced by the lack of response in the auxin triple mutant *tir1 afb2 afb3* to *Ps*WCS417 treatment. Furthermore, sugar transport, specifically the genes *SWEET11* and *SWEET12*, has a role in the interaction between the plant and *Ps*WCS417, most likely by controlling the amount of sugar that is transported from the shoot to the root and into the rhizosphere (Desrut et al., 2020).

Beyond promotion of plant growth, *Ps*WCS417 contributes to plant health through the activation of induced systemic resistance (ISR) that is effective against a broad spectrum of plant pathogens (Pieterse et al., 2014). *Ps*WCS417-ISR is not only effective in Arabidopsis (Pieterse et al., 1996) but also in crop species such as banana (Nel et al., 2006) and grapevine (Verhagen et al., 2010), among many others. More recent studies have shown that root colonization by *Ps*WCS417 also enhances plant resilience to abiotic stresses. For example, Chiappero et al., (2019) showed that *Ps*WCS417 inoculation increased drought tolerance in peppermint plants by enhancing their antioxidant capacity. Similarly, *Ps*WCS417 has been shown to promote growth under alkaline conditions in tomato (Aparicio et al., 2025). These findings underscore the potential of *Ps*WCS417 to improve resilience across various plant species and under diverse biotic and abiotic stress conditions.

While most studies have focused on direct root inoculation, it is now evident that VOCs produced by *Ps*WCS417 can also promote plant growth and modify root architecture, without requiring physical contact between the plant and the bacteria (Blom et al., 2011; Desrut et al., 2020; Zamioudis et al., 2013). Although *Ps*WCS417 has proven to be an excellent model for studying plant-microbe interactions (Pieterse et al., 2021), there is still limited knowledge regarding the effects of its VOCs on plant resilience to abiotic stress, their influence on the composition of rhizobacterial communities, and their role in modulating plant growth under well-watered and drought conditions.This study investigates whether VOCs emitted by *Pseudomonas* spp. enhance plant resilience to drought and aims to identify the mechanisms underlying this effect. We hypothesize that Pseudomonas VOCs (*Ps*VOCS), specifically those from the PGPB *P. simiae* WCS417 (PsWCS417), *P. simiae* WCS315 (*Ps*WCS315) and *P. protegens* CHA0 (*Ps*CHA0), prime plant drought tolerance responses by inducing specific transcriptional reprograming. To test this hypothesis, we analysed *Ps*VOCs-induced transcriptional changes in axenically grown *Arabidopsis thaliana* and used Arabidopsis mutants and metabolomics to dissect the molecular mechanism involved in the response to *Ps*VOCs. We further investigate whether this effect is conserved in soil-grown *Brassica oleracea* and whether *Ps*VOCs contribute to drought resilience by altering the rhizosphere microbial community, which we examined using metabarcoding of *B. oleracea* roots exposed to *Ps*VOCs in soil. Finally, we sought to identify candidate volatile compounds responsible for enhanced drought resilience through GC-MS analyses.

## Materials and Methods

### 1. Arabidopsis exposure to *Pseudomonas* VOCs in sterile media

#### 1.1 Plant Material

*Arabidopsis thaliana* accession Columbia (Col-0) was used as a wild-type control. The Arabidopsis mutants *bcat4-1* (SALK_013627), *sweet11/12* (N68845) and *snrk2*.*2/2*.*3* (GABI_807G04/ SALK_107315), all in Col-0 background, were obtained from the European Arabidopsis Stock Centre (NASC). Homozygous T-DNA plants were selected by PCR (Table S1) and seeds were harvested for subsequent experiments.

#### 1.2 Plant growth and treatments

For growth in sterile media, Arabidopsis seeds were surface sterilized for 5 min with 1.5% bleach and 0.02% Triton X-100 solution and washed 10 times with sterile MiliQ water. Seeds were sown in Petri dishes containing half strength Murashige and Skoog (MS) (Duchefa Biochemie), 0.1% w/v 2-(N-morpholino) ethanesulfonic acid (MES) and 1% w/v Daishin agar (Duchefa Biochemie) (pH 5.8, adjusted with KOH). After three days of stratification at 4 °C, Petri dishes were transferred to racks at a 70-degree angle in a climate chamber (22°C, 16 h light, the light intensity ranged from 80 to 200 µmol/m^2^/s, RH 60%). Uniform 5-day-old seedlings were carefully transferred to new Petri dishes containing either control or drought media. The control media contained half-strength MS (Duchefa Biochemie), 0.1% MES w/v and 1% w/v Daishin agar (Duchefa Biochemie) (pH 5.8, adjusted with KOH). Drought treatment media were additionally supplemented with 150 mM sorbitol (Duchefa Biochemie). Salt stress treatment was induced with 75mM NaCl. Experiments involving root architecture phenotyping were performed in square dishes (120 x 120 x 17 mm). In the experiment where Arabidopsis Col-0 was treated with the three different *Pseudomonas* strains, the plants and the bacteria grew in the same media (Fig.1A, Fig. S1). For the experiment where we investigated the effect of *Ps*WCS417 VOCs in different Arabidopsis mutants (Fig. 3, Fig. S1), a 0.5 cm strip of media was cut to ensure the physical separation between the plant and the bacteria. Additionally, in that experiment the bacteria were always grown in half-strength MS supplemented with 150 mM of sorbitol. In all experiments performed in square dishes, five seedlings were grown per dish, and five dishes were used per treatment. For the RNA-seq experiment (Fig. 2, Fig S1), two-compartment round dishes (100 mm diameter) with a centre partition were used, and ∼ 30 seedlings were grown per dish.

**Figure 1.**
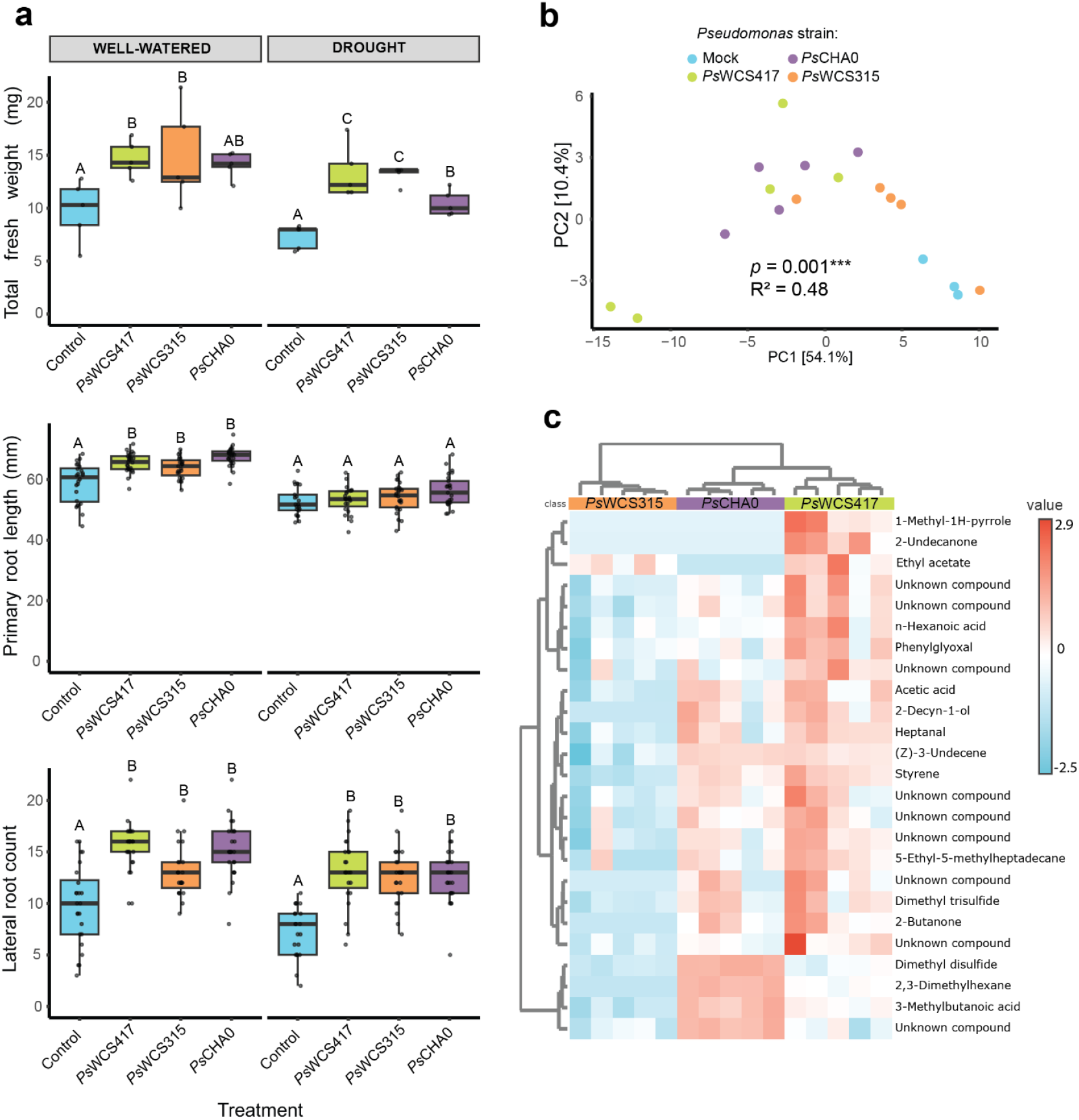
Effect of *Ps*VOCs on Arabidopsis drought tolerance in sterile media. (a) Total fresh weight of shoot and root (top), primary root length (middle), and lateral root count (bottom) of Arabidopsis seedlings co-cultivated with different *Pseudomonas* strains (*Ps*WCS417, *Ps*WCS315 and *Ps*CHA0) or mock under either control (half-strength MS) or drought (half-strength MS with sorbitol) conditions. Boxplots display median, interquartile range, and data distribution. Different letters indicate statistically significant differences among treatments (Tukey’s HSD, P < 0.05, fresh weight: n= 5, root system: n= 25). (b) Principal Component Analysis (PCA) of VOC profiles emitted by *Pseudomonas* strains, showing distinct clustering of strains based on their VOC composition (PERMANOVA, p <0.001, R2 = 0.479, n = 3-5). (c) Heatmap of VOCs emitted by *Pseudomonas* strains. Each row represents an individual VOC, and colours indicate the relative abundance (red: high, blue: low). Hierarchical clustering groups strains based on their emitted VOCs profiles.

**Figure 2.**
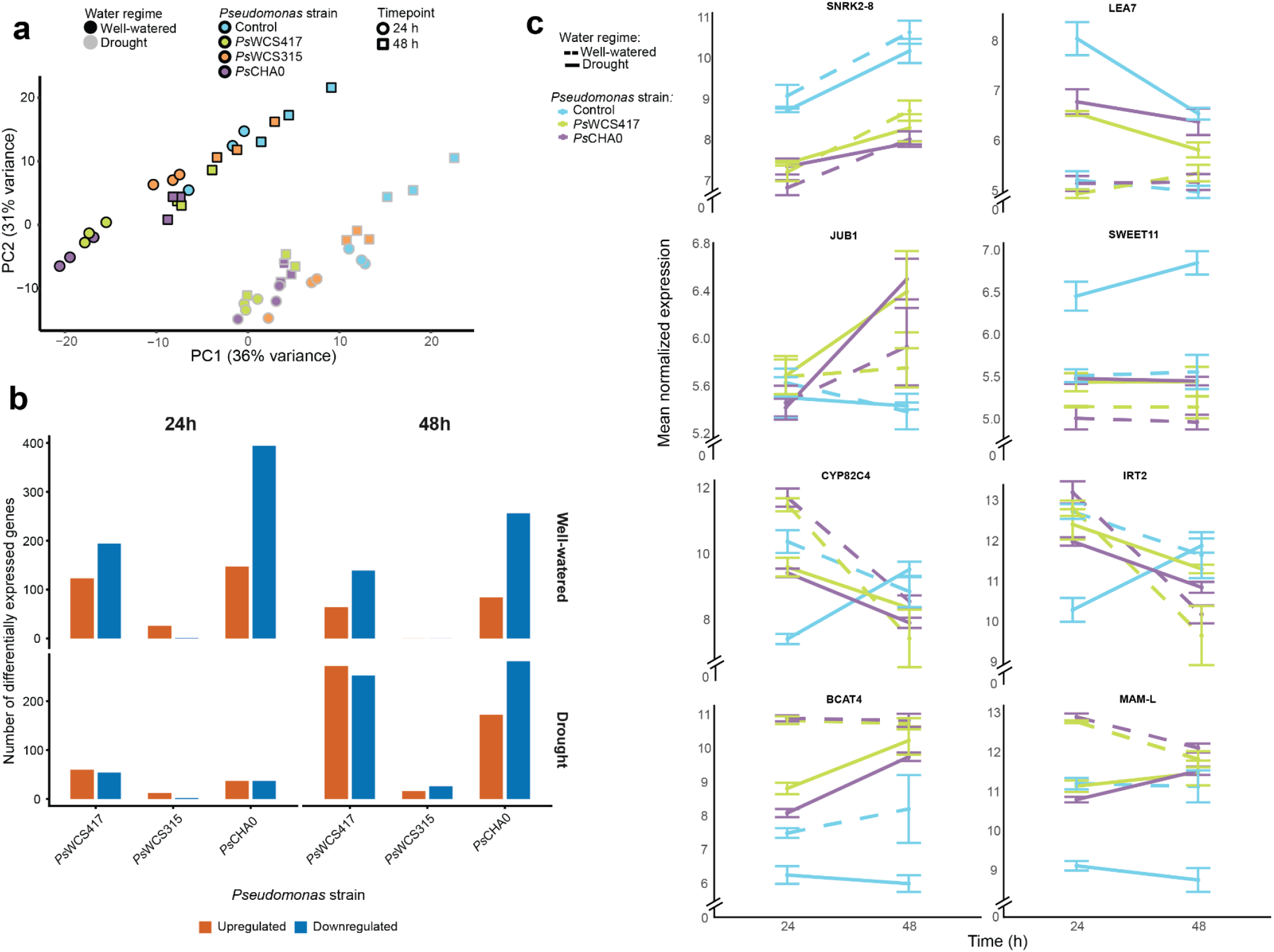
Transcriptomic analysis of Arabidopsis roots exposed to *Pseudomonas* VOCs under well-watered and drought conditions in sterile media. (a) Principal Component Analysis (PCA) of Arabidopsis root transcriptomic profiles in response to *Pseudomonas* VOCs under well-watered and drought conditions. Fill colours represent different *Pseudomonas* strains (mock, *Ps*WCS417, *Ps*WCS315, *Ps*CHA0), stroke colour indicates water regime (well-watered, drought) and shapes represent the different time points (24 h, 48 h, n= 3). (b) Bar plot showing the number of differentially expressed genes (DEGs) upregulated and downregulated in response to each *Pseudomonas* VOCs treatment, water regime, and timepoint. Genes were considered differentially expressed at false discovery rate (FDR) < 0.05 and │fold2change│ ≥ 1. Color indicate gene regulation: red bars represents upregulated genes, and blue bars represents downregulated genes. Abbreviations: *Ps*WCS417, *P. simiae* WCS417; *Ps*WCS315, *P. simiae* WCS315; *Ps*CHA0 = *P. protegenes* CHA0. (c) Expression dynamics of selected drought-responsive genes (*SNRK2-8, LEA7, JUB1, SWEET11, CYP82C4, IRT1, BCAT4* and *MAM-L*) in Arabidopsis roots exposed to mock, *Ps*WCS417 or *Ps*CHA0 VOCs at 24 and 48 h. Dashed and solid lines represent well-watered and drought treatments, respectively. Error bars indicate standard error of the mean.

**Figure 3.**
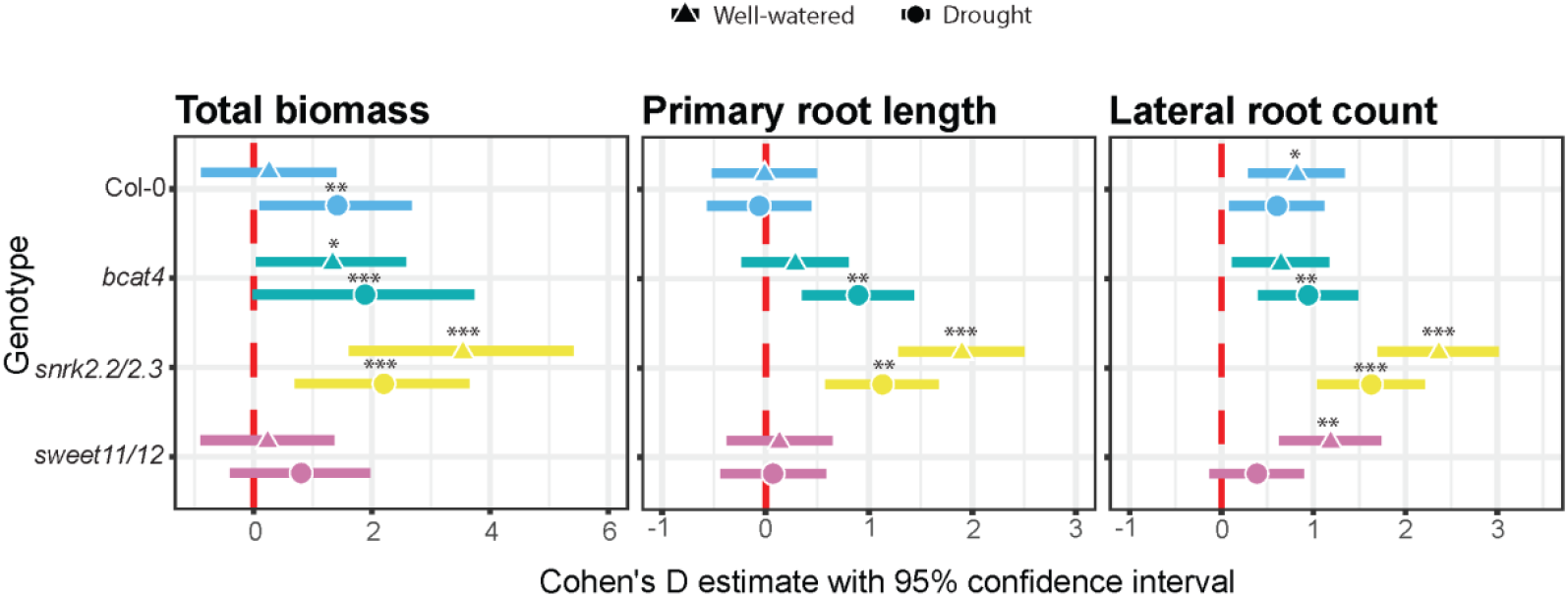
Effect of *Ps*WCS417 VOCs in Col-0, *bcat-4, snrk2.2/2.3* and *sweet11/12* Arabidopsis lines growth. Forest plots show Cohen’s D size estimates with 95% confidence intervals of *Ps*WCS417 VOCs on the total biomass, primary root length and lateral root count of the different Arabidopsis lines seedlings under two treatments: well-watered (triangle) and drought (circle), compared to mock treated seedlings. Values above the dashed red line indicate an increase in the *Pseudomonas* treatment. Significant differences due to *Ps*WCS417 VOCs expositions within each line and water regime are indicated (GLMM, ANOVA II test; * = p < 0.05; ** = p < 0.01; *** = p < 0.001).

#### 1.3 Cultivation of *Pseudomonas* strains and inoculation

The *Pseudomonas* strains *Pseudomonas simiae* WCS417, *Pseudomonas simiae* WCS315 and *Pseudomonas protegens* CHA0 were cultured in King’s B (KB) media agar dishes (Duchefa Biochemie) and incubated for 24 h at 28°C (Zamioudis et al. 2013). Cells were collected in 10 mM MgSO_4_ and washed twice by centrifuging at 4000*g*. Then, the bacteria were re-suspended in 10 mM MgSO_4_ to an optical density at 600 nm (OD600nm) of 0.002. For the experiments in square Petri dishes, 240 µl of the bacterial suspensions, or 10 mM MgSO_4_ for the mock treatment, were spotted at the lower end of the dish. For experiments performed in two-compartment round dishes, 120 µl of bacterial suspension was applied in the compartment next to the plant compartment.

#### 1.4 Phenotypic Data Collection and Analysis

For plant root architecture phenotyping, dish images were collected with a scanner at two time points: (1) the day of the seedling transfer and inoculation and (2) seven or nine days after co-cultivation with the different *Pseudomonas* strains. Primary root growth, total root length and number of lateral roots were assessed with Rootnav software (Pound et al., 2013). For shoot and root fresh biomass, the total biomass of the seedlings of each dish was measured on an Mettler AE 163 analytical balance (Mettler-Toledo, Switzerland) with a readability of 0.1 mg, 10 days after co-cultivation. Then, seedlings were sectioned at the root-shoot junction and the biomass of the shoots was measured. The root fresh biomass was calculated by subtracting the shoot fresh biomass from the total fresh weight.

For the experiment where we tested the effect of different *Pseudomonas* strains under well water and drought conditions, statistical analyses were done with a generalized linear mixed model (GLMM) using the glmmTMB package (Brooks et al., 2017; McGillycuddy et al., 2025), with *Pseudomonas* strain as a fixed effect and dish as a random effect to account for experimental variation among Petri dishes. Data distribution were assessed and adjusted for each data subset. Post-hoc pairwise comparisons between treatments were performed using estimated marginal means (emmeans) (Lenth et al., 2025), and adjusted using Tukey. Significance letters were calculated with multicomp package (Hothorn et al., 2008). For the experiments where we tested the effect of PsWCS417 on the resilience to different drought intensities and salt, data were analyses using Student t-test (normal distribution) and Wilcoxon rank-sum test (non-normal distribution).

#### 1.5 Total RNA extraction

Arabidopsis root samples for gene expression analysis were harvested 24 h (time point 1) or 48 h (time point 2) after five-day-old seedlings were transferred to Petri dishes and exposed to *Ps*VOCs.. The seedlings were excised at the shoot-root junction, and roots were immediately frozen in liquid nitrogen, and then stored at – 80 °C (*n* = 3 pools of 60 roots). Total RNA was isolated with the FavorPrep Plant Total RNA Mini Kit (FAVORGEN). RNA library preparation and transcriptome sequencing were conducted by Novogene UK Co. Ltd (Cambridge, UK) on a NovaSeq 6000 platform, as described by Nicolas et al., (2025).

#### 1.6 RNA-seq data analysis

RNA-seq data were pre-processed with nf-core pipeline (Ewels et al., 2020). Differential expression analysis was performed with the DESeq2 package (Love et al., 2014). Genes with a false discovery rate (FDR) < 0.05 and a |fold2change| ≥ 1, were considered differentially expressed and used for further analyses. Differential expression was assessed separately for each treatment condition (well-watered and drought) and time point (24 h and 48 h). Principle Component Analysis (PCA) was performed on variance-stabilized count data using the plotPCA function from DESeq2. To quantify the effects of experimental factors on multivariate gene expression patterns, a PERMANOVA was conducted using adonis2 function from the vegan package (Oksanen et al., 2009), followed by post hoc pairwise comparisons using the pairwise.adonis2 function. To visualized the expression pattern of differentially significant genes across the different groups a heatmap was built using pheatmap package (Kolde, 2025). For the heatmap, a threshold of FDR < 0.05 and |fold2change| ≥ 2 for at least one of the groups was used. GO enrichment analysis was performed using the clusterProfiler package (Yu, 2024).

#### 1.7 Soluble Sugars Measurement

Arabidopsis root and shoot samples for sugar analysis were harvested 7 days after exposure to *Ps*VOCs grown under well-watered or drought-stress conditions. The extraction of soluble sugars and measurements were done as published before (Dinh et al., 2019). Data were analyzed using a two-way ANOVA, followed by Tukey’s post hoc test.

#### 1.8 Glucosinolates and Coumarins extraction and analysis by UHPLC–HRMS

Arabidopsis roots collected as described above for sugar measurements were ground in liquid nitrogen to a fine powder. Root extracts were prepared by adding extraction solvent (99.867% MeOH acidified with 0.133% FA) at the 5:1 solvent: root tissue ratio, vortexing for 10 sec, sonicating in an ultrasonic bath for 10 min, centrifuging for 3 min at 21000 x g and the resulting supernatant was used for LC-MS analysis.

Intact glucosinolates (GSLs) and coumarins were analyzed by ultra-high-performance liquid chromatography coupled to high-resolution mass spectrometry (UHPLC–HRMS). Analyses were performed using a Vanquish™ Horizon UHPLC system interfaced with an Exploris™ 120 Orbitrap mass spectrometer (Thermo Fisher Scientific, Dreieich, Germany). For each sample, 5 µL of extract was injected onto an Acquity UPLC BEH C18 column (1.7 µm particle size, 2.1 × 150 mm; Waters) maintained at 40 °C. Chromatographic separation was conducted at a constant flow rate of 400 µL min^−1^ using a binary mobile phase system consisting of 0.1% (v/v) formic acid in water (mobile phase A) and 0.1% (v/v) formic acid in acetonitrile (mobile phase B). The gradient elution program was as follows: isocratic 5% B from 0.0 to 1.0 min; linear increase to 75% B from 1.0 to 22.0 min; linear ramp to 90% B from 22.1 to 23.0 min; isocratic hold at 90% B from 23.1 to 26.0 min; followed by re-equilibration to 5% B from 26.1 to 27.0 min.

Mass spectrometric detection was performed in negative electrospray ionization (ESI^−^) mode, which was selected to ensure robust ionization of intact GSLs and coumarins. Full-scan MS data were acquired over an m/z range of 90–1350 at a resolving power of 60,000 (m/Δm, at m/z 200). The ESI source parameters were set to a spray voltage of 3.0 kV and a capillary temperature of 290 °C. Chromatographic peak detection and integration for both GSLs and coumarins were performed using the Qual Browser module of Xcalibur software (version 4.1, Thermo Fisher Scientific). Because all plant extracts were prepared using an identical solvent-to-sample mass ratio, LC–MS peak areas were used as semi-quantitative measures and directly compared across samples.

For structural annotation and confirmation, pooled quality-control samples were analyzed using combined full-scan MS and data-dependent MS/MS acquisition (Full MS/dd-MS^2^, top-N) in negative ionization mode. Full-scan MS parameters included a resolving power of 60,000 (m/Δm), an automated gain control (AGC) target of 1 × 10^6^, a maximum injection time of 100 ms, and an m/z acquisition range of 90–1350. Data-dependent MS^2^ spectra were acquired for the four most intense precursor ions per scan at a resolving power of 15,000 (m/Δm), using an AGC target of 1 × 10^5^, a maximum injection time of 118 ms, and an isolation window of 1.5 m/z. Fragmentation was achieved using stepped normalized collision energies (NCE) of 20, 40, and 100 eV to generate diagnostic fragment ions supporting the annotation of both intact glucosinolates and coumarin.

LC-MS peak areas for the GSLs and coumarins were log-transformed and Pareto-scaled, and significant differences among treatments were assessed using ANOVA (p < 0.05). Tukey post hoc comparisons were then conducted, and group significance letters were assigned. Different letters indicate significant differences among the treatments.

### 2. *Brassica oleracea* exposure to *Pseudomonas* VOCs in soil

#### 2.1 Plant material

*Brassica oleracea* var. Rivera uncoated seeds were obtained from Bejo (Warmenhuizen, The Netherlands). *B. oleracea* roots were exposed to *Ps*VOCs by using a two-compartment pot system (Moisan et al., 2020). The two-compartment pot system consisted of an upper compartment for the plant and soil, and a lower compartment for the different volatiles treatments. Both compartments were separated by a nylon membrane of 1 μm mesh width (Plastok associates Ltd., Birkenhead Wirral, UK), which allowed the volatile exchange between the two compartments, but no physical contact.

#### 2.2 Plant growth and treatments

*Brassica oleracea* seeds were stratified on wet filter paper at 4°C for three days and then sown in a germination tray containing a soil mixture (potting soil:sand (1:1, vol/vol)). The potting soil was obtained from (Lensli BV) and no additional fertilizers were added. After six days, seedlings were transferred to the upper compartment of the two-compartment pot system, with all pots containing the same weight of soil mixture. Soil samples were weighed before and after oven drying in order to calculate the actual soil mass added to the pots. This was used to calculate the amount of water needed to be added to reach the desired gravimetric water content. Plants were grown in a climate chamber under a long day photoperiod (16 h light), 22 °C and 65 % relative humidity. Seven plants were grown per treatment. All plants were watered to 80% gravimetric water content (gwc) for 12 days. After that, half of the pots were kept in well-watered control conditions (70 % gwc) and the other half in drought conditions (30 % gwc). On the same day, roots started to be exposed to the *Ps*VOCs treatment. For that, KB media (Duchefa Biochemie) containing Petri dishes were inoculated with 200 µl of bacterial suspension of *Ps*WCS417, *Ps*WCS315 or *Ps*CHA0 or with 200 µl 10 mM MgSO_4_ (mock). Strain suspensions were prepared as described above, with the bacterial titer adjusted to an OD600 nm of 0.5. After incubation for two days at 22°C, one Petri dish was placed in each lower compartment of a two-compartment pot system. Petri dishes containing the *Pseudomonas* strains were replaced every 2-3 days, while control dishes were replaced every day to avoid contamination with microorganisms. After 4 weeks of drought treatment and exposure to *Ps*VOCs, plants were harvested. Shoots were excised from the roots and fresh and dry weights were measured on an analytical balance. For the dry weight, shoots were dried at 70 °C for three days. Soil samples were taken from each pot and stored at −20°C until further processing for microbiome analyses.

#### 2.3 Data analysis

Statistical analysis was performed with the non-parametric Kruskal-Wallis test, followed by the pairwise analysis with the Mann Whitney U test.

#### 2.4 Microbiome DNA extraction and amplicon sequencing

DNA from *B. oleracea* soil samples was extracted using a modified protocol of Qiagen’s DNeasy PowerSoil Kit, by adjusting the volumes of the reagents for larger samples (van Bentum et al., 2025). 16S libraries were prepared and sequenced by BGI Genomics (Hong Kong). The V3-V4 region was amplified using the universal primers 341F (CCTACGGGNGGCWGCAG) and 805R (GACTACHVGGGTATCTAATCC), with addition of PNA clamps to block amplification of mitochondrial and plastid DNA (mPNA: GGCAAGTGTTCTTCGGA, pPNA: GGCTCAACCCTGGACAG). Samples were multiplexed and paired-end reads of 250 bp were sequenced on a NovaSeq (Illumina).

#### 2.5 Amplicon sequence processing and analysis

Demultiplexed reads were processed to amplicon sequence variants (ASVs) using a modified DADA2 pipeline optimized for NovaSeq data (Holland-Moritz et al.; Callahan et al., 2016). Notably, instead of assigning taxonomy using DADA2’s assignTaxonomy function, we used the function classify-sklearn in Qiime2 version 2022.2 (Caporaso et al., 2010). For 16S amplicons, we used a custom-made classifier based on the Silva database, version 138.1 (Quast et al., 2013), filtered and trimmed to our V3-V4 primers with Rescript in Qiime2.

ASVs assigned to Archaea, mitochondria or chloroplasts were removed. The Decontam package (Davis et al., 2018) was used to remove contaminants. ASVs with less than 380 bp were removed, as well as samples with less than 10,000 reads. After these cleaning and filtering steps, 48,027 bacterial ASVs were retained.

Alpha-diversity was assessed with Shannon index, Simpson index and Observed richness, all calculated with the Phyloseq package (McMurdie & Holmes, 2013), and statistically evaluated with ANOVA or Kruskal-Wallis, depending on the data distribution. For beta-diversity, data were further filtered by retaining only ASVs with a minimum relative abundance of 0.001% and present in at least 50% of the samples, per water regime and *Pseudomonas* strain treatment combination. Additionally, data were normalized by cumulative sum scaling (CSS) using the metagenomeSeq package (Paulson et al., 2013). Bray-Curtis dissimilarities were calculated with the vegan package (Oksanen et al., 2009), and differences in microbial community composition between water regimes and *Pseudomonas* strain treatments were tested with PERMANOVA using the adonis2 function from the vegan package. Non-metric Multi-Dimensional Scaling (NMDS) ordination was calculated with Phyloseq package (McMurdie & Holmes, 2013) to visualize sample clustering. For differential abundance analysis, data were subset into well-watered and drought samples, and low prevalence ASVs (detected in less than 20% of the samples) were removed. Then, Analysis of Compositions of Microbiome with Bias Correction (ANCOM-BC) with structural zeros (ANCOMBC package, Lin & Peddada (2020)) was used to investigate the ASVs significantly enriched and depleted due to exposure to *Ps*VOCs, under drought and well-watered conditions.

### 3. Pseudomonas VOC profiling

#### 3.1 Pseudomonas inoculation and growth

To study volatile profiles emitted by the three bacterial strains, strain suspensions were prepared as described above, with the bacterial titer adjusted to an OD at 600 nm of 0.5. Then, glass Petri dishes with specialized lids equipped with outlets for volatile traps were used, as described by Garbeva et al., (2014). The dishes were filled with 0.1× MS medium prior to inoculation with 150 μl of the strain suspensions, or 150 μl 10 mM MgSO4 for control samples. The dishes were sealed with parafilm and stored for three days in a climate room with a temperature of 20 °C to allow for bacterial growth. Three to five glass Petri dishes were grown per treatment.

#### 3.2 Volatile trapping and analysis

After three days, volatile compounds were collected via passive diffusion using a steel trap containing 200 mg Tenax TA (C1-AAXX-5003, Markes International Ltd, Bridgend, UK) connected to the outlets of the Petri dish lids for 24 h. The volatiles were measured in a GC-QTOF system (7890B GC with 7200AB QTOF, Agilent Technologies Inc, Santa Clara, USA) equipped with an automated thermal desorption unit (Unity TD-100, Markes International, Bridgend, UK). The Tenax traps were heated at 280 °C for 8 min to release the trapped volatiles, which were then further trapped in a cold trap (−10 °C) in the thermal desorption unit (TDU). The cold trap was heated at 300 °C for 5 min. to release volatiles which were transferred with split 1:10 (280 °C transfer line) to the GC-QTOF, equipped with a DBms ultra-inert column (122-5532UI, 30 m length, 0.25 mm internal diameter, 0.25 μm film thickness, Agilent Technologies, Inc., Santa Clara, CA, USA). The GC temperature program was held at 39 °C for 1 min, ramped to 315 °C at 10°C min^−1^, and held at 315 °C for 7 min. A constant Helium (He) flow of 1.2 ml min-1 was used. Then the volatile compounds were ionized in electron ionization (EI) mode at 70 eV. Mass spectra were acquired in full-scan mode (33-400 AMU, 4 scans/sec, 2 GHz Extended Dynamic Range). An alkane mix of C8 till C20 (04070-5 ml, Sigma Aldrich, St. Louis, Missouri, USA) was used to determine the retention index.

The mass spectra of the volatile compounds were visualized using MassHunter Qualitative Analysis software (version 10.0, build 10305.0; Agilent Technologies, Santa Clara, USA) and exported as .mzData XML files for high-throughput data analysis in MZmine (version 3.9). Mass feature tables were filtered to remove silica fragments and features detected in blank samples. The resulting dataset was then subjected to metabolomic analysis using the MetaboAnalyst 5.0 web interface (Pang et al., 2021). Prior to statistical analysis, the data were log-transformed and Pareto scaled. The medium-only control served as a baseline reference for background VOCs emitted by the MS medium. Compounds present exclusively or at significantly higher intensities in bacterial samples were interpreted as bacteria-derived, while control-derived peaks were used to filter out medium-associated VOCs during data processing. Principal Component Analysis (PCA) and hierarchical clustering were subsequently performed to identify group separation and key discriminatory features. PERMANOVA was conducted using the adonis2 package with Euclidean distances. Mass features of interest (key discriminatory features) were tentatively identified by comparing their mass spectra with those in the NIST 2020 and an in-house libraries, using the NIST MS Search program version 2. The tentative identifications were further verified by comparing the Kováts retention indices of the analyzed compounds with those of the reference compounds in the libraries.

## Results

### *Pseudomonas* VOCs increase drought resilience in Arabidopsis

To evaluate the effect of PGPR on drought stress tolerance, three *Pseudomonas* strains were tested for their ability to increase Arabidopsis drought resilience through their volatile blends. Arabidopsis seedlings were grown in half-strength MS (well-watered) or half-strength MS with sorbitol (as a proxy for drought) and co-cultivated with either one of the *Pseudomonas* strains or with a mock solution, without physical contact between bacteria and plants, after which seedling biomass and root architecture were assessed. Under well-watered conditions, *Ps*WCS417 and *Ps*WCS315 VOCs significantly increased seedling biomass compared to control (Fig. 1a, Tables S2-S4). Under drought conditions all three *Pseudomonas* strains significantly increased seedlings biomass (Fig. 1a, Tables S5-S7), with *Ps*WCS417 and *Ps*WCS315 showing a stronger effect. Regarding root architecture, VOCs from all three *Pseudomonas* strains significantly increased primary root length under well-watered conditions (Fig. 1a, Tables S8-S10), but not under drought stress conditions (Fig. 1a, Tables S11-S13). On the other hand, the number of lateral roots was significantly increased by *Ps*VOCs under both well-watered (Fig. 1a, Tables S14-S16) and drought conditions (Fig. 1a, Tables S17-S19), indicating that lateral root proliferation may contribute to enhanced drought resilience. Overall, these results show that VOCs from all three *Pseudomonas* strains enhance seedling biomass and lateral root formation and might, as such, affect Arabidopsis drought tolerance.

Furthermore, we tested the effect of *Ps*WC417 VOCs on Arabidopsis seedlings at different severity of osmotic stress (75, 150 and 250mM of sorbitol). Our data showed that at all treatments, VOCs had a significantly positive effect on plant growth even under very severe stress conditions at 250mM sorbitol (Fig. S2). Since drought and salt stress both induce water and nutrient deficiency in plants, we next tested the effect of *Ps*WC417 VOCs on seedling growth under salt stress conditions. In agreement with the plant growth promoting effect under drought stress, also under salt stress, the VOCs induced plant biomass and lateral root numbers (Fig. S3). The data suggest that *Ps*WC417 VOCs can have a more universal effect and help plants to survive and grow under both drought and salinity stress.

### *Pseudomonas* VOC profiles show similarities but also differences across different strains

To compare VOC composition among the three *Pseudomonas* strains, we measured and identified the VOCs using GC-MS and performed principal component analysis (PCA) on their emitted volatile blends. The PCA and statistical analyses revealed significant separation among strains (Fig.1 b, Fig. S4, Tables S20 and S21), indicating distinct VOC compositions. Further hierarchical clustering of the VOC profiles revealed strain-specific emissions, with several compounds differing in relative abundance among strains (Fig.1c, Fig. S4). Notably, *Ps*WCS417, *Ps*WCS315 and *Ps*CHA0 shared a substantial number of volatile compounds (Fig. S4).. Based on these findings we formulate the hypothesis that VOCs produced by *Ps*WCS417, *Ps*WCS315 and *Ps*CHA0 may play a key role in triggering drought adaptation mechanisms in Arabidopsis. The limited (low-abundance) VOC production in *Ps*WCS315 could be attributed to a slower growth rate of this strain.

### *Pseudomonas* VOCs induce transcriptional changes in Arabidopsis roots

To investigate the molecular mechanisms underlying the enhanced performance of Arabidopsis under drought when exposed to *Ps*VOCs, we performed transcriptomic analyses on the roots of 5 and 6 day-old axenically grown seedlings exposed to drought, and well-watered as control, at two time points: 24 and 48 h of exposure to *Ps*VOCs. Principal component analysis (PCA) (Fig. 2a) revealed a clear separation of samples based on *Pseudomonas* strain, watering regime (well-watered and drought) and time point, which was statistically significant according to the PERMANOVA analysis (Table S22). Transcriptomes of roots exposed to each of the three different *Pseudomonas* strains were significantly different from transcriptomes of roots exposed to mock solution (Table S23). However, the root transcriptomes of plants exposed to *Ps*WCS417 and *Ps*CHA0 were not significantly different from each other (Table S23). This clustering pattern is consistent with our hypothesis, supported by bacterial VOCs profile, which showed that *Ps*WCS315 produced lower amounts of VOCs compared to the other two strains. Differential expression analysis (fold change >1, FDR < 0.05) revealed the number of differentially expressed genes (DEGs) by each strain under each water regime and time point (Fig. 2b, Table S24). To identify candidate genes involved in *Ps*VOCs-mediated drought resilience, we compared the DEGs shared by all three *Pseudomonas* treatments at both time points and under both well-watered and drought conditions. Due to the very low number of DEGs in the *Ps*WCS315 treatment, we focused on the common DEGs affected by *Ps*WCS417 and *Ps*CHA0 VOCs (Fig S5 and S6). At 24 h, we observed 61 DEGs upregulated and 136 DEGs downregulated in roots exposed to *Ps*WCS417 and *Ps*CHA0 VOCs under well water conditions, and 3 DEGs upregulated and downregulated under drought conditions (Fig. S5). However, at 48 h, the number of DEGs was greater under drought conditions, 85 upregulated and 50 downregulated, than under well-watered conditions, 17 upregulated and 18 downregulated. (Fig. S6). Additionally, we found differences in the expression pattern of DEGs across the different *Ps*VOCs treatments and water regimes (Fig. S7). We next performed gene ontology (GO) term analysis of DEGs in *Ps*VOCs-treated roots compared to controls in both well-watered and drought conditions. This analysis revealed enrichment of genes in pathways involved in glucosinolate biosynthesis, response to oxidative stress, L-leucine biosynthesis, and branch-chain amino acid metabolic process, as well as nitrate, sulfur compound and sucrose transmembrane transporter activity (Fig. S8).

Notably, *Ps*VOCs resulted in downregulation of several drought-responsive genes during drought stress (Fig. 2c). For instance, *SNRK2-8*, a gene involved in abscisic acid (ABA) signalling, was downregulated by *Pseudomonas* at both time points under drought and well-watered conditions. Similarly, *LEA7*, an ABA-dependent gene encoding a drought tolerance protein, was downregulated by *Ps*VOCs but only under drought conditions. The *SWEET11* gene coding for a sugar transporter, was downregulated by *Ps*VOCs under drought, compared to mock-treated plants. *JUB1*, a NAC transcription factor known to enhance drought resilience in an ABA independent manner, was highly upregulated under drought conditions, 48 h after exposure to *Ps*VOCs, compared to plants treated with mock control. Similarly, genes involved in iron uptake, *IRT1 and coumarin metabolism, CYP82C4* were upregulated in roots exposed to *Ps*VOCs for 24 h under drought conditions, compared to the drought mock control, while under well-watered conditions this upregulation was milder. Additionally, genes involved in aliphatic glucosinolate biosynthesis, *BCAT4* and *MAM-L*, were upregulated under both drought and well-watered conditions when exposed to *Ps*VOCs compared to their respective mock controls, although the upregulation was greater under drought conditions after 48 h. Interestingly, most of these genes were only differentially expressed upon *Ps*VOCs treatment under drought conditions (Table S25). Taken together, these findings suggest that *Ps*VOCs induce specific and ABA independent drought-related transcriptional changes that likely contribute to the enhanced performance of Arabidopsis seedlings under drought when exposed to a *Pseudomonas* volatile blend.

### ABA and aliphatic glucosinolates affect the response of roots to *Pseudomonas* volatiles

To investigate whether some of the molecular pathways identified by the RNA-seq are required for Arabidopsis-*Pseudomonas* interactions, we selected representative Arabidopsis mutant lines from the ABA signaling pathway (*snrk2.2-2.3*), aliphatic glucosinolate biosynthesis (*bcat-4*) and sugar transport (*sweet11/12*) and exposed them to the *Ps*WCS417 VOCs. Our results showed that *bcat-4* and *snrk2.2/2.3* mutant seedlings increased more in biomass than Col-0 seedlings when exposed to *Ps*WCS417 VOCs in both well-watered and drought conditions (Fig. 3, Fig. S9, Tables S26-S31). Similar trends were observed for primary root length and lateral root count, with a stronger VOC response in *snrk2.2/2.3* in both well-watered and drought conditions, and a stronger VOC response in *bcat-4* only during drought stress (Tables S34-S47). Altogether, these results suggest that both the aliphatic glucosinolate biosynthetic pathway and the ABA pathway may play a negative role in Arabidopsis’ responses to *Ps*VOCs. In contrast, the *sweet11/12* mutant was unaffected by *Ps*WCS417 VOCs in any of the measured parameters (Tables S32-S33, S40-S41 and S48-S49), except for the lateral root count under drought conditions, suggesting that SWEET and ABA signaling genes need to be downregulated for a positive effect of *Ps*VOCs on Arabidopsis.

### Identification of Arabidopsis root metabolic changes during drought and *Ps*VOCs treatment

To confirm the observed gene expression changes in the sugars, aliphatic glucosinolate, and coumarin pathways, we analyzed the levels of these metabolites using HPAEC-PAD Dionex and LC-MS/MS, respectively. Our data showed that from the measured aliphatic glucosinolates (GLS), 4-methylsulfinylbutyl and 5-methylsulfinylpentyl glucosinolates were significantly decreased during the drought treatment in the roots, while 6-methylsulfinylhexyl and 7-methylsulfinylheptyl GLS were decreased only in the combination of drought stress and *Ps*WCS417 VOCs (Fig. S10, Table S51). From the indolic glucosinolates, only 4-methoxy-3-indolylmethyl-glucosinolate was significantly reduced under drought stress (Fig. S11, Table S51).

None of the identified GLS compounds was significantly up-regulated under drought stress or under the combined drought and the *Ps*VOCs treatment compared to the controls. However, for coumarins, we observed accumulation of scopoletin in the roots due to *Ps*WCS417 VOCs treatments, as well as for their precursors: p-coumaric acid and 2-hydroxycinnamic acid (Fig. S11, Table S50). Both compounds were up-regulated in the roots under drought stress. The 2-hydroxycinnamic acid was shown to have antioxidant properties and to reduce the ROS accumulation during water stress (Khawula et al., 2024). In the presence of *Ps*WCS417 VOCs and drought, we observed a decrease of 2-hydroxycinnamic acid, which suggests, in agreement with the better performance of the seedlings, that the plants experience less stress (and ROS accumulation).

To assess the effects of drought and *Ps*VOCs on plant sugar allocation, we measured glucose, fructose, and sucrose concentrations in roots and shoots when exposed to *Ps*WCS417 VOCs. The concentration of the three different sugars was significantly higher in drought conditions compared to well-watered controls in both tissues (Fig. S13, Table S52). For glucose and fructose content, we found a significant effect of the treatment (*Ps*WCS417 VOCs) in roots, with the level of these sugars lower in roots exposed to *Ps*WCS417 VOCs, compared to mock treated roots (Fig. S13, Table S52). Interestingly, we found a significant interaction between water regime and treatment for the concentration of sucrose in roots. While in well-watered conditions, there was no difference in the concentration of sucrose between mock-treated and *Ps*WCS417 VOCs-treated roots under drought conditions, the concentration of sucrose was significantly lower in *Ps*WCS417 VOCs-exposed roots compared to mock-treated roots (Fig. S13, Tables S52-S53).

Altogether, these results show that *Ps*VOCs affect the concentration of specific GLS, coumarins, and sugars in the plant and that some of these *Ps*VOCs induce metabolic changes only occur when plants experience drought stress.

### *Pseudomonas* volatiles enhance the performance of *Brassica oleracea* under well-watered and drought conditions in soil

To investigate whether the enhanced drought resilience from *Ps*VOCs observed in sterile media in the model plant Arabidopsis was maintained in a crop species grown in soil, we grew white cabbage, *Brassica oleracea*, under well-watered and drought conditions in a mix of soil and sand in a two-compartment pot system (Moisan et al., 2020). Via this pot system, the roots were exposed to VOCs produced by one of the three *Pseudomonas* strains or mock (non-inoculated) medium (Fig. 4a). Our results showed that the *Ps*VOCs from the three different strains significantly increased the shoot biomass of *B. oleracea* grown in soil, both in well-watered (up to 100% increase) and in drought conditions (Fig. 4c, left, Tables S54-S56). In addition, the water content of plants exposed to *Ps*VOCs of the three different strains was also significantly higher in both well-watered and drought conditions than those exposed to control media (Fig. 4c, right, Tables S57-S59). In summary, our results demonstrate that *Ps*VOCs of the three *Pseudomonas* strains used can promote the growth of *B. oleracea* both under well-water and drought conditions in soil.

**Figure 4.**
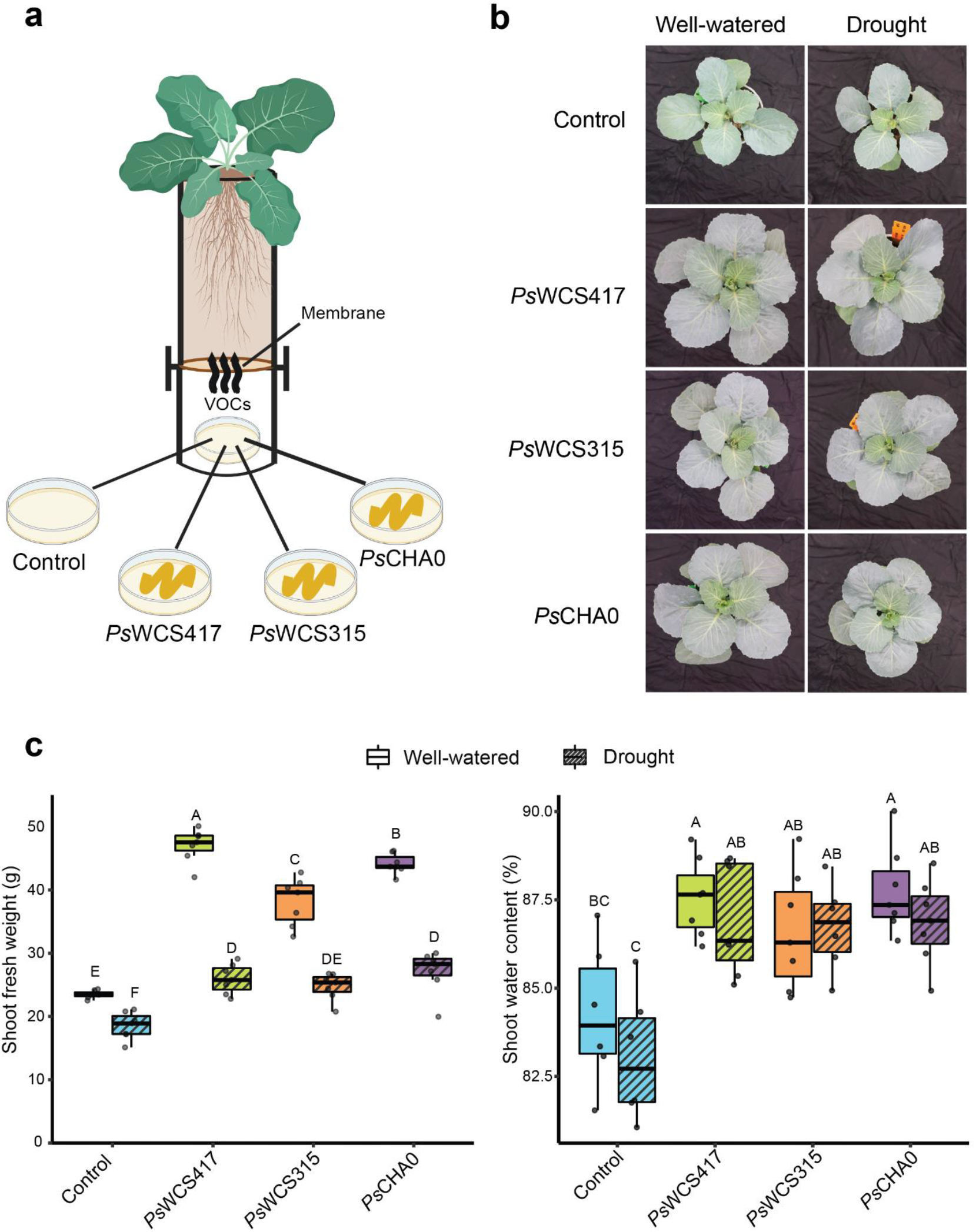
*In vivo* effects of *Ps*WCS417, *Ps*WCS315 and *Ps*CHA0 VOCs on *B. oleracea* plants grown in soil under well-watered and drought conditions. (a) Schematic representation of the two-compartment pot system (Moisan et al., 2020) to expose roots of *B. oleracea*, growing in the top compartment, to volatiles emitted by one of the *Pseudomonas* strains growing in a Petri dish in the bottom compartment or by a control Petri dish containing only media. (b) Representative pictures of *B. oleracea* plants exposed to the different VOC treatments on the day of the harvest (Control = mock, *Ps*WCS417 = *Pseudomonas simiae* WCS417, *Ps*WCS315 = *P. simiae* WCS315, *Ps*CHA0 = *P. protegens* CHA0). (c) Shoot fresh weight (left) and percentage of shoot water content (right) (b) shoot fresh weight and (c) percentage of water content (Wilcoxon test, p< 0.05, n= 7, different letters indicate significant differences, boxplots display median, interquartile range, and data distribution.

### *Pseudomonas* volatiles modify the root microbiome of *Brassica oleracea* under drought

To investigate whether *Ps*VOCs influence not only *B. oleracea* growth but also shape its root microbiome composition, we performed 16S amplicon sequencing on soil samples collected from the two-compartment pot system (Fig. 4a). Non-metric multidimensional scaling (NMDS) analysis revealed a distinct separation between the root microbiomes of plants grown under well-watered and drought conditions (Fig. 5a). Additionally, clustering of samples was also observed according to the volatile emitting *Pseudomonas* strain. PERMANOVA further confirmed that both the water regime and the *Pseudomonas* strain VOCs had significant effects on the microbial community (Table S60). However, alpha diversity was not significantly influenced by either the water regime or the *Pseudomonas* treatment, except for Simpsons diversity of plants grown under the control water regime (Figure S14, Tables S61-S65). Since VOCs from all three *Pseudomonas* strains enhanced *B. oleracea* growth (Fig. 4c), we focused on the ASVs that were significantly enriched or depleted by exposure to VOCs of the three strains (Fig. 5b). Unexpectedly, most of the enriched ASVs were common among the three strains, whereas only a small number of the depleted ASVs were common across treatments, both for well-watered and drought conditions. In both well-watered and drought conditions, the majority of commonly enriched ASVs by the three *Pseudomonas* strains belonged to the phylum Proteobacteria, followed by Planctomycetota and Actinobacteriota (Fig. 5c, Fig. S15, Table S66). Our results demonstrate that *Ps*VOCs can reshape the root microbiome of *B. oleracea* and suggest that, beyond modifying the plant transcriptome, *Ps*VOCs may enhance plant growth and drought resilience by directly or indirectly (by changing the root exudates) recruiting beneficial bacteria to the roots.

**Figure 5.**
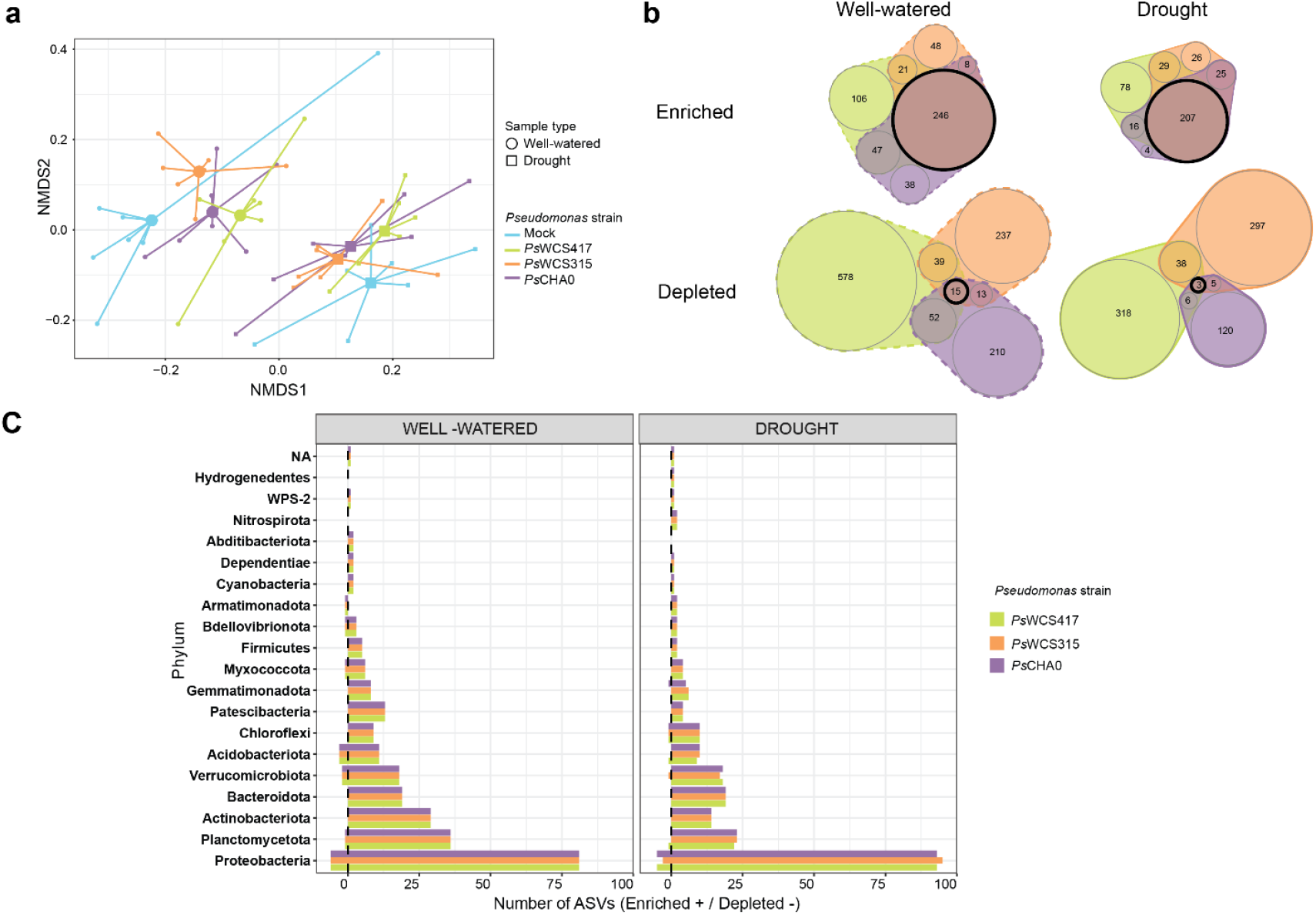
Impact of *Pseudomonas* VOCs on the *B. oleracea* root microbiome under well-watered and drought conditions. (a) Non-metric multidimensional scaling (NMDS) showing differences in bacterial community composition between different water regimes and treatments with *Pseudomonas* strains VOCs. Shapes represent well-watered (circles) and drought (squares) regimens, while colours indicate the different *Pseudomonas* strains (blue = mock, green = *Ps*WCS417, orange = *Ps*WCS315 and purple = *Ps*CHA0, n= 7). (b) Venn diagrams displaying the number of ASVs enriched (top) or depleted (bottom) in response to *Ps*VOCs under well-watered (dashed line) and drought (continuous line) conditions, which were obtained from the differential abundance test performed with ANCOM-BC test. Colours indicate the different *Pseudomonas* strains (green = *Ps*WCS417, orange = *Ps*WCS315 and purple = *Ps*CHA0). The size of each circle represents the number of ASVs, with overlapping areas indicating shared ASVs among *Pseudomonas* treatments. Black circle indicates the ASVs commonly enriched/depleted by the three *Pseudomonas* strains. (c) Number of ASVs enriched or depleted by the three different *Pseudomonas* strains under well-watered (top) and drought (bottom) conditions. X-axis shows number of ASVs and Y-axis the bacterial phylum. Each bar represents an individual ASV, color-coded according to the *Pseudomonas* strains used for the treatments.

## Discussion

Plants live in a close relationship with soil microorganisms, which contribute to plant health by increasing their growth and resilience towards (a)biotic stresses (Porter et al., 2020; Rubin et al., 2017). Soil microorganisms can produce a wide range of secondary metabolites that can directly or indirectly protect plants from stresses, for example by modulating plant hormonal pathways or inhibiting the proliferation of pathogenic microorganisms (Li et al., 2021; Moisan et al., 2019, 2020; Pascale et al., 2020). Some of the microbial secondary metabolites are non-volatile, while others are volatile. Bacterial VOCs act as signaling molecules capable of travelling short and long distances through the soil and are known to play important roles in both intra- and inter-kingdom interactions (Schulz-Bohm et al., 2017; Weisskopf et al., 2021). However, our understanding of microbial volatiles in the rhizosphere and their role in induced plant resilience to (a)biotic stresses is still limited (Garbeva & Weisskopf, 2020). While previous work has extensively studied the plant-growth promoting effect of *Ps*WCS417 and its role in conferring systemic induced resistance (Pieterse et al., 2021), its ability to increase plant resilience to abiotic stress is not well explored (Aparicio et al., 2025; Chiappero et al., 2019). Additionally, most of the studies focused on the effect of the direct inoculation of *Ps*WCS417 on plant roots, and less attention has been paid to the effects of its VOCs (Desrut et al., 2020; Zamioudis et al., 2013). In this study we show that VOCs of the *Pseudomonas simiae* WCS417 and two other strains, *P. simiae* WCS315 and *P. protegens* CHA0, (1) increase plant performance under drought in sterile media in *Arabidopsis thaliana* and in soil in *Brassica oleracea* and (2) modify the root microbiome composition to resemble that of a well-watered plant, and (3) the underlying molecular mechanism is associated with the ABA pathway and aliphatic GSL biosynthesis.

Drought stress causes a series of physiological changes in plants, including an increase in the plant hormone ABA, which is an important regulator of stomatal opening and plant growth and resilience to drought in general (Mo et al., 2024). Interestingly, our data show that without any physical contact between the rhizobacteria and roots, bacterial VOCs, from *Ps*WCS417 and *Ps*CHA0, triggered transcriptional changes in Arabidopsis roots associated with suppression of ABA biosynthesis and signaling. For example, *SnRK2.8*, which phosphorylates downstream targets of the ABA pathway leading to stomatal closure and growth inhibition, and *LEA7* (Kumar et al., 2019; Ng et al., 2014; Waadt et al., 2015), were both downregulated. Similarly, *AT5G28520* (encoding a jacalin lectin) (Jia & Rock, 2013) and *TSPO*, which regulates aquaporin expression (Hachez et al., 2014), were specifically downregulated by *Ps*VOCs under drought conditions. Since ABA is generally associated with improved drought resilience, the observation that *Ps*VOCs downregulate ABA-related responses yet still promote plant growth under drought conditions appears counterintuitive. We hypothesize that *Ps*VOCs may activate ABA-independent stress response pathways, like JUB1, thereby enabling the suppression of ABA signaling, which could be the cause of the growth promotion effect. By modulating these pathways, *Ps*VOCs may help the plant bypass ABA-induced growth arrest and stomatal closure, allowing growth even under water-limited conditions. Moreover, *snrk2.2/2.3* (an ABA insensitive mutant) plants exhibited increased growth when exposed to *Ps*WCS417 VOCs, compared to Col-0 plants. Interestingly, ABA-activated *SnRK2*s phosphorylate *SWEET*s in the shoot to enhance root sucrose allocation and drought resilience (Chen et al., 2022). Thus, the downregulation of *SWEET*s observed in response to exposure to *Ps*VOCs, as well as the specific decrease in sucrose on Arabidopsis roots exposed to *Ps*VOCs under drought, may reflect an ABA-independent mechanism for promoting root growth. Since *SWEET* genes also regulate sucrose rhizodeposition and microbiome assembly (Hennion et al., 2019; Loo et al., 2024), their repression may limit exudation in the soil, potentially reducing microbial competition in the rhizosphere. Complementarily, *RS4*, a raffinose synthase gene, was upregulated under drought conditions. Raffinose, synthesized from sucrose, functions as an osmoprotectant and ROS scavenger (Li et al., 2020; Pantigoso et al., 2025), suggesting that *Ps*VOCs also enhance drought tolerance through osmolyte accumulation. These data support our hypothesis (Figure 6) that *Ps*VOCs enhance plant growth under drought stress by suppressing ABA and suggests a negative interaction between *Ps*VOCs and ABA signaling in plants. Other transcriptional changes also involved key pathways related to coumarin and iron metabolism, sugar transport, aliphatic GSL biosynthesis and lateral root development. Furthermore, genes associated with regulation of lateral root development and increased shoot biomass were differentially regulated after exposure to *Ps*VOCS.

**Figure 6.**
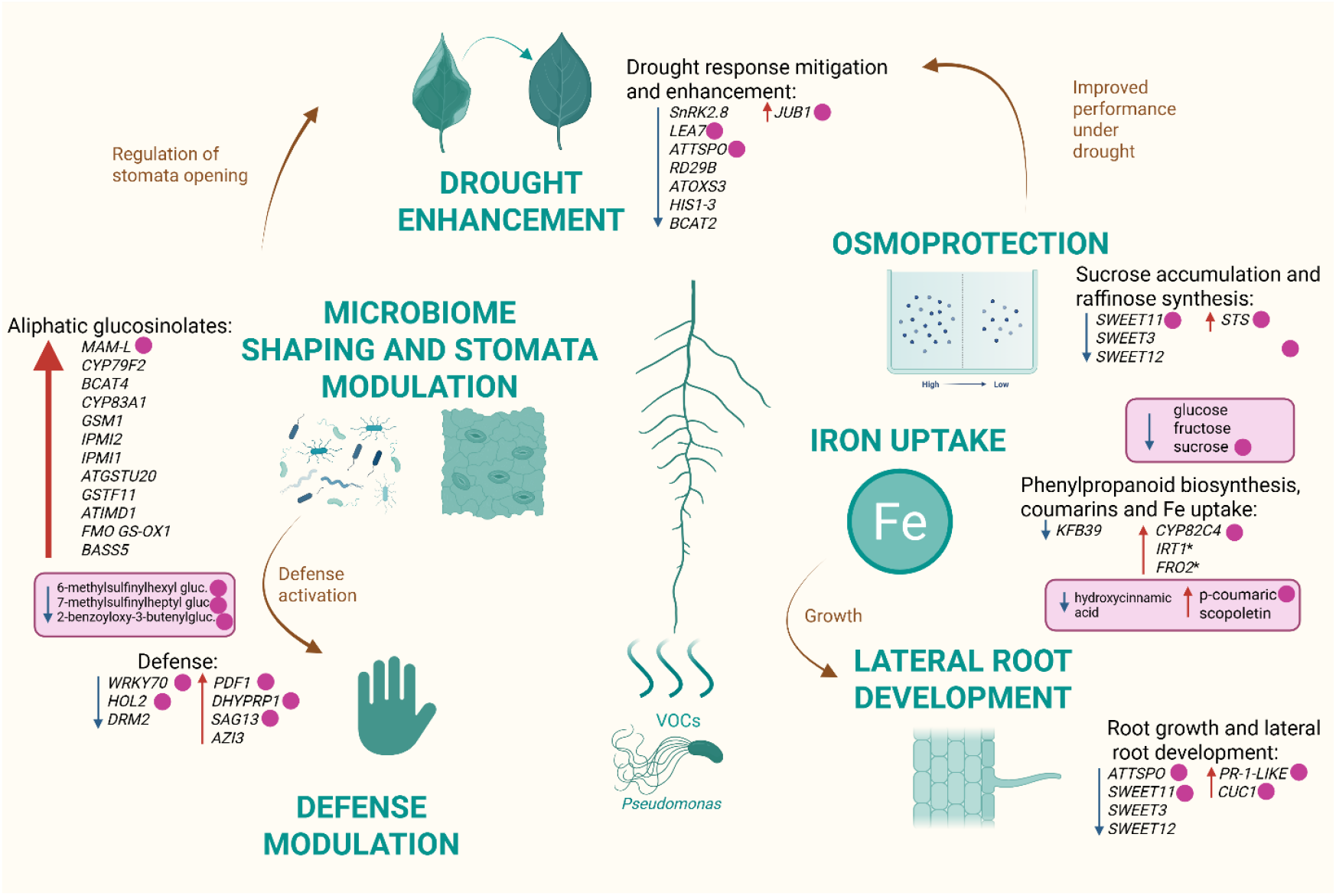
Model of interaction of *Ps*VOCs with plants. Summary of the molecular pathways affected in the plant upon exposure to Pseudomonas volatiles. Soil bacteria, such as Pseudomonas species, emit signalling molecules, including volatile organic compounds (VOCs). Bacterial VOCs can travel short and long distances through the soil and are perceived by plant roots, leading to significant changes in the rhizosphere microbiome and in root gene expression and growth under drought stress. PsVOCS induced the expression of JUB1, a master regulator of drought tolerance in plants, and downregulated the SnRK and SWEET11 transporters, leading to reduced sucrose in the roots. These changes in gene expression also affect the production of plant secondary metabolites, such as coumarins (important for Fe uptake under drought stress) and aliphatic glucosinolates, which are then exuded and alter the plants’ rhizosphere microbiome. The changes in the rhizosphere microbiome also led to phenotypic changes in plants and alterations in root architecture. Overall, these plant-microbiome interactions mediated by PsVOCs contribute to improved plant growth and survival under drought stress. Arrows indicate upregulation (red) or downregulation (blue) of genes. Pink circles indicate genes that were only affected under drought conditions (Padj < 0.05 and |Log2FC| > ±2). * indicate genes which Log2FC was below ±2 but show a consistent pattern. Pink boxes indicate changes in metabolite levels, where pink circles also indicate those compounds that were affected by *Ps*VOCs only in drought conditions.

As summarized in Figure 6, we hypothesize that *Ps*VOCs also suppress branched-chain amino acid (BCAA, *BCAT2*) levels. Opposite to proline, BCAA accumulation typically results from ABA-dependent protein degradation under drought, and *BCAT2* was shown to function in restoring the basal BCAA levels once the drought stress is alleviated (Huang & Jander, 2017). These data further suggest that *Ps*VOC-exposed plants perceived reduced drought stress, which allows them to maintain growth similar to plants grown in well-watered conditions. The transcription factor *JUB1* was upregulated upon *Ps*VOCs exposure under drought conditions. *JUB1* enhances drought tolerance by increasing the proline content, reducing ROS, and maintaining water content (Samantaray et al., 2025; Thirumalaikumar et al., 2018), further reinforcing the increased growth under drought observed in our experiments. The enhanced plant growth can also be due to changes in the root architecture and growth, which will lead to higher nutrient and water uptake. Genes such as *PR-1-LIKE* and *CUC1*, which are involved in auxin dependent lateral root development (Hu et al., 2024; Zhang et al., 2020; He et al., 2005) were upregulated under drought, linking the *Ps*VOCs function to morphological changes in the plant root system. Interestingly, beyond growth-related responses, *Ps*VOCs also influenced plant immunity. For example, genes associated with systemic acquired resistance (SAR), such as *PDF1* and *DHYPRP1*, were induced, likely through JA-mediated pathways known to antagonize ABA signaling (Yasuda et al., 2008). *SAG13*, a defense gene against bacterial and fungal pathogens (Dhar et al., 2020), was also upregulated, suggesting a broader immune activation. Interestingly, genes *HOL2* and *SnRK2.8*, shown to be involved in susceptibility to *Pseudomonas*, were downregulated. *HOL2* contributes to plant glucosinolate metabolism and pathogen resistance, and its mutation enhances susceptibility to *P. syringae* (Nagatoshi & Nakamura, 2009). Moreover, *SnRK2.8* phosphorylates the *P. syringae* effector *AvrPtoB*, promoting bacterial virulence (Lei et al., 2020). These data suggest that *Ps*VOCs may selectively suppress host defenses to facilitate beneficial *Pseudomonas* colonization while maintaining broader pathogen resistance.

In agreement with previous findings on *Ps*WCS417 inoculation (Stringlis et al., 2018; Zamioudis et al., 2015), here we show that *Ps*VOCs can also upregulate genes involved in phenylpropanoid biosynthesis and iron uptake. For instance, *KFB39*, a repressor of *PAL (*Phenylalanine ammonia-lyase) (X. Zhang et al., 2015), was downregulated. Meanwhile, *CYP82C4, FRO2* and *IRT1*, key genes in coumarin metabolism and iron mobilization, were upregulated under drought. Their coordinated expression under *Ps*VOC treatment aligns with enhanced root iron acquisition. Notably, *CYP82C4* converts fraxetin to sideretin, which is secreted to the rhizosphere under iron deficiency to facilitate iron uptake by *IRT1* and *FRO2* (Stassen et al., 2021; Rajniak et al., 2018). Consistent with the transcriptomics data, our metabolomics analysis also showed upregulation of scopoletin as well as p-coumaric acids and 2-hydroxicinamic acid by *Ps*VOCs treatment. Interestingly, p-coumaric acid was shown to be involved in the recruitment of bacteria at the phylo and rhizosphere as well as inducing siderophore biosynthesis in the algal bacterial symbiont *Phaeobacter inhibens* (Chen et al., 2022; Su et al., 2024; Wang et al., 2022). The relationship between iron homeostasis, drought stress and the root microbiome was recently described for sorghum (Xu et al., 2021). The authors hypothesized that under drought stress, the plant root microbiome is enriched in bacteria that can accumulate iron from the soil, as iron availability decreases for both plant roots and rhizosphere microbiota during water deprivation. In agreement with this, we show that *Ps*VOC-induced activation of iron uptake pathways may prepare roots to better compete for iron under drought conditions, influencing both plant health and microbial interactions. Other metabolites shown to be important for the rhizosphere microbial composition (Bressan et al., 2009; Chroston et al., 2024) and drought resilience in plants (AbdElgawad et al., 2023; Martínez-Ballesta et al., 2015) are aliphatic GLS. Interestingly, genes involved in aliphatic GLS biosynthesis were also strongly upregulated by *Ps*VOCs, especially under drought, suggesting that *Ps*VOCs modulate chemical signaling and plant-microbiome interactions with effects on microbiome assembly. However, in contrast to coumarins and transcriptomics data, we observed a reduction of GLS (aliphatic and indolic) under drought or combination of drought and VOCs treatment, supporting the hypothesis of GLS being exuded and having a role in root microbiome assembly, as it was shown for the phyllosphere (Unger et al., 2024). Furthermore, our two-compartment pot system showed that *Ps*VOCs can shape and modulate the rhizosphere composition under both well-watered and drought soil conditions. Interestingly, Proteobacteria was the phylum with the most enriched number of ASVs in *B. oleracea* roots exposed to *Ps*VOCs, even under drought stress. This is surprising, as previous studies have shown that drought stress typically leads to a depletion of Proteobacteria and an enrichment of Actinobacteria in the root microbiome (Fitzpatrick et al., 2018; Naylor et al., 2017; Santos-Medellín et al., 2021; Berdaguer et al., 2024). Our results suggest that *Ps*VOC exposure maintains a root microbiome composition more characteristic of healthy, unstressed plants. This shift could be due to the direct communication between *Pseudomonas* and the native soil microbial community via VOCs (Yuan et al., 2017), or indirectly, through changes in *B. oleracea* root exudates induced by transcriptional responses to *Ps*VOCs (Kong et al., 2021). Consistent with our RNA-seq data, Xu et al., (2021) reported that drought stress leads to the downregulation of plant iron uptake genes. In their study, this downregulation was necessary for the enrichment of Actinobacteria in the rhizosphere under drought conditions, as demonstrated using iron uptake-deficient maize mutants. Therefore, the upregulation of iron uptake genes observed in Arabidopsis roots exposed to *Ps*VOCs under drought may explain the lack of Actinobacteria enrichment in *Ps*VOC-treated *B. oleracea* plants under drought. Altogether, these findings highlight the critical role of bacterial VOCs in shaping rhizosphere microbial communities that support plant resilience under drought stress.

In addition, we used GC-MS to analyse and compare the volatile blends of the three *Pseudomonas* strains used in this study. Our data show that *Ps*WCS417 and *Ps*CHA0 emit very similar blends of VOCs, some of which are known to induce plant-growth promotion. For example, dimethyl disulfide – a common bacterial VOC - promotes plant growth by enhancing sulfur nutrition (Meldau et al., 2013). Hexanoic acid enhances pathogen resistance in tomato by priming both the JA and SA pathways (Anderson & Kim, 2021; Aranega-Bou et al., 2014; Scalschi et al., 2013). Acetic acid has been described as a biostimulant that enhances drought resilience in *Arabidopsis*, rapeseed, maize, rice, and wheat by stimulating the JA signalling pathway (Kim et al., 2017). Furthermore, a recent study showed that acetic acid induces drought tolerance in apple trees by modulating the ABA- and JA-induced MAPK signalling pathways (Sun et al., 2022). Taken together, these results suggest that the volatile blends of *Ps*WCS417 and *Ps*CHA0 have the potential to enhance the growth and drought tolerance of the plants, as supported by our transcriptomic analysis. Therefore, our data suggest a “core PGP VOCs blend” that can be produced by different bacteria and has the same function on plants.

Our two-compartment pot system experiment demonstrated that *Ps*VOCs can enhance growth, water uptake and retention both under well-watered and drought conditions in crops grown in soil. Moreover, amplicon analysis revealed that *Ps*VOCs can alter the bacterial composition of the soil attached to *B. oleracea* roots, consistent with previous reports on other bacterial strains (Yuan et al., 2017). Our data showed that bacterial VOCs can act as signaling molecules in plants and can be used in microbial microbe-assisted breeding strategies to enhance crop biomass and drought resilience.

Overall, our work shows that microbial volatiles play an important role in the modification of root architecture in response to drought stress and in re-shaping rhizosphere communities. This offers new avenues for in-depth studies on below-ground perception of VOCs by plants, and identification of chemical communication between microbes and microbes and/or plants that can be used to develop new approaches for drought stress resilience in crops. Our data also confirm that the plant and its microbiota form a holobiont, and that we should embrace the complexity of their interactions when studying drought responses in soil grown plants.

## Author contributions

ZCL, KJK, MD, CT and RK design the research; ZC, MSR, RB, CBL, CSA and JW performed the research and collected the data; ZC, MSR, KJK, MD, CT, PG and RK analyzed and interpreted the data; ZCL wrote the initial version of the manuscript and all authors contributed to its revision.

## Acknowledgements

We thank Hans Zweers for running the GC-MS. We thank Eveline Bosman, Ariadna Mendoza Mederos, Franco Sanchez Izurieta, [Francel Verstappen](https://www.sciencedirect.com/author/7004194910/francel-w-a-verstappen) (Plant physiology, WUR) and Annemarie Dechesne (Plant Breeding, WUR) for their contribution to this project. This work was supported by the Dutch Research Council (NWO/OCW), as part of the MiCRop Consortium Programme, Harnessing the second genome of plants (grant number 024.004.014)

## Data availability statement

Raw sequence data can be found under NCBI Bioproject PRJNA1289803 for 16S sequencing of the Brassica oleracea root microbiome and RNA-seq sequencing of Arabidopsis roots. All scripts for data analysis are available in https://github.com/zulemacl/Pseudomonas_VOCs.

