## Supplementary material for "*Pseudomonas* volatiles shape the root transcriptome and microbiome to promote plant growth under drought": SI

***New Phytologist* Supporting Information**

Article acceptance date: Click here to enter a date.

The following Supporting Information is available for this article:

**Fig. S1** Schematic representation of the different experimental setups used for Arabidopsis growth in sterile conditions, depending on the goal of the experiment.

**Fig. S2** Effect of *Ps*WCS417 VOCs on Arabidopsis osmotic stress tolerance in sterile media.

**Fig. S3** Effect of *Ps* WCS417 VOCs on Arabidopsis salt stress tolerance in sterile media.

**Fig. S4** Heatmap of VOCs emitted by *Pseudomonas* strains.

**Fig. S5** Intersection of differentially expressed genes across treatments and water regimes at 24 h visualized using an UpSet plot.

**Fig. S6** Intersection of differentially expressed genes across treatments and water regimes at 48 h visualized using an UpSet plot.

**Fig. S7** Heatmap of log2 fold change (L2FC) for differentially expressed genes across *Ps*WCS417 *and Ps*CHA0 treatments and water regime.

**Fig. S8** Gene ontology (GO) enrichment of differentially expressed genes.

**Fig. S9** Effect of *Ps*WCS417 VOCs Col-0, bcat-4, snrk2.2/2.3 and sweet11/12 Arabidopsis accessions grown in sterile media.

**Fig. S10** Effect of *Ps*WCS417 VOCs on aliphatic glucosinolate levels in roots of axenically grown Arabidopsis under well-watered and drought conditions.

**Fig. S11** Effect of *Ps*WCS417 VOCs on indolic glucosinolate levels in roots of axenically grown Arabidopsis under well-watered and drought conditions.

**Fig. S12** Effect of *Ps*WCS417 VOCs on coumarin levels in roots of axenically grown Arabidopsis under well-watered and drought conditions.

**Fig. S13.** Effect of *Ps*WCS417 VOCs on plant sugar content.

**Fig. S14** Alpha diversity analysis.

**Fig. S15** Impact of *Pseudomonas* VOCs on the *B. oleracea* root microbiome under well-watered and drought conditions.

**Table S1** Primers used for the genotyping of the Arabidopsis mutant lines.

**Table S2** Type III Analysis of Deviance (Wald Chi-Square Test) for Seedling Biomass.

**Table S3** Pairwise Comparisons of Treatments Using Tukey's HSD.

**Table S4** Compact Letter Display (CLD) of Estimated Means Across Treatments.

**Table S5** Type III Analysis of Deviance (Wald Chi-Square Test) for Seedling Biomass under drought conditions.

**Table S6** Pairwise Comparisons of Treatments Using Tukey's HSD.
**Table S7** Compact Letter Display (CLD) of Estimated Means Across Treatments.
**Table S8** Type III Analysis of Deviance (Wald Chi-Square Test) for Seedling Primary Root length under well-watered conditions.
**Table S9** Pairwise Comparisons of Treatments Using Tukey's HSD.
**Table S10** Compact Letter Display (CLD) of Estimated Means Across Treatments.
**Table S11** Type III Analysis of Deviance (Wald Chi-Square Test) for Seedling Primary Root length under drought conditions.
**Table S12** Pairwise Comparisons of Treatments Using Tukey's HSD.
**Table S13** Compact Letter Display (CLD) of Estimated Means Across Treatments.

**Table S14** Type III Analysis of Deviance (Wald Chi-Square Test) for Seedling Lateral Root Count under well-watered conditions.
**Table S15** Pairwise Comparisons of Treatments Using Tukey's HSD.
**Table S16** Compact Letter Display (CLD) of Estimated Means Across Treatments.
**Table S17** Type III Analysis of Deviance (Wald Chi-Square Test) for Seedling Lateral Root Count under drought conditions.
**Table S18** Pairwise Comparisons of Treatments Using Tukey's HSD.
**Table S19** Compact Letter Display (CLD) of Estimated Means Across Treatments.
**Table S20** PERMANOVA Results (Bray-Curtis Dissimilarity) on volatile blends of *Pseudomonas* strains and mock.
**Table S21** Pairwise PERMANOVA Results on volatile blends of *Pseudomonas* strains and mock.
**Table S22.** PERMANOVA Results Based on PCA Scores.
**Table S23** Pairwise PERMANOVA results based on Euclidean distances of PCA scores from RNA-seq data in Arabidopsis roots.
**Table S24**. List of genes retrieved from DESEq2 of roots of Arabidopsis thaliana exposed to VOCs of PsWCS417, PsCHA0 and PsWCS315 for 24 and 48 h.
**Table S25** Gene Ontology (GO) annotations for genes uniquely differentially expressed in seedlings exposed to *Ps*WCS417 and *Ps*CHA0 under drought conditions.
**Table S26** Type II Analysis of Deviance (Wald Chi-Square Test) for Seedling Biomass in Col-0 seedlings grown in control conditions.
**Table S27** Type II Analysis of Deviance (Wald Chi-Square Test) for Seedling Biomass in Col-0 seedlings grown in drought conditions.
**Table S28** Type II Analysis of Deviance (Wald Chi-Square Test) for Seedling Biomass in *bcat4* seedlings grown in control conditions.
**Table S29** Type II Analysis of Deviance (Wald Chi-Square Test) for Seedling Biomass in *bcat4* seedlings grown in drought conditions.
**Table S30** Type II Analysis of Deviance (Wald Chi-Square Test) for Seedling Biomass in *snrk2.2/2.3* seedlings grown in control conditions**.**
**Table S31** Type II Analysis of Deviance (Wald Chi-Square Test) for Seedling Biomass in *snrk2.2/2.3* seedlings grown in drought conditions.

**Table S32** Type II Analysis of Deviance (Wald Chi-Square Test) for Seedling Biomass in *sweet11/12* seedlings grown in control conditions.

**Table S33** Type II Analysis of Deviance (Wald Chi-Square Test) for Seedling Biomass in *sweet11/12* seedlings grown in drought conditions.

**Table S34** Type II Analysis of Deviance (Wald Chi-Square Test) for Primary Root Length in Col-0 seedlings grown in control conditions.

**Table S35** Type II Analysis of Deviance (Wald Chi-Square Test) for Primary Root Length in Col-0 seedlings grown in drought conditions.

**Table S36** Type II Analysis of Deviance (Wald Chi-Square Test) for Primary Root Length in *bcat4* seedlings grown in control conditions.

**Table S37** Type II Analysis of Deviance (Wald Chi-Square Test) for Primary Root Length in *bcat4* seedlings grown in drought conditions.

**Table S38** Type II Analysis of Deviance (Wald Chi-Square Test) for Primary Root Length in *snrk2.2/2.3* seedlings grown in control conditions.

**Table S39** Type II Analysis of Deviance (Wald Chi-Square Test) for Primary Root Length in *snrk2.2/2.3* seedlings grown in drought conditions.

**Table S40** Type II Analysis of Deviance (Wald Chi-Square Test) for Primary Root Length in *sweet11/12* seedlings grown in control conditions.

**Table S41** Type II Analysis of Deviance (Wald Chi-Square Test) for Primary Root Length in *sweet11/12* seedlings grown in drought conditions.

**Table S42** Type II Analysis of Deviance (Wald Chi-Square Test) for Lateral Root Count in Col-0 seedlings grown in control conditions.

**Table S43** Type II Analysis of Deviance (Wald Chi-Square Test) for Lateral Root Count in Col-0 seedlings grown in drought conditions**.**

**Table S44** Type II Analysis of Deviance (Wald Chi-Square Test) for Lateral Root Count in *bcat4* seedlings grown in control conditions.

**Table S45** Type II Analysis of Deviance (Wald Chi-Square Test) for Lateral Root Count in *bcat4* seedlings grown in drought conditions.

**Table S46** Type II Analysis of Deviance (Wald Chi-Square Test) for Lateral Root Count in *snrk2.2/2.3* seedlings grown in control conditions.

**Table S47** Type II Analysis of Deviance (Wald Chi-Square Test) for Lateral Root Count in *snrk2.2/2.3* seedlings grown in drought conditions.

**Table S48** Type II Analysis of Deviance (Wald Chi-Square Test) for Lateral Root Count in *sweet11/12* seedlings grown in control conditions.

**Table S49** Type II Analysis of Deviance (Wald Chi-Square Test) for Lateral Root Count in *sweet11/12* seedlings grown in drought conditions.

**Table S50** Statistical analysis of coumarins detected by LC-MS.

**Table S51** Statistical analysis of glucosinolates detected by LC-MS.

**Table S52** ANOVA results for sugar concentrations of axenically grown Arabidopsis seedlings.

**Table S53** Tukey HSD post-hoc pairwise comparison showing pairwise comparisons for sucrose concentrations

**Table S54** Kruskal-Wallis Test Result on Shoot Fresh Biomass of B. oleracea.

**Table S55** Pairwise Wilcoxon Test (BH Adjusted) Result on Shoot Fresh Biomass of B. oleracea.
**Table S56** Compact Letter Display (CLD) on Shoot Fresh Biomass of *B. oleracea*.
**Table S57** Kruskal-Wallis Test Result on B. oleracea Shoot Water Content.
**Table S58** Pairwise Wilcoxon Test (BH Adjusted) on B. oleracea Shoot Water Content.
**Table S59** Compact Letter Display (CLD) on B. oleracea Shoot Water Content.
**Table S60** PERMANOVA Results on root microbiome.
**Table S61** ANOVA Results for Shannon Diversity on root microbiome.
**Table S62** Kruskal-Wallis Test Results Simpson Diversity on root microbiome.
**Table S63** Dunn's Post-Hoc Test Results
**Table S64** ANOVA Results for Simpson Diversity
**Table S65** ANOVA Results for Observed Diversity
**Table S66** List of ASVs retrieved from ANCOM-BC differential abundance analyses performed on *Brassica oleracea* root microbiome 16S amplicon sequencing data.

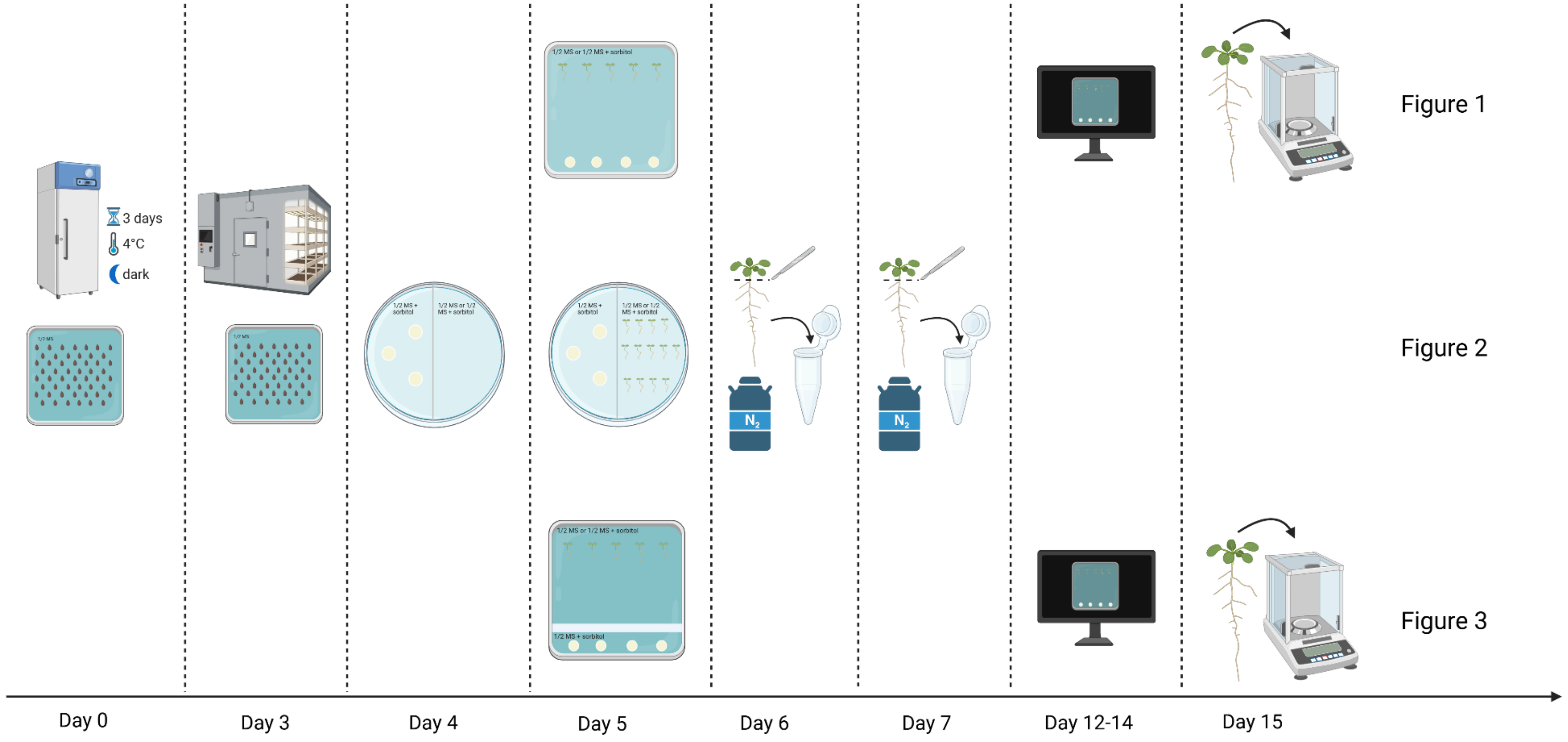

3 days
4°C
dark
1/2 MS or 1/2 MS + sorbitol
1/2 MS + sorbitol
1/2 MS or 1/2 MS + sorbitol
$N_2$
Figure 1
Figure 2
Figure 3
Day 0
Day 3
Day 4
Day 5
Day 6
Day 7
Day 12-14
Day 15

**Fig. S1** Schematic representation of the different experimental setups used for Arabidopsis growth in sterile conditions, depending on the goal of the experiment. Schematic representation of the different experimental setups used for *Arabidopsis* growth in sterile conditions, depending on the goal of the experiment. Root architecture phenotyping experiments were conducted using square plates. In experiments where Arabidopsis Col-0 was co-cultivated with three different *Pseudomonas* strains (Fig. 1A), as well as in the experiments were we tested different sorbitol concentrations and salt stress, plants and bacteria were grown in the same medium. For the experiment investigating the effects of *Pseudomonas simiae* WCS417 volatile organic compounds (VOCs) on various Arabidopsis mutants (Fig. 3), a 0.5 cm strip of agar was removed from the medium to ensure physical separation between plants and bacteria. In this setup, bacteria were consistently grown in half-strength Murashige and Skoog (½ MS) medium supplemented with 150 mM sorbitol. For the RNA-seq experiment (Fig. 2), two-compartment Petri dishes (100 mm diameter) with a central partition were used to separate plant and bacterial growth areas.

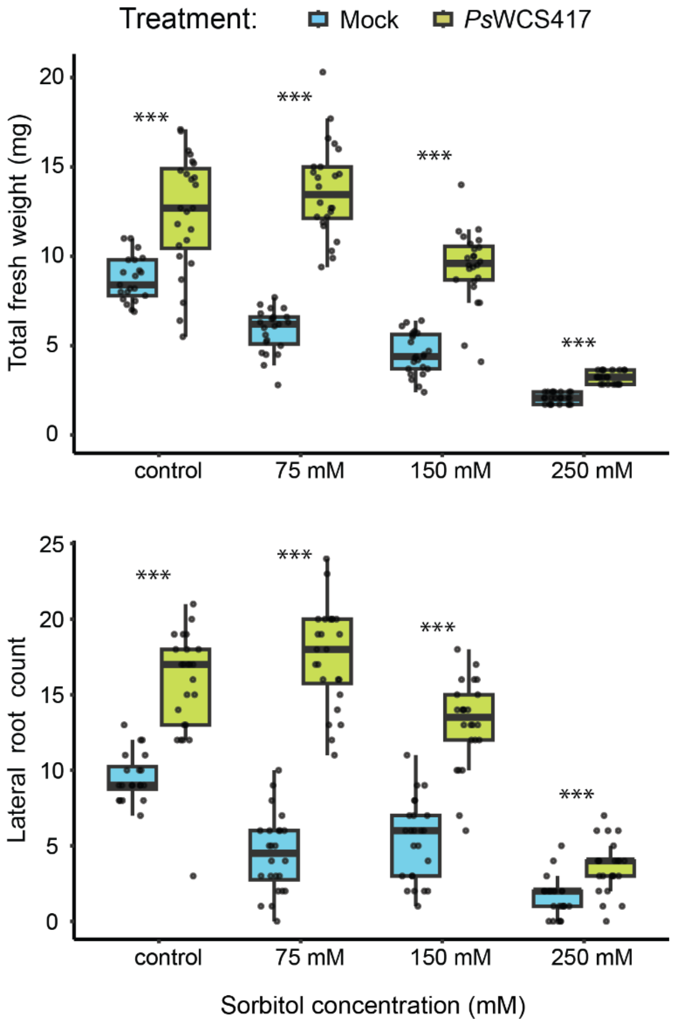


**Fig. S2** Effect of *Ps*WCS417 VOCs on Arabidopsis osmotic stress tolerance in sterile media. Seedling fresh weight (top) and lateral root count (bottom) of Arabidopsis seedlings co-cultivated with *Ps*WCS417 or mock under different sorbitol concentrations (0 (control), 75, 150 and 250 mM). Boxplots show the median, interquartile range, and data distribution. Data was tested for normality and analyzed using a Student's t-test (normal distribution) or a Wilcoxon rank-sum test (non-normal distribution). Asterisks indicate statistically significant differences among treatments ( $P < 0.05$ *, $P < 0.01$ **, $P < 0.001$***) (n= 20-25).

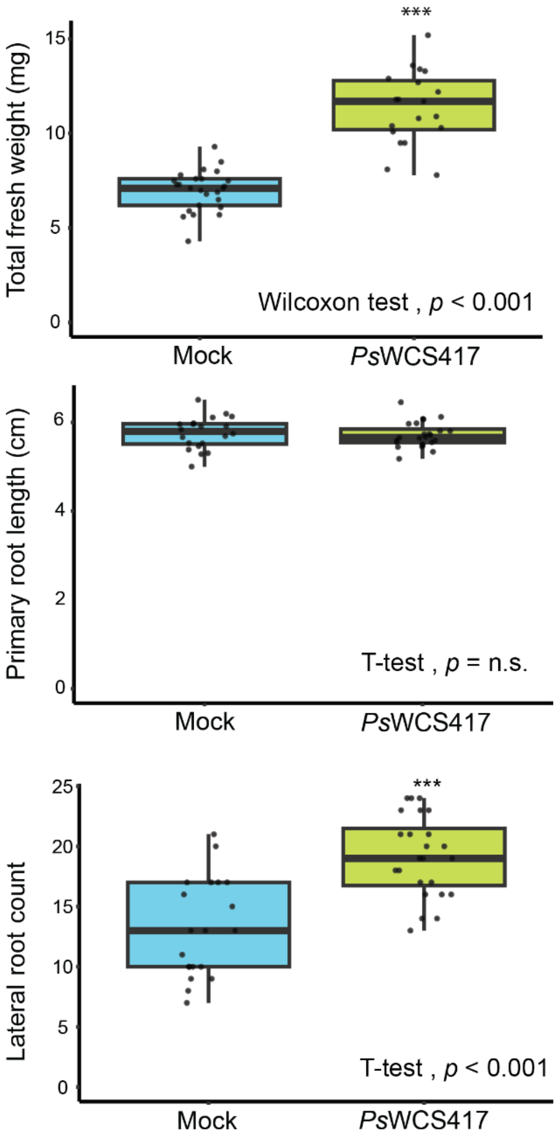


**Fig. S3** Effect of *Ps*WCS417 VOCs on Arabidopsis salt stress tolerance in sterile media. Seedling fresh weight (top), primary root length (middle), and lateral root count (bottom) of Arabidopsis seedlings co-cultivated with *Ps*WCS417 or mock under salt (half-strength MS with 75mM NaCl) conditions. Boxplots show the median, interquartile range, and data distribution. Data was tested for normality and analyzed using a Student's t-test (normal distribution) or a Wilcoxon rank-sum test (non-normal distribution). Asterisks indicate statistically significant differences among treatments ( $P < 0.05$ *, $P < 0.01$ **, $P < 0.001$***) (n= 20-25).

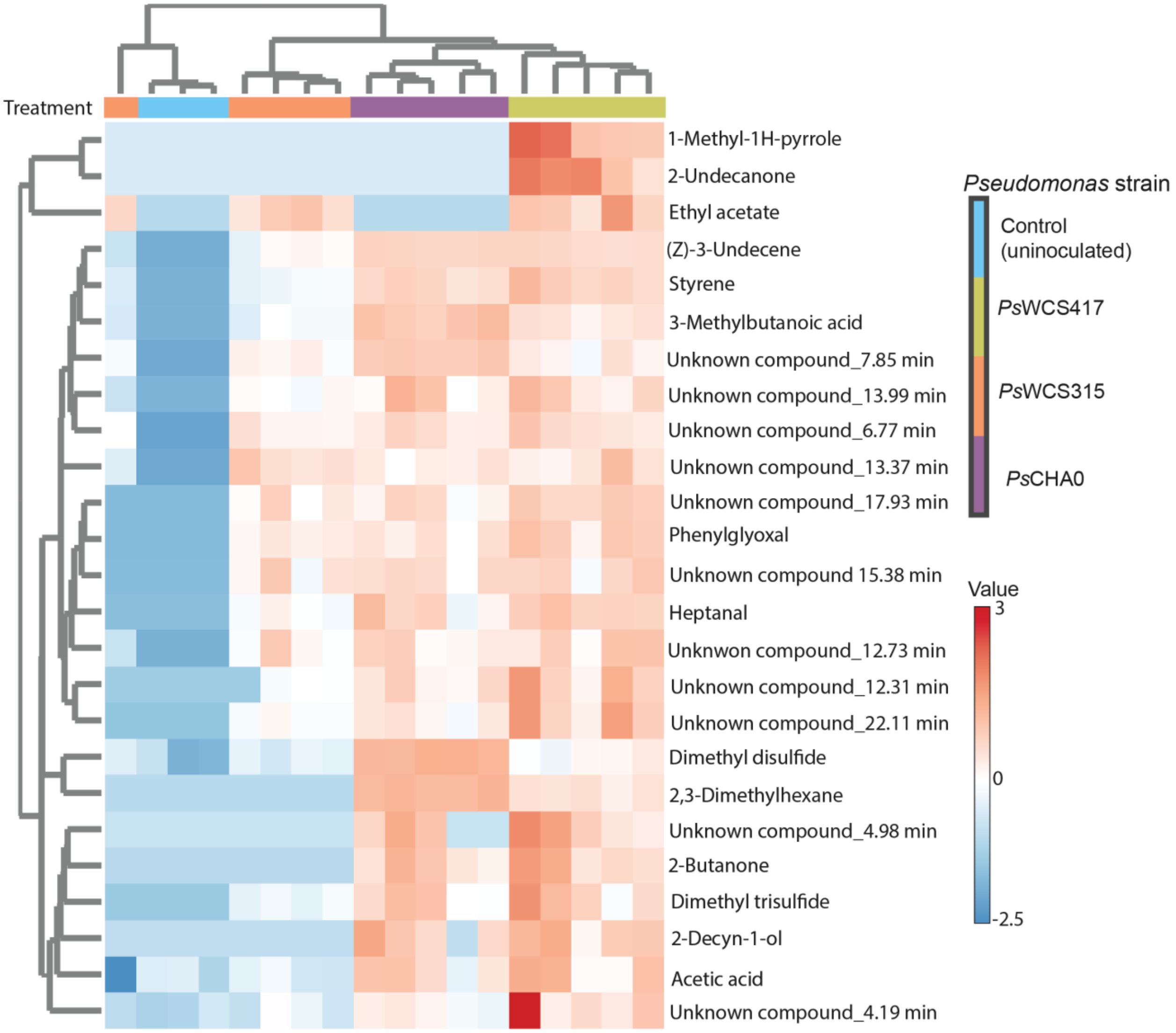


**Fig. S4** Heatmap of VOCs emitted by *Pseudomonas* strains. Each row represents an individual VOC, and colours indicate the relative abundance (red: high, blue: low). Data were log10-transformed and Pareto scaled prior to analysis. Hierarchical clustering groups strains based on their emitted VOCs profiles. Colors indicate *Pseudomonas* strain (blue = control, green = *Ps*WCS417, orange = *Ps*WCS315, purple = *Ps*CHA0)

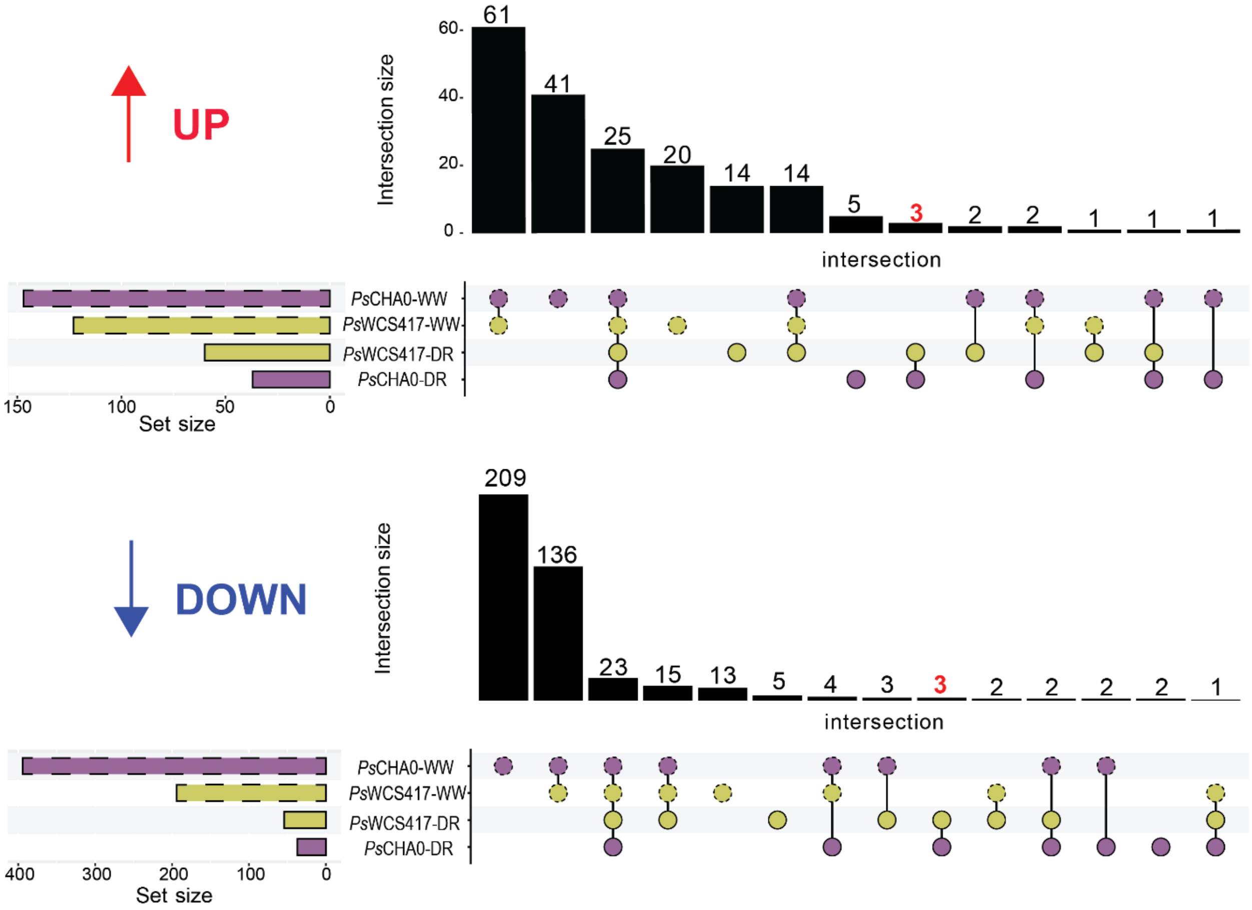


**Fig. S5** Intersection of differentially expressed genes across treatments and water regimes at 24 h visualized using an UpSet plot. Horizontal bars indicate the total number of differentially expressed genes (DEGs) identified for each treatment and water regime combination, while vertical bars represent the number of shared DEGs among combinations of groups. Connected dots denote the specific set combinations contributing to each intersection. The upper panel shows upregulated genes, and the lower panel shown downregulated genes. Only genes meeting the significance threshold ($P < 0.05$, $|\log2fold| \geqslant 1$) were included in the analyses. The *Ps*WCS315 treatment was excluded due to the low number of DEGs detected. Colors indicate treatment (green = *Ps*WCS417, purple = *Ps*CHA0) and outlines indicate water regime (dashed = well-watered, solid = drought).

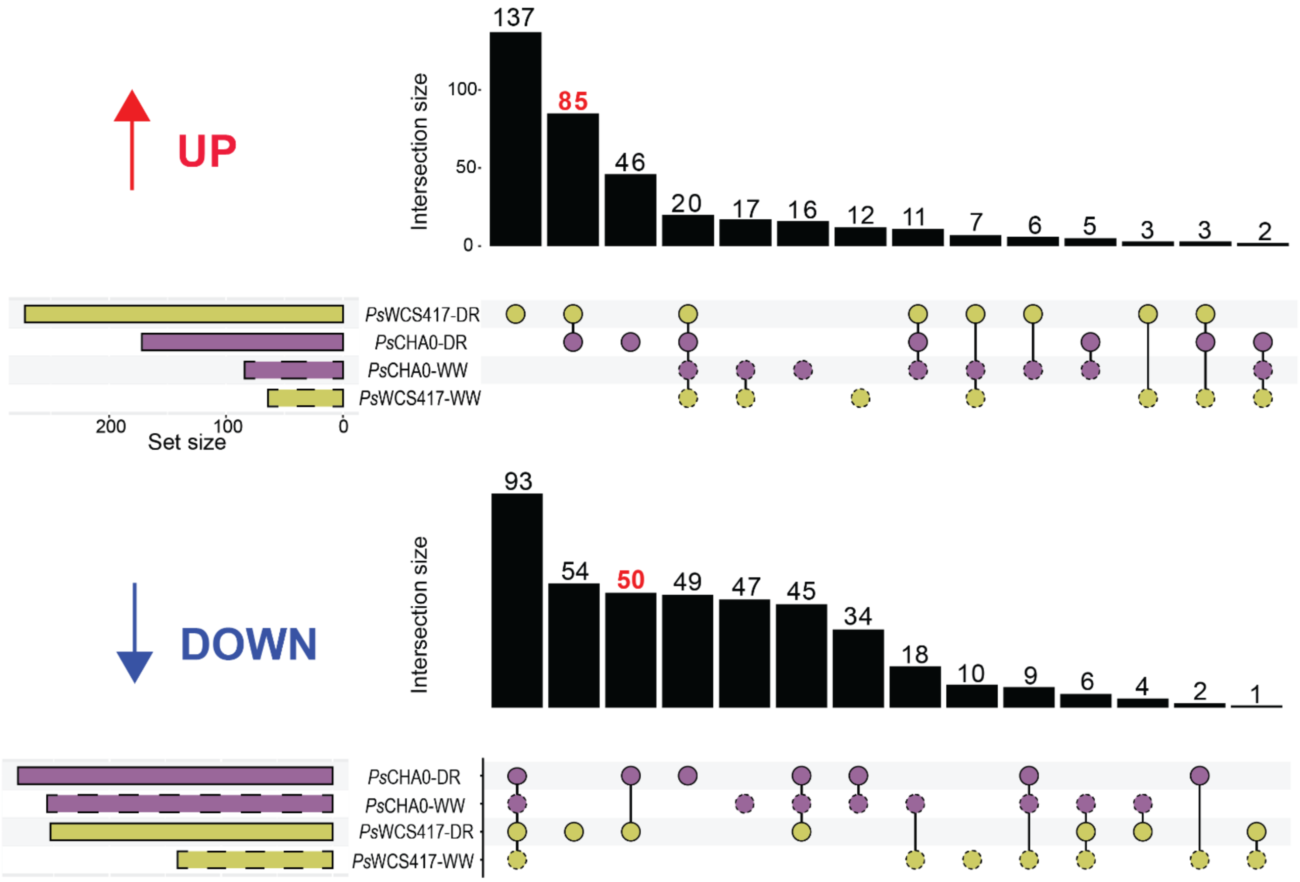


**Fig. S6** Intersection of differentially expressed genes across treatments and water regimes at 48 h visualized using an UpSet plot. Horizontal bars indicate the total number of differentially expressed genes (DEGs) identified for each treatment and water regime combination, while vertical bars represent the number of shared DEGs among combinations of groups. Connected dots denote the specific set combinations contributing to each intersection. The upper panel shows upregulated genes, and the lower panel shown downregulated genes. Only genes meeting the significance threshold ($P < 0.05$, $|\log2fold| \geqslant 1$) were included in the analyses. The *Ps*WCS315 treatment was excluded due to the low number of DEGs detected. Colors indicate treatment (green = *Ps*WCS417, purple = *Ps*CHA0) and outlines indicate water regime (dashed = well-watered, solid = drought).

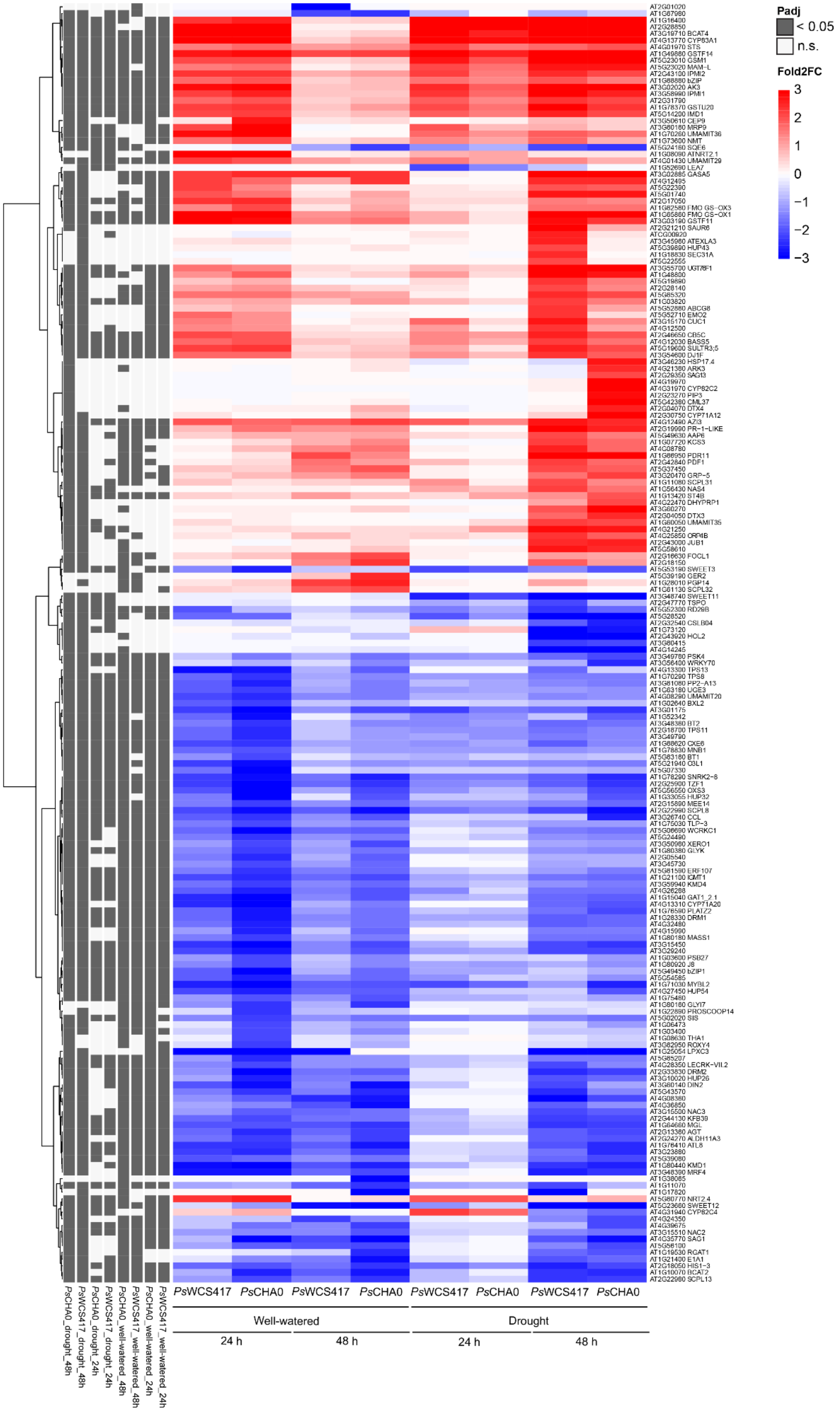
Padj
< 0.05
n.s.
Fold2FC
3
2
1
0
−1
−2
−3
PsWCS417
PsCHA0
PsWCS417
PsCHA0
PsWCS417
PsCHA0
PsWCS417
PsCHA0
Well-watered
Drought
24 h
48 h
24 h
48 h
PsCHA0_drought_48h
PsWCS417_drought_48h
PsCHA0_drought_24h
PsWCS417_drought_24h
PsCHA0_well-watered_48h
PsWCS417_well-watered_48h
PsCHA0_well-watered_24h
PsWCS417_well-watered_24h

**Fig. S7** Heatmap of log2 fold change (L2FC) for differentially expressed genes across *Ps*WCS417 *and Ps*CHA0 treatments and water regime. A heatmap of L2FC values for differentially expressed genes (DEGs), clustered by gene (rows) and treatment combination (columns). L2FC values were clipped at the 95th percentile of their absolute distribution to reduce the impact of extreme values and enhance visualization contrast. Treatments are grouped by *Pseudomonas* strain (*Ps*WCS417 and *Ps*CHA0) , water regime (well-watered and drought) and time point (24 h and 48 h). Gene names (rows) were downloaded from Tair and added to the heatmap. Red indicates upregulation and blue downregulation. Genes included in the plot met the significance criteria ($P < 0.05$, $|\log2fold| \geqslant 2$) in at least one of the groups.

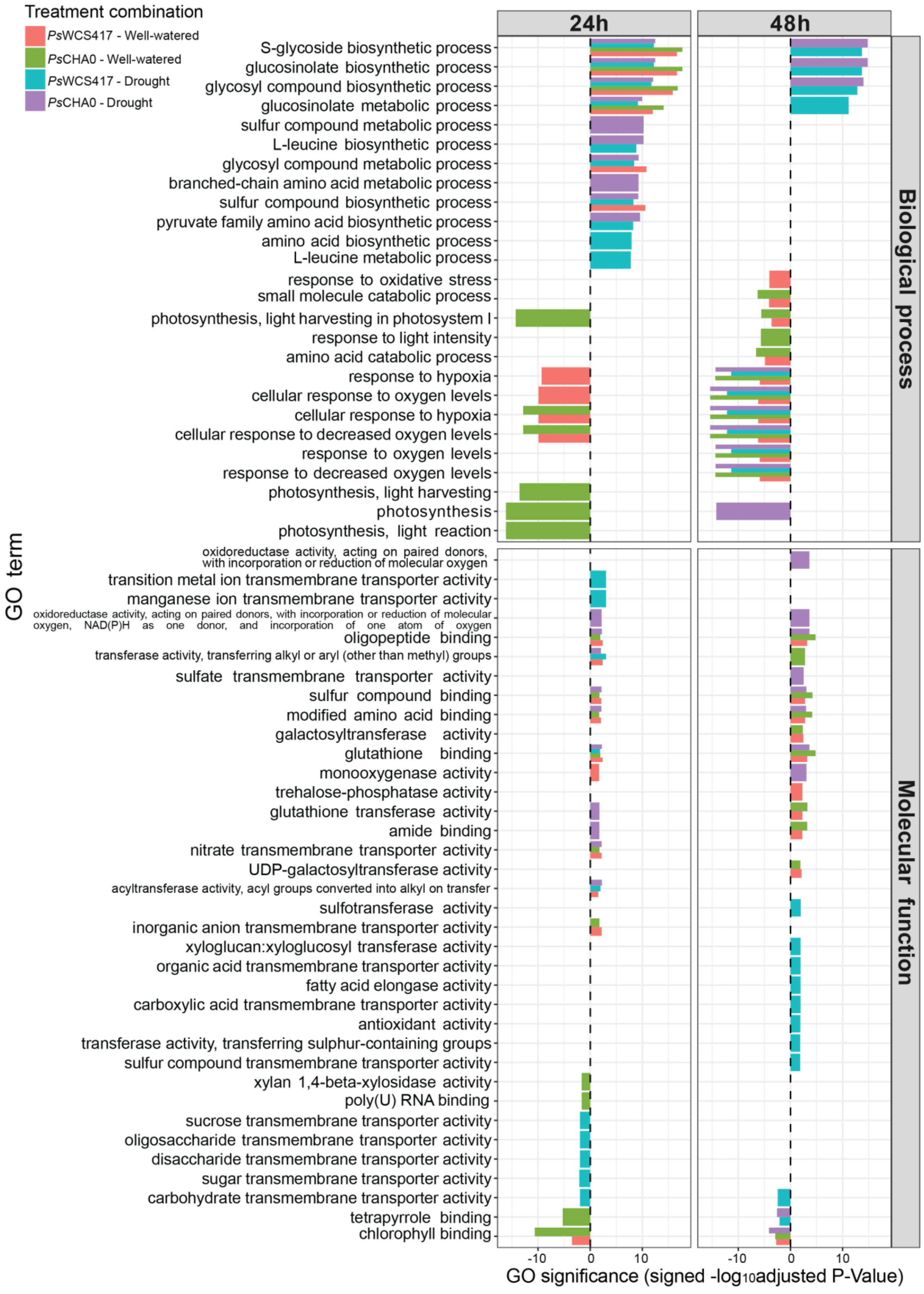

Treatment combination
PsWCS417 - Well-watered
PsCHA0 - Well-watered
PsWCS417 - Drought
PsCHA0 - Drought
24h
48h
Biological process
Molecular function
GO term
S-glycoside biosynthetic process
glucosinolate biosynthetic process
glycosyl compound biosynthetic process
glucosinolate metabolic process
sulfur compound metabolic process
L-leucine biosynthetic process
glycosyl compound metabolic process
branched-chain amino acid metabolic process
sulfur compound biosynthetic process
pyruvate family amino acid biosynthetic process
amino acid biosynthetic process
L-leucine metabolic process
response to oxidative stress
small molecule catabolic process
photosynthesis, light harvesting in photosystem I
response to light intensity
amino acid catabolic process
response to hypoxia
cellular response to oxygen levels
cellular response to hypoxia
cellular response to decreased oxygen levels
response to oxygen levels
response to decreased oxygen levels
photosynthesis, light harvesting
photosynthesis
photosynthesis, light reaction
oxidoreductase activity, acting on paired donors, with incorporation or reduction of molecular oxygen
transition metal ion transmembrane transporter activity
manganese ion transmembrane transporter activity
oxidoreductase activity, acting on paired donors, with incorporation or reduction of molecular oxygen, NAD(P)H as one donor, and incorporation of one atom of oxygen
oligopeptide binding
transferase activity, transferring alkyl or aryl (other than methyl) groups
sulfate transmembrane transporter activity
sulfur compound binding
modified amino acid binding
galactosyltransferase activity
glutathione binding
monooxygenase activity
trehalose-phosphatase activity
glutathione transferase activity
amide binding
nitrate transmembrane transporter activity
UDP-galactosyltransferase activity
acyltransferase activity, acyl groups converted into alkyl on transfer
sulfotransferase activity
inorganic anion transmembrane transporter activity
xyloglucan:xyloglucosyl transferase activity
organic acid transmembrane transporter activity
fatty acid elongase activity
carboxylic acid transmembrane transporter activity
antioxidant activity
transferase activity, transferring sulphur-containing groups
sulfur compound transmembrane transporter activity
xylan 1,4-beta-xylosidase activity
poly(U) RNA binding
sucrose transmembrane transporter activity
oligosaccharide transmembrane transporter activity
disaccharide transmembrane transporter activity
sugar transmembrane transporter activity
carbohydrate transmembrane transporter activity
tetrapyrrole binding
chlorophyll binding
-10
0
10
-10
0
10
GO significance (signed -log10adjusted P-Value)

**Fig. S8** Gene ontology (GO) enrichment of differentially expressed genes. GO enrichment was performed separately for each *Ps*VOC treatment, water regime, and timepoint using differentially expressed genes identify by RNA-seq analyses ($\mathrm{FDR} < 0.05$ and ｜fold2change｜ $\geqslant$ 1). Bars represent the significance of enrichment of GO terms. The x-axis shows signed enrichment significance (-log10 adjusted P-value), where positive values indicate upregulated genes and negative values indicate enrichment among downregulated genes. Bar color denotes the *Ps*VOC x water regime treatment combination. The top 10 GO terms were selected for each treatment combination.

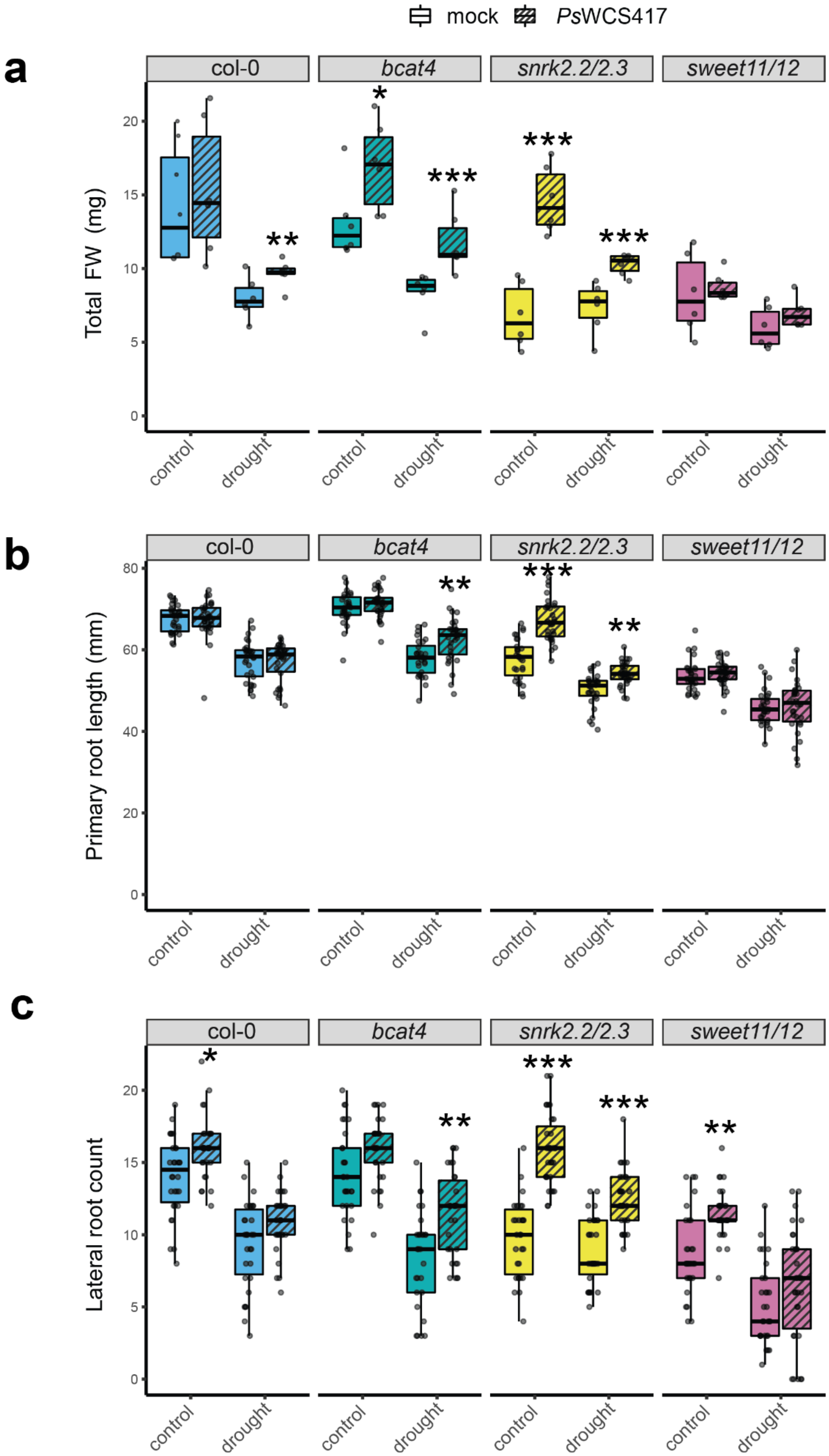

mock
PsWCS417
a
col-0
bcat4
snrk2.2/2.3
sweet11/12
Total FW (mg)
control
drought
b
Primary root length (mm)
c
Lateral root count

**Fig. S9** Effect of *Ps*WCS417 VOCs in Col-0, bcat-4, snrk2.2/2.3 and sweet11/12 Arabidopsis lines grown *in vitro*. A) Total fresh weight (root and shoot), B) primary root length and C) lateral root count of seedlings grown wither in control (half-strength MS) or drought (half-strength MS with sorbitol) conditions and exposed to mock or *Ps*WCS417 VOCs. Boxplots display median, interquartile range, and data distribution. Significant differences due to *Ps*WCS417 VOCs expositions within each line and growth media are indicated (GLMM, ANOVA II test; * = $p < 0.05$; ** = $p < 0.01$; *** = $p < 0.001$).

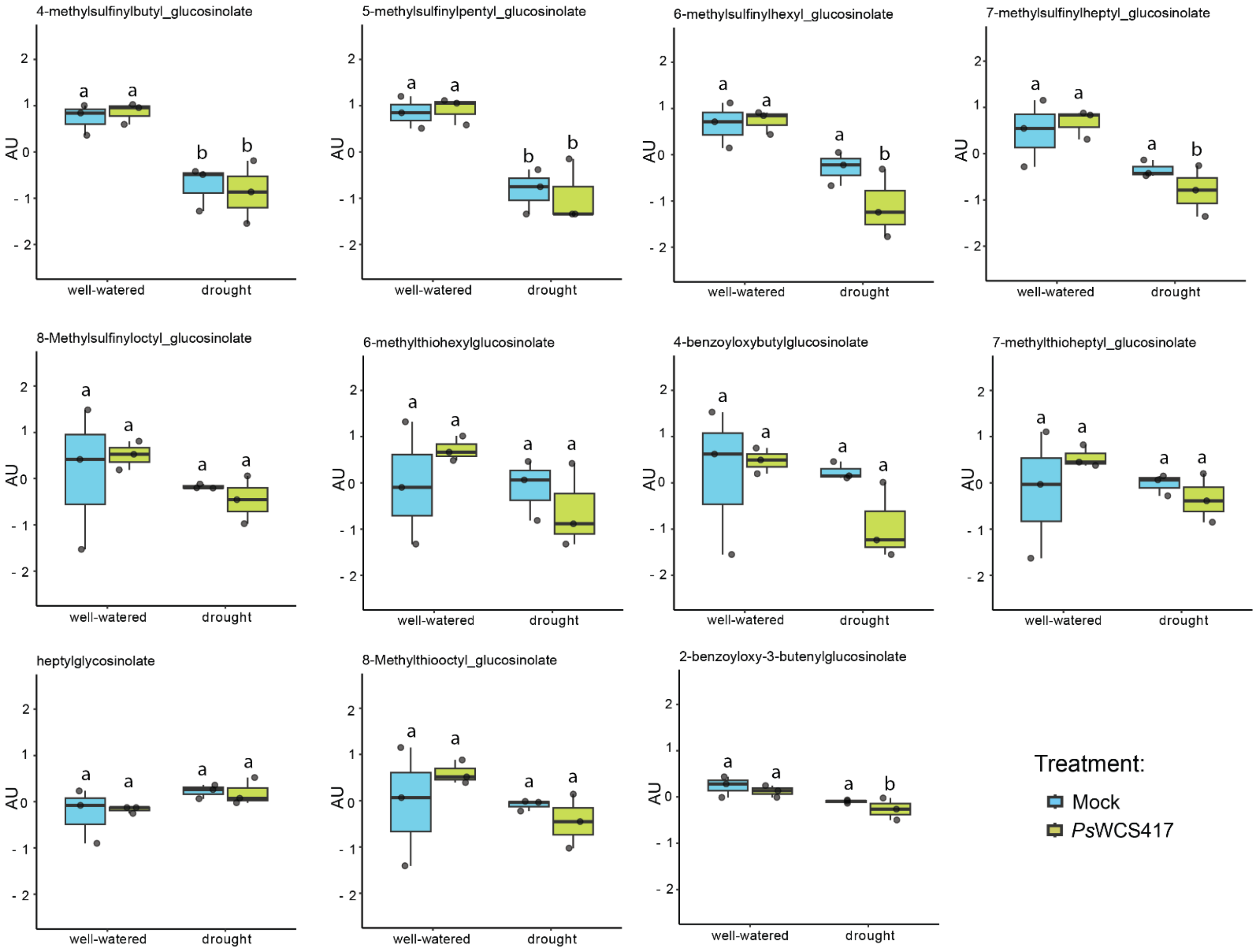


**Fig. S10** Effect of *Ps*WCS417 VOCs on aliphatic glucosinolate levels (relative abundance) in roots of axenically grown Arabidopsis under well-watered and drought conditions. Aliphatic glucosinolate levels are shown as arbitrary units (AU). The x-axis indicates water regime (well-watered and drought). Colors indicate treatment (blue = mock, green *Ps*WCS417) Data were log-transformed and Pareto scaled prior to analysis. Boxplots show the median, interquartile range, and data distribution. Data were analysed by PERMANOVA, and different letters indicate statistically significant differences among groups (n = 3)

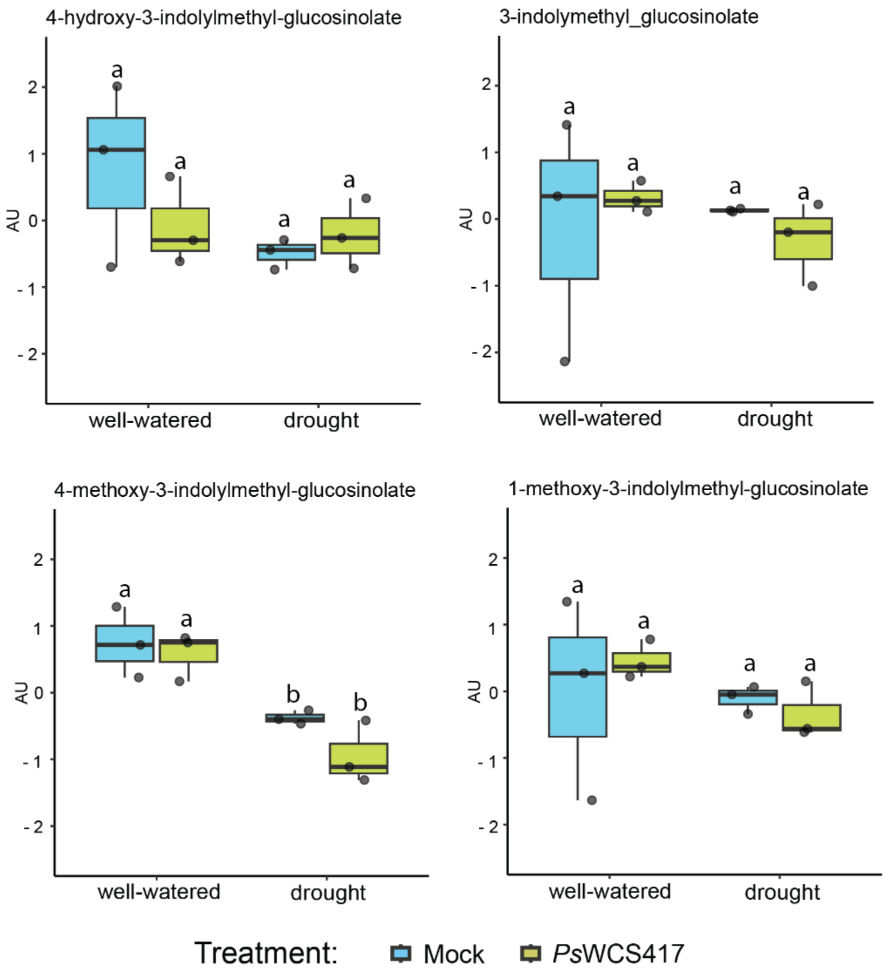


**Fig. S11** Effect of *Ps*WCS417 VOCs on indolic glucosinolate levels (relative abundance) in roots of axenically grown Arabidopsis under well-watered and drought conditions. Indolic glucosinolate levels are shown as arbitrary units (AU). The x-axis indicates water regime (well-watered and drought). Colors indicate treatment (blue = mock, green *Ps*WCS417) Data were log-transformed and Pareto scaled prior to analysis. Boxplots show the median, interquartile range, and data distribution. Data were analysed by PERMANOVA, and different letters indicate statistically significant differences among groups (n = 3)

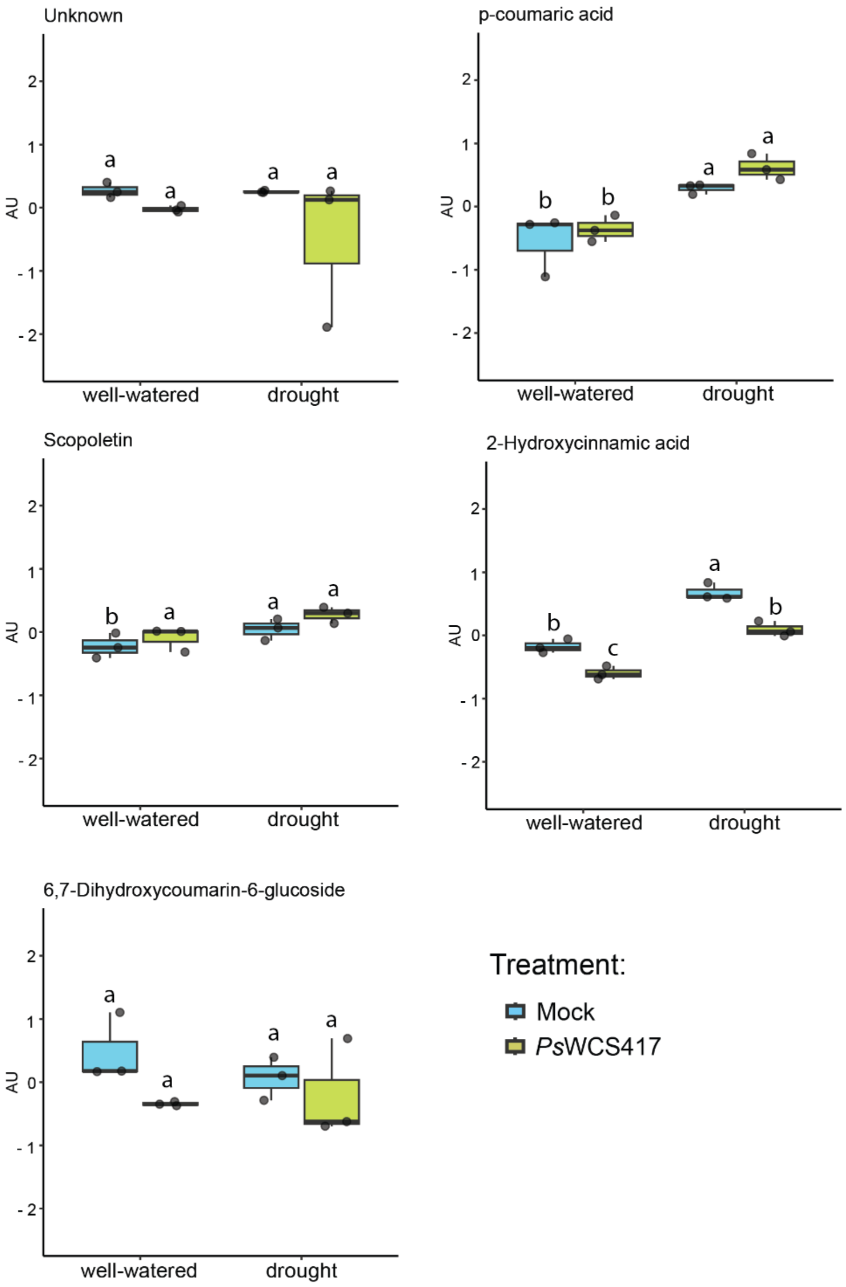


**Fig. S12** Effect of *Ps*WCS417 VOCs on coumarin levels (relative abundance) in roots of axenically grown Arabidopsis under well-watered and drought conditions. Coumarin levels are shown as arbitrary units (AU). The x-axis indicates water regime (well-watered and drought). Colors indicate treatment (blue = mock, green *Ps*WCS417) Data were log-transformed and Pareto scaled prior to analysis. Boxplots show the median, interquartile range, and data distribution. Data were analysed by PERMANOVA, and different letters indicate statistically significant differences among groups (n = 3)

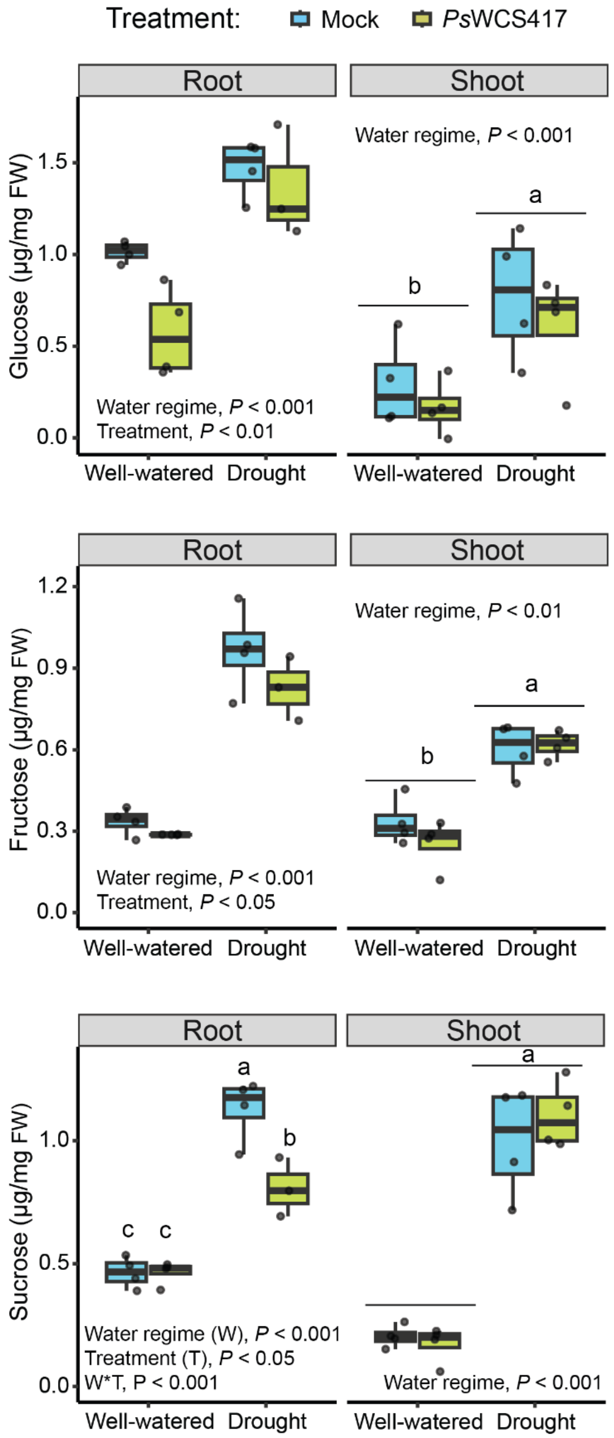


**Fig. S13** Effect of *Ps*WCS417 VOCs on plant sugar content. Boxplots show the concentration of (A) glucose, (B) fructose and (C) sucrose in roots and shoots of axenically grown Arabidopsis seedlings under well-watered or drought conditions, either mock-treated or exposed to *Ps*WCS417 VOCs. Boxplots show the median, interquartile range, and data distribution. Data were analyzed using two-way ANOVA followed by Tukey's post hoc test. Different letters indicate statistically significant differences among groups (n = 4).

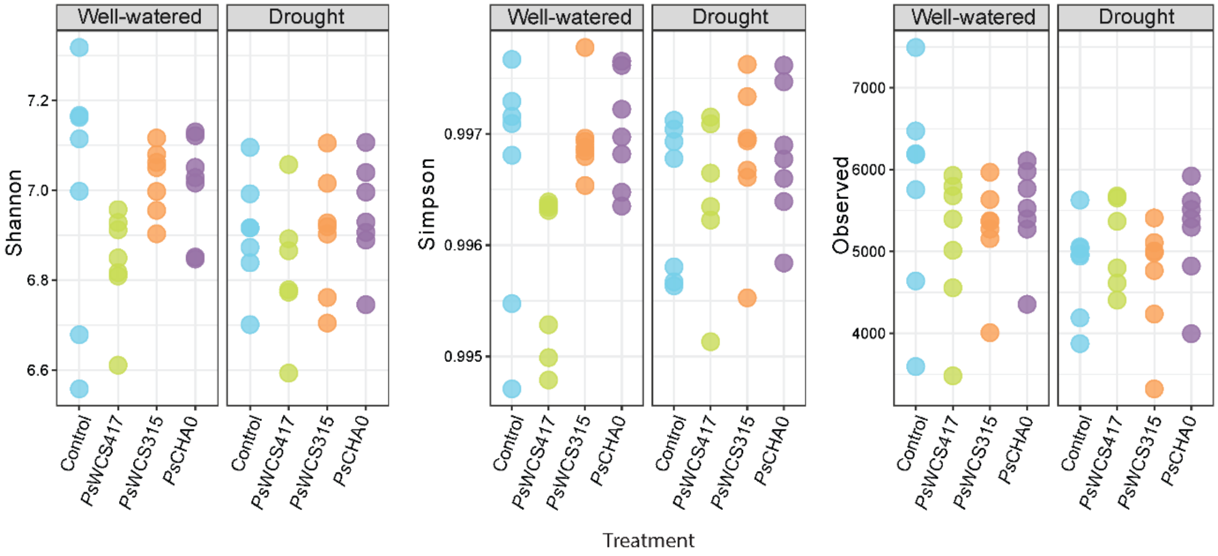


**Fig. S14** Alpha diversity analysis**.** Alpha diversity metrics for well-watered and drought samples exposed to different *Pseudomonas* strains. Shannon diversity index, Simpson diversity index and Observed species richness in plant root microbiome.

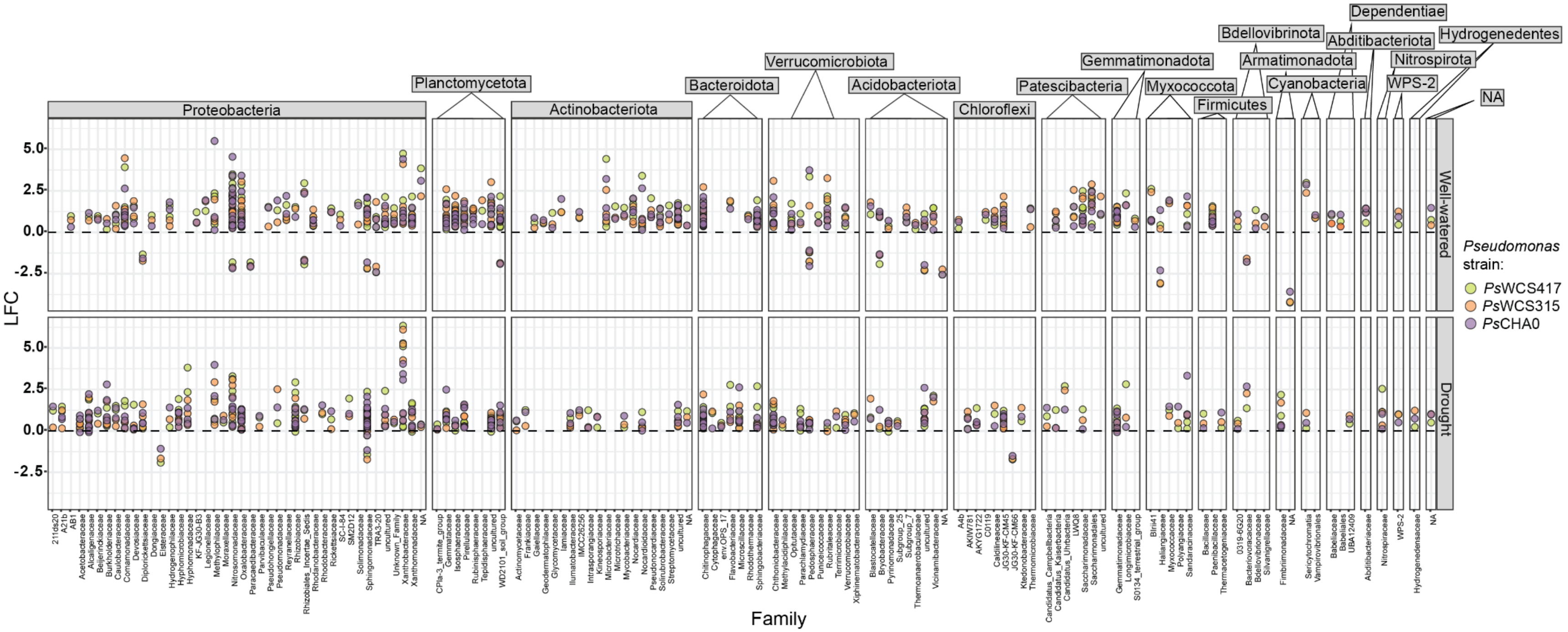

Proteobacteria
Planctomycetota
Actinobacteriota
Bacteroidota
Verrucomicrobiota
Acidobacteriota
Chloroflexi
Patescibacteria
Gemmatimonadota
Myxococcota
Firmicutes
Bdellovibrionota
Armatimonadota
Cyanobacteria
Dependentiae
Abditibacteriota
Hydrogenedentes
Nitrospirota
WPS-2
NA
Well-watered
Drought
LFC
5.0
2.5
0.0
-2.5
Family
Pseudomonas strain:
PsWCS417
PsWCS315
PsCHA0

**Fig. S15** Impact of *Pseudomonas* VOCs on the *B. oleracea* root microbiome under well-watered and drought conditions. Logarithmic fold change (LFC) values of ASVs enriched or depleted by the three different Pseudomonas strains under well-watered (top) and drought (bottom) conditions. Y-axis shows LFC value and X-axis the bacterial families. Each dots represents an individual ASV, color-coded according to the *Pseudomonas* strains used for the treatments. Families are grouped by bacterial phylum.

**Table S1** Primers used for the genotyping of the Arabidopsis mutant lines.

| Candidate gene | Forward primer (5' → 3') | Reverse primer (5' → 3') |
|---|---|---|
| **BCAT4-1** | AAGTCTCCTCAGGCTAGACCG | AGGGCA ATGTTGTATCGACAC |
| **SNRK2.2** | ACCGCAAGACCATACATCTGCAAGC | GCGGATTTGTAATGCCGGACGGT |
| **SNRK2.3** | GGTAGCGGGATCAGCCACGAAG | TGGCGGTGAACTTTACGAGCGG |
| **SWEET11** | CCGAAGAGTAATGTGACCACG | TGAAGTGGGTGCTTTTGTTTC |
| **SWEET12** | TCAAAGGCCAAAGCAATATACC | ATGCAGGCCAACGTTCTATAG |

**Table S2** Type III Analysis of Deviance (Wald Chi-Square Test) for Seedling Biomass under well-watered conditions. Results from a generalized linear mixed model (GLMM) assessing the effect of treatment on total seedling biomass (mg) when grown in control conditions. The model included treatment as a fixed effect and plate ID as a random effect.

| Factor | Chi² | DF | p-value |
|---|---|---|---|
| (Intercept) | 452.59 | 1 | < 0.001 |
| **treatment** | **11.94** | **3** | **0.008** |

**Table S3** Pairwise Comparisons of Treatments Using Tukey's HSD. Estimated differences in total seedling biomass between treatments when grown in control conditions, based on Tukey-adjusted post-hoc tests. Significant pairwise differences ($p < 0.05$) are highlighted in bold. Degrees of freedom (DF) reflect the model's residuals.

| Contrast | Estimate | SE | DF | t-ratio | p-value |
|---|---|---|---|---|---|
| **Ctrl - WCS417** | **-0.403** | **0.134** | **14** | **-2.996** | **0.042** |
| **Ctrl - WCS315** | **-0.427** | **0.134** | **14** | **-3.182** | **0.030** |
| Ctrl - CHA0 | -0.362 | 0.136 | 14 | -2.664 | 0.078 |
| WCS417 - WCS315 | -0.024 | 0.116 | 14 | -0.209 | 0.997 |
| WCS417 - CHA0 | 0.041 | 0.117 | 14 | 0.352 | 0.984 |
| WCS315 - CHA0 | 0.066 | 0.117 | 14 | 0.560 | 0.942 |

**Table S4** Compact Letter Display (CLD) of Estimated Means Across Treatments. Estimated marginal means of total seedling biomass per treatment with 95% confidence intervals.

Treatments not sharing a letter are significantly different (p < 0.05) based on multiple comparison tests using the multcomp package. Standard errors and degrees of freedom are included for reference.

| Treatment | Estimated Mean | SE | df | lower.CL | upper.CL | Group |
|---|---|---|---|---|---|---|
| Ctrl | 2.27 | 0.10683523 | 14 | 1.967879 | 2.577758 | a |
| CHA0 | 2.63 | 0.08440094 | 14 | 2.393439 | 2.875250 | ab |
| WCS417 | 2.68 | 0.08242024 | 14 | 2.440461 | 2.910964 | ab |
| WCS315 | 2.70 | 0.08296408 | 14 | 2.463128 | 2.936736 | b |

**Table S5** Type III Analysis of Deviance (Wald Chi-Square Test) for Seedling Biomass under drought conditions. Results from a generalized linear mixed model (GLMM) assessing the effect of treatment on total seedling biomass (mg) when grown in drought conditions. The model included treatment as a fixed effect and plate ID as a random effect.

| Factor | $Chi^2$ | DF | p-value |
|---|---|---|---|
| (Intercept) | 1,165.14 | 1 | < 0.001 |
| treatment | 77.09 | 3 | < 0.001 |

**Table S6** Pairwise Comparisons of Treatments Using Tukey's HSD. Estimated differences in total seedling biomass between treatments when grown in drought conditions, based on Tukey-adjusted post-hoc tests. Significant pairwise differences (p < 0.05) are highlighted in bold. Degrees of freedom (DF) reflect the model's residuals

| Contrast | Estimate | SE | DF | t-ratio | p-value |
|---|---|---|---|---|---|
| **Ctrl - WCS417** | **-0.603** | **0.078** | **14** | **-7.715** | **<0.001** |
| **Ctrl - WCS315** | **-0.598** | **0.079** | **14** | **-7.576** | **<0.001** |
| **Ctrl - CHA0** | **-0.364** | **0.079** | **14** | **-4.588** | **0.002** |
| WCS417 - WCS315 | 0.005 | 0.076 | 14 | 0.064 | 1.000 |
| **WCS417 - CHA0** | **0.239** | **0.076** | **14** | **3.135** | **0.033** |
| **WCS315 - CHA0** | **0.234** | **0.076** | **14** | **3.084** | **0.036** |

**Table S7** Compact Letter Display (CLD) of Estimated Means Across Treatments. Estimated marginal means of total seedling biomass per treatment with 95% confidence intervals. Treatments not sharing a letter are significantly different (p < 0.05) based on multiple comparison tests using the multcomp package. Standard errors and degrees of freedom are included for reference.

| Treatment | Estimated Mean | SE | df | lower.CL | upper.CL | Group |
|---|---|---|---|---|---|---|
| Ctrl | 1.98 | 0.05795795 | 14 | 1.812920 | 2.143778 | a |
| CHA0 | 2.34 | 0.05415606 | 14 | 2.187655 | 2.496810 | b |
| WCS315 | 2.58 | 0.05306090 | 14 | 2.424628 | 2.727532 | c |
| WCS417 | 2.58 | 0.05354612 | 14 | 2.428107 | 2.733781 | c |

**Table S8** Type III Analysis of Deviance (Wald Chi-Square Test) for Seedling Primary Root length under well-watered conditions. Results from a generalized linear mixed model (GLMM) assessing the effect of treatment on seedling primary root length (mm) when grown in control conditions. The model included treatment as a fixed effect and plate ID as a random effect.

| Factor | $Chi^2$ | DF | p-value |
|---|---|---|---|
| (Intercept) | 62,442.67 | 1 | < 0.001 |
| treatment | 32.63 | 3 | < 0.001 |

**Table S9** Pairwise Comparisons of Treatments Using Tukey's HSD. Estimated differences in seedling primary root length between treatments when grown in control conditions, based on Tukey-adjusted post-hoc tests. Significant pairwise differences (p < 0.05) are highlighted in bold. Degrees of freedom (DF) reflect the model's residuals.

| Contrast | Estimate | SE | DF | t-ratio | p-value |
|---|---|---|---|---|---|
| **CHA0 - Ctrl** | **0.137** | **0.025** | **91** | **5.498** | **<0.001** |
| CHA0 - WCS315 | 0.056 | 0.024 | 91 | 2.271 | 0.112 |
| CHA0 - WCS417 | 0.030 | 0.024 | 91 | 1.269 | 0.585 |
| **Ctrl - WCS315** | **-0.081** | **0.025** | **91** | **-3.182** | **0.011** |
| **Ctrl - WCS417** | **-0.106** | **0.025** | **91** | **-4.244** | **<0.001** |
| WCS315 - WCS417 | -0.025 | 0.025 | 91 | -1.020 | 0.738 |

**Table S10** Compact Letter Display (CLD) of Estimated Means Across Treatments.
Estimated marginal means of primary seedling length (mm) per treatment with 95% confidence intervals. Treatments not sharing a letter are significantly different (p < 0.05) based on multiple comparison tests using the multcomp package. Standard errors and degrees of freedom are included for reference.

| Treatment | Estimated Mean | SE | df | lower.CL | upper.CL | Group |
|---|---|---|---|---|---|---|
| Ctrl | 4.08 | 0.01825054 | 91 | 4.030087 | 4.122842 | a |
| WCS315 | 4.16 | 0.01775395 | 91 | 4.112357 | 4.202588 | b |
| WCS417 | 4.18 | 0.01710179 | 91 | 4.139150 | 4.226067 | b |
| CHA0 | 4.21 | 0.01686005 | 91 | 4.170234 | 4.255922 | b |

**Table S11** Type III Analysis of Deviance (Wald Chi-Square Test) for Seedling Primary Root length under drought conditions. Results from a generalized linear mixed model (GLMM) assessing the effect of treatment on seedling primary root length (mm) when grown in drought conditions. The model included treatment as a fixed effect and plate ID as a random effect.

| Factor | $Chi^2$ | DF | p-value |
|---|---|---|---|
| (Intercept) | 2,575.32 | 1 | < 0.001 |
| treatment | 5.72 | 3 | 0.126 |

**Table S12** Pairwise Comparisons of Treatments Using Tukey's HSD. Estimated differences in seedling primary root length between treatments when grown in drought conditions, based on Tukey-adjusted post-hoc tests. Significant pairwise differences ($p < 0.05$) are highlighted in bold. Degrees of freedom (DF) reflect the model's residuals.

| Contrast | Estimate | SE | DF | t-ratio | p-value |
|---|---|---|---|---|---|
| CHA0 - Ctrl | 3.569 | 1.618 | 88 | 2.206 | 0.129 |
| CHA0 - WCS315 | 2.082 | 1.568 | 88 | 1.328 | 0.548 |
| CHA0 - WCS417 | 2.971 | 1.592 | 88 | 1.867 | 0.250 |
| Ctrl - WCS315 | -1.487 | 1.618 | 88 | -0.919 | 0.795 |
| Ctrl - WCS417 | -0.598 | 1.640 | 88 | -0.364 | 0.983 |
| WCS315 - WCS417 | 0.889 | 1.592 | 88 | 0.559 | 0.944 |

**Table S13** Compact Letter Display (CLD) of Estimated Means Across Treatments.
Estimated marginal means of seedling primary root length (mm) per treatment with 95% confidence intervals. Treatments not sharing a letter are significantly different ($p < 0.05$) based on multiple comparison tests using the multcomp package. Standard errors and degrees of freedom are included for reference.

| Treatment | Estimated Mean | SE | df | lower.CL | upper.CL | Group |
|---|---|---|---|---|---|---|
| Ctrl | 52.70 | 1.177660 | 88 | 49.70806 | 55.69740 | a |
| WCS417 | 53.30 | 1.142080 | 88 | 50.39611 | 56.20451 | a |
| WCS315 | 54.19 | 1.108854 | 88 | 51.36964 | 57.00905 | a |
| CHA0 | 56.27 | 1.108854 | 88 | 53.45194 | 59.09135 | a |

**Table S14** Type III Analysis of Deviance (Wald Chi-Square Test) for Seedling Lateral Root Count under well-watered conditions. Results from a generalized linear mixed model (GLMM) assessing the effect of treatment on seedling lateral root count when grown in control conditions. The model included treatment as a fixed effect and plate ID as a random effect

| Factor | Chi² | DF | p-value |
|---|---|---|---|
| (Intercept) | 385.93 | 1 | < 0.001 |
| treatment | 33.15 | 3 | < 0.001 |

**Table S15** Pairwise Comparisons of Treatments Using Tukey's HSD. Estimated differences in seedling lateral root count between treatments when grown in control conditions, based on Tukey-adjusted post-hoc tests. Significant pairwise differences ($p < 0.05$) are highlighted in bold. Degrees of freedom (DF) reflect the model's residuals.

| Contrast | Estimate | SE | DF | t-ratio | p-value |
|---|---|---|---|---|---|
| **CHA0 - Ctrl** | **5.303** | **1.097** | **91** | **4.834** | **<0.001** |
| CHA0 - WCS315 | 1.934 | 1.105 | 91 | 1.751 | 0.304 |
| CHA0 - WCS417 | -0.320 | 1.091 | 91 | -0.293 | 0.991 |
| **Ctrl - WCS315** | **-3.369** | **1.110** | **91** | **-3.035** | **0.016** |
| **Ctrl - WCS417** | **-5.623** | **1.097** | **91** | **-5.126** | **<0.001** |
| WCS315 - WCS417 | -2.254 | 1.105 | 91 | -2.040 | 0.181 |

**Table S16** Compact Letter Display (CLD) of Estimated Means Across Treatments.
Estimated marginal means of seedling lateral root count per treatment with 95% confidence intervals. Treatments not sharing a letter are significantly different ($p < 0.05$) based on multiple comparison tests using the multcomp package. Standard errors and degrees of freedom are included for reference.

| Treatment | Estimated Mean | SE | df | lower.CL | upper.CL | Group |
|---|---|---|---|---|---|---|
| Ctrl | 9.86 | 0.7797731 | 91 | 7.87501 | 11.83806 | a |
| WCS315 | 13.23 | 0.7906188 | 91 | 11.21692 | 15.23509 | b |
| CHA0 | 15.16 | 0.7716931 | 91 | 13.19901 | 17.12100 | b |
| WCS417 | 15.48 | 0.7716931 | 91 | 13.51901 | 17.44100 | b |

**Table S17** Type III Analysis of Deviance (Wald Chi-Square Test) for Seedling Lateral Root Count under drought conditions. Results from a generalized linear mixed model (GLMM) assessing the effect of treatment on seedling lateral root count when grown in drought conditions. The model included treatment as a fixed effect and plate ID as a random effect

| Factor | Chi² | DF | p-value |
|---|---|---|---|
| (Intercept) | 447.39 | 1 | < 0.001 |
| treatment | 57.90 | 3 | < 0.001 |

**Table S18** Pairwise Comparisons of Treatments Using Tukey's HSD. Estimated differences in total seedling lateral root count between treatments when grown in drought conditions, based on Tukey-adjusted post-hoc tests. Significant pairwise differences ($p < 0.05$) are highlighted in bold. Degrees of freedom (DF) reflect the model's residuals.

| Contrast | Estimate | SE | DF | t-ratio | p-value |
|---|---|---|---|---|---|
| **CHA0 - Ctrl** | **5.290** | **0.871** | **88** | **6.071** | **<0.001** |
| CHA0 - WCS315 | -0.120 | 0.840 | 88 | -0.143 | 0.999 |
| CHA0 - WCS417 | -0.547 | 0.855 | 88 | -0.639 | 0.919 |
| **Ctrl - WCS315** | **-5.410** | **0.871** | **88** | **-6.209** | **<0.001** |
| **Ctrl - WCS417** | **-5.836** | **0.887** | **88** | **-6.582** | **<0.001** |
| WCS315 - WCS417 | -0.427 | 0.855 | 88 | -0.499 | 0.959 |

**Table S19** Compact Letter Display (CLD) of Estimated Means Across Treatments.
Estimated marginal means of seedling lateral root count per treatment with 95% confidence intervals. Treatments not sharing a letter are significantly different ($p < 0.05$) based on multiple comparison tests using the multcomp package. Standard errors and degrees of freedom are included for reference.

| Treatment | Estimated Mean | SE | df | lower.CL | upper.CL | Group |
|---|---|---|---|---|---|---|
| Ctrl | 7.27 | 0.6376201 | 88 | 5.648856 | 8.891666 | a |
| CHA0 | 12.56 | 0.5938105 | 88 | 11.049975 | 14.069978 | b |
| WCS315 | 12.68 | 0.5938105 | 88 | 11.170000 | 14.190003 | b |
| WCS417 | 13.11 | 0.6150464 | 88 | 11.542515 | 14.670520 | b |

**Table S20** PERMANOVA Results (Bray-Curtis Dissimilarity) on volatile blends of *Pseudomonas* strains and mock. Permutational multivariate analysis of variance (PERMANOVA) results testing the differences on volatile blend composition among mock and Pseudomonas strains. Significant effects (p<0.05) are considered statistically significant

| Factor | DF | Sum Sq | $R^2$ | F-value | p-value |
|---|---|---|---|---|---|
| **strain** | **3** | **561.49** | **0.48** | **4.3** | **< 0.001** |
| **Residual** | **14** | **608.96** | **0.52** | | **NA** |
| **Total** | **17** | **1,170.45** | **1.00** | | **NA** |

**Table S21** Pairwise PERMANOVA Results on volatile blends of *Pseudomonas* strains and mock.

| Comparison | Term | Df | SumOfSqs | R2 | F | Pr(>F) |
|---|---|---|---|---|---|---|
| CHA0_vs_WCS417 | strain | 1 | 86.61 | 0.173 | 1.670 | 0.200 |
| | Residual | 8 | 414.84 | 0.827 | | |
| | Total | 9 | 501.45 | 1.000 | | |
| CHA0_vs_Control | strain | 1 | 236.86 | 0.589 | 8.608 | **0.022** |
| | Residual | 6 | 165.10 | 0.411 | | |
| | Total | 7 | 401.96 | 1.000 | | |
| CHA0_vs_WCS315 | strain | 1 | 148.81 | 0.328 | 3.906 | **0.018** |
| | Residual | 8 | 304.74 | 0.672 | | |
| | Total | 9 | 453.54 | 1.000 | | |
| WCS417_vs_Control | strain | 1 | 348.00 | 0.534 | 6.863 | **0.035** |
| | Residual | 6 | 304.22 | 0.466 | | |
| | Total | 7 | 652.22 | 1.000 | | |
| WCS417_vs_WCS315 | strain | 1 | 255.17 | 0.365 | 4.599 | **0.020** |
| | Residual | 8 | 443.86 | 0.635 | | |
| | Total | 9 | 699.03 | 1.000 | | |
| Control_vs_WCS315 | strain | 1 | 65.25 | 0.252 | 2.017 | 0.111 |

| Comparison | Term | Df | SumOfSqs | R2 | F | Pr(>F) |
|---|---|---|---|---|---|---|
| | Residual | 6 | 194.12 | 0.748 | | |
| | Total | 7 | 259.37 | 1.000 | | |

**Table S22** PERMANOVA Results Based on PCA Scores. Permutational multivariate analysis of variance (PERMANOVA) results testing the effects of *Pseudomonas* strain, time point and water regime on PCA scores derived from multivariate data. The analysis was based on Euclidean distances. Significant effects (p<0.05) are considered statistically significant

| Factor | DF | Sum Sq | $R^2$ | F-value | p-value |
|---|---|---|---|---|---|
| **Pseudomonas** | **3** | **2,835.99** | **0.304** | **56.71** | **0.001** |
| **Time** | **1** | **1,215.13** | **0.130** | **72.90** | **0.001** |
| **Water regime** | **1** | **4,578.38** | **0.491** | **274.67** | **0.001** |
| **Residual** | **42** | **700.09** | **0.075** | | **NA** |
| **Total** | **47** | **9,329.59** | **1.000** | | **NA** |

**Table S23** Pairwise PERMANOVA results based on Euclidean distances of PCA scores from RNA-seq data in Arabidopsis roots. 999 permutations, Bonferroni adjusted p-values. Each model tested the pairwise effect of *Pseudomonas* strain, time point and water regime. Significant effects (p < 0.05) are marked in bold.

**Comparison: control_vs_WCS417**

| Factor | DF | Sum Sq | $R^2$ | F-value | p-value |
|---|---|---|---|---|---|
| **Pseudomonas** | **1** | **1,909.752** | **0.375** | **103.167** | **0.001** |
| **Time** | **1** | **641.816** | **0.126** | **34.672** | **0.001** |
| **Treatment** | **1** | **2,168.012** | **0.426** | **117.119** | **0.001** |
| Residual | 20 | 370.225 | 0.073 | | |
| Total | 23 | 5,089.804 | 1.000 | | |

**Comparison: control_vs_WCS315**

| Factor | DF | Sum Sq | $R^2$ | F-value | p-value |
|---|---|---|---|---|---|
| **Pseudomonas** | **1** | **430.127** | **0.105** | **25.169** | **0.001** |
| **Time** | **1** | **747.154** | **0.183** | **43.720** | **0.001** |
| **Treatment** | **1** | **2,558.234** | **0.627** | **149.697** | **0.001** |
| Residual | 20 | 341.788 | 0.084 | | |
| Total | 23 | 4,077.304 | 1.000 | | |

**Comparison: control_vs_CHA0**

| Factor | DF | Sum Sq | $R^2$ | F-value | p-value |
|---|---|---|---|---|---|
| **Pseudomonas** | **1** | **2,141.751** | **0.392** | **108.903** | **0.001** |
| **Time** | **1** | **654.522** | **0.120** | **33.281** | **0.001** |
| **Treatment** | **1** | **2,269.622** | **0.416** | **115.405** | **0.001** |
| Residual | 20 | 393.332 | 0.072 | | |
| Total | 23 | 5,459.226 | 1.000 | | |

**Comparison: WCS417_vs_WCS315**

| Factor | DF | Sum Sq | $R^2$ | F-value | p-value |
|---|---|---|---|---|---|
| **Pseudomonas** | **1** | **527.904** | **0.143** | **40.120** | **0.001** |
| **Time** | **1** | **564.717** | **0.152** | **42.917** | **0.001** |
| **Treatment** | **1** | **2,348.247** | **0.634** | **178.462** | **0.001** |
| Residual | 20 | 263.164 | 0.071 | | |
| Total | 23 | 3,704.033 | 1.000 | | |

**Comparison: WCS417_vs_CHA0**

| Factor | DF | Sum Sq | $R^2$ | F-value | p-value |
|---|---|---|---|---|---|
| Pseudomonas | 1 | 10.005 | 0.004 | 0.709 | 0.443 |
| **Time** | **1** | **486.864** | **0.170** | **34.505** | **0.001** |
| **Treatment** | **1** | **2,077.366** | **0.727** | **147.228** | **0.001** |
| Residual | 20 | 282.197 | 0.099 | | |
| Total | 23 | 2,856.432 | 1.000 | | |

**Comparison: WCS315_vs_CHA0**

| Factor | DF | Sum Sq | $R^2$ | F-value | p-value |
|---|---|---|---|---|---|
| **Pseudomonas** | **1** | **652.436** | **0.165** | **40.331** | **0.001** |
| **Time** | **1** | **574.929** | **0.145** | **35.540** | **0.001** |
| **Treatment** | **1** | **2,415.080** | **0.609** | **149.290** | **0.001** |
| Residual | 20 | 323.543 | 0.082 | | |
| Total | 23 | 3,965.988 | 1.000 | | |

**Table S24** List of genes retrieved from DESEq2 of roots of Arabidopsis thaliana exposed to VOCs of *Ps*WCS417, *Ps*CHA0 and *Ps*WCS315 for 24 and 48 h. Ths table is provided in excel.

**Table S25** Gene Ontology (GO) annotations for genes uniquely differentially expressed in seedlings exposed to *Ps*WCS417 and *Ps*CHA0 under drought conditions. Only genes with threshold $p < 0.05$ and |log2fold| ≥ 1 were included. For each gene, the Arabidopsis gene identifier (TAIR ID) and gene symbol are shown, together with associated GO terms grouped by Biological Process, Molecular Function, and Cellular Component. GO annotations were retrieved from the TAIR Gene Ontology Annotation (GOA) database. This table is provided in excel.

**Table S26** Type II Analysis of Deviance (Wald Chi-Square Test) for Seedling Biomass in Col-0 seedlings grown in control conditions. Results from a generalized linear mixed model (GLMM) assessing the effect of treatment on total seedling biomass (mg) when grown in control conditions. The model included treatment as a fixed effect and plate ID as a random effect.

| Factor | Chi² | DF | p-value | Significance |
|---|---|---|---|---|
| treatment | 0.17 | 1 | 0.678 | n.s. |

**Table S27** Type II Analysis of Deviance (Wald Chi-Square Test) for Seedling Biomass in Col-0 seedlings grown in drought conditions. Results from a generalized linear mixed model (GLMM) assessing the effect of treatment on total seedling biomass (mg) when grown in drought conditions. The model included treatment as a fixed effect and plate ID as a random effect.

| Factor | Chi² | DF | p-value | Significance |
|---|---|---|---|---|
| **treatment** | **7.03** | **1** | **0.008** | ** |

**Table S28** Type II Analysis of Deviance (Wald Chi-Square Test) for Seedling Biomass in *bcat4* seedlings grown in control conditions. Results from a generalized linear mixed model (GLMM) assessing the effect of treatment on total seedling biomass (mg) when grown in control conditions. The model included treatment as a fixed effect and plate ID as a random effect.

| Factor | Chi² | DF | p-value | Significance |
|---|---|---|---|---|
| **treatment** | **6.41** | **1** | **0.011** | * |

**Table S29** Type II Analysis of Deviance (Wald Chi-Square Test) for Seedling Biomass in *bcat4* seedlings grown in drought conditions. Results from a generalized linear mixed model (GLMM) assessing the effect of treatment on total seedling biomass (mg) when grown in drought conditions. The model included treatment as a fixed effect and plate ID as a random effect.

| Factor | Chi² | DF | p-value | Significance |
|---|---|---|---|---|
| treatment | 12.81 | 1 | <0.001 | *** |

**Table S30** Type II Analysis of Deviance (Wald Chi-Square Test) for Seedling Biomass in *snrk2.2/2.3* seedlings grown in control conditions. Results from a generalized linear mixed model (GLMM) assessing the effect of treatment on total seedling biomass (mg) when grown in control conditions. The model included treatment as a fixed effect and plate ID as a random effect.

| Factor | Chi² | DF | p-value | Significance |
|---|---|---|---|---|
| treatment | 36.03 | 1 | <0.001 | *** |

**Table S31** Type II Analysis of Deviance (Wald Chi-Square Test) for Seedling Biomass in *snrk2.2/2.3* seedlings grown in drought conditions. Results from a generalized linear mixed model (GLMM) assessing the effect of treatment on total seedling biomass (mg) when grown in drought conditions. The model included treatment as a fixed effect and plate ID as a random effect.

| Factor | $Chi^2$ | DF | p-value | Significance |
|---|---|---|---|---|
| treatment | 17.51 | 1 | <0.001 | *** |

**Table S32** Type II Analysis of Deviance (Wald Chi-Square Test) for Seedling Biomass in *sweet11/12* seedlings grown in control conditions. Results from a generalized linear mixed model (GLMM) assessing the effect of treatment on total seedling biomass (mg) when grown in control conditions. The model included treatment as a fixed effect and plate ID as a random effect.

| Factor | $Chi^2$ | DF | p-value | Significance |
|---|---|---|---|---|
| treatment | 0.21 | 1 | 0.651 | n.s. |

**Table S33** Type II Analysis of Deviance (Wald Chi-Square Test) for Seedling Biomass in *sweet11/12* seedlings grown in drought conditions. Results from a generalized linear mixed model (GLMM) assessing the effect of treatment on total seedling biomass (mg) when grown in drought conditions. The model included treatment as a fixed effect and plate ID as a random effect.

| Factor | $Chi^2$ | DF | p-value | Significance |
|---|---|---|---|---|
| treatment | 0.096 | 1 | 0.096 | n.s. |

**Table S34** Type II Analysis of Deviance (Wald Chi-Square Test) for Primary Root Length in Col-0 seedlings grown in control conditions. Results from a generalized linear mixed model (GLMM) assessing the effect of treatment on primary root length (mm) when grown in control conditions. The model included treatment as a fixed effect and plate ID as a random effect.

| Factor | $Chi^2$ | DF | p-value | Significance |
|---|---|---|---|---|
| treatment | 0 | 1 | 0.968 | n.s. |

**Table S35** Type II Analysis of Deviance (Wald Chi-Square Test) for Primary Root Length in Col-0 seedlings grown in drought conditions. Results from a generalized linear mixed model (GLMM) assessing the effect of treatment on primary root length (mm) when grown in drought conditions. The model included treatment as a fixed effect and plate ID as a random effect.

| Factor | Chi² | DF | p-value | Significance |
|---|---|---|---|---|
| treatment | 0.05 | 1 | 0.824 | n.s. |

**Table S36** Type II Analysis of Deviance (Wald Chi-Square Test) for Primary Root Length in *bcat4* seedlings grown in control conditions. Results from a generalized linear mixed model (GLMM) assessing the effect of treatment on primary root length (mm) when grown in control conditions. The model included treatment as a fixed effect and plate ID as a random effect.

| Factor | Chi² | DF | p-value | Significance |
|---|---|---|---|---|
| treatment | 1.24 | 1 | 0.265 | n.s. |

**Table S37** Type II Analysis of Deviance (Wald Chi-Square Test) for Primary Root Length in *bcat4* seedlings grown in drought conditions. Results from a generalized linear mixed model (GLMM) assessing the effect of treatment on primary root length (mm) when grown in drought conditions. The model included treatment as a fixed effect and plate ID as a random effect.

| Factor | Chi² | DF | p-value | Significance |
|---|---|---|---|---|
| **treatment** | **7.89** | **1** | **0.005** | ** |

**Table S38** Type II Analysis of Deviance (Wald Chi-Square Test) for Primary Root Length in *snrk2.2/2.3* seedlings grown in control conditions. Results from a generalized linear mixed model (GLMM) assessing the effect of treatment on primary root length (mm) when grown in control conditions. The model included treatment as a fixed effect and plate ID as a random effect.

| Factor | Chi² | DF | p-value | Significance |
|---|---|---|---|---|
| **treatment** | **20.35** | **1** | **<0.001** | *** |

**Table S39** Type II Analysis of Deviance (Wald Chi-Square Test) for Primary Root Length in *snrk2.2/2.3* seedlings grown in drought conditions. Results from a generalized linear mixed model (GLMM) assessing the effect of treatment on primary root length (mm) when grown in drought conditions. The model included treatment as a fixed effect and plate ID as a random effect.

| Factor | Chi² | DF | p-value | Significance |
|---|---|---|---|---|
| **treatment** | **10.33** | **1** | **0.001** | ** |

**Table S40** Type II Analysis of Deviance (Wald Chi-Square Test) for Primary Root Length in *sweet11/12* seedlings grown in control conditions. Results from a generalized linear mixed model (GLMM) assessing the effect of treatment on primary root length (mm) when grown in control conditions. The model included treatment as a fixed effect and plate ID as a random effect.

| Factor | Chi² | DF | p-value | Significance |
|---|---|---|---|---|
| treatment | 0.27 | 1 | 0.601 | n.s. |

**Table S41** Type II Analysis of Deviance (Wald Chi-Square Test) for Primary Root Length in *sweet11/12* seedlings grown in drought conditions. Results from a generalized linear mixed model (GLMM) assessing the effect of treatment on primary root length (mm) when grown in drought conditions. The model included treatment as a fixed effect and plate ID as a random effect.

| Factor | Chi² | DF | p-value | Significance |
|---|---|---|---|---|
| treatment | 0.02 | 1 | 0.892 | n.s. |

**Table S42** Type II Analysis of Deviance (Wald Chi-Square Test) for Lateral Root Count in Col-0 seedlings grown in control conditions. Results from a generalized linear mixed model (GLMM) assessing the effect of treatment on lateral root count when grown in control conditions. The model included treatment as a fixed effect and plate ID as a random effect.

| Factor | Chi² | DF | p-value | Significance |
|---|---|---|---|---|
| **treatment** | **5.22** | **1** | **0.022** | ***** |

**Table S43** Type II Analysis of Deviance (Wald Chi-Square Test) for Lateral Root Count in Col-0 seedlings grown in drought conditions. Results from a generalized linear mixed model (GLMM) assessing the effect of treatment on lateral root count when grown in drought conditions. The model included treatment as a fixed effect and plate ID as a random effect.

| Factor | Chi² | DF | p-value | Significance |
|---|---|---|---|---|
| treatment | 3.5 | 1 | 0.061 | n.s. |

**Table S44** Type II Analysis of Deviance (Wald Chi-Square Test) for Lateral Root Count in *bcat4* seedlings grown in control conditions. Results from a generalized linear mixed model (GLMM) assessing the effect of treatment on lateral root count when grown in control conditions. The model included treatment as a fixed effect and plate ID as a random effect.

| Factor | Chi² | DF | p-value | Significance |
|---|---|---|---|---|
| treatment | 3.68 | 1 | 0.055 | n.s. |

**Table S45** Type II Analysis of Deviance (Wald Chi-Square Test) for Lateral Root Count in *bcat4* seedlings grown in drought conditions. Results from a generalized linear mixed model (GLMM) assessing the effect of treatment on lateral root count when grown in drought conditions. The model included treatment as a fixed effect and plate ID as a random effect.

| Factor | Chi² | DF | p-value | Significance |
|---|---|---|---|---|
| **treatment** | **7.36** | **1** | **0.007** | ** |

**Table S46** Type II Analysis of Deviance (Wald Chi-Square Test) for Lateral Root Count in *snrk2.2/2.3* seedlings grown in control conditions. Results from a generalized linear mixed model (GLMM) assessing the effect of treatment on lateral root count when grown in control conditions. The model included treatment as a fixed effect and plate ID as a random effect.

| Factor | Chi² | DF | p-value | Significance |
|---|---|---|---|---|
| treatment | 27.07 | 1 | <0.001 | *** |

**Table S47** Type II Analysis of Deviance (Wald Chi-Square Test) for Lateral Root Count in *snrk2.2/2.3* seedlings grown in drought conditions. Results from a generalized linear mixed model (GLMM) assessing the effect of treatment on lateral root count when grown in drought conditions. The model included treatment as a fixed effect and plate ID as a random effect.

| Factor | Chi² | DF | p-value | Significance |
|---|---|---|---|---|
| treatment | 16.62 | 1 | <0.001 | *** |

**Table S48** Type II Analysis of Deviance (Wald Chi-Square Test) for Lateral Root Count in *sweet11/12* seedlings grown in control conditions. Results from a generalized linear mixed model (GLMM) assessing the effect of treatment on lateral root count when grown in control conditions. The model included treatment as a fixed effect and plate ID as a random effect.

| Factor | Chi² | DF | p-value | Significance |
|---|---|---|---|---|
| **treatment** | **8.46** | **1** | **0.004** | ** |

**Table S49** Type II Analysis of Deviance (Wald Chi-Square Test) for Lateral Root Count in *sweet11/12* seedlings grown in drought conditions**.** Results from a generalized linear mixed model (GLMM) assessing the effect of treatment on lateral root count when grown in drought conditions. The model included treatment as a fixed effect and plate ID as a random effect.

| Factor | $Chi^2$ | DF | p-value | Significance |
|---|---|---|---|---|
| treatment | 0.56 | 1 | 0.454 | n.s. |

**Table S50** Statistical analyses of coumarins detected by LC-MS. LC-MS peak areas for coumarins were log-transformed and Pareto-scaled prior to analysis. Significant differences among groups were assed using ANOVA.

| Retention time (min) | Measured [M-H]- | Calculated [M-H]- | ppm error ([M-H]-) | Elemental composition | Compound identity | ANOVA p-value |
|---|---|---|---|---|---|---|
| 1,25 | 191,0343 | 191,0350 | 4,0203 | C10H8O4 (Estimated) | Unknown | ND |
| 6,13 | 163,0408 | 163,0401 | 4,3548 | C9H8O3 | p-coumaric acid | 0,0034327 |
| 6,20 | 191,0354 | 191,0350 | 2,0938 | C10H8O4 | Scopoletin | 0,0381631 |
| 7,17 | 163,0406 | 163,0401 | 3,0667 | C9H8O3 | 2-Hydroxycinnamic acid | 0,0000071 |
| 7,21 | 339,0721 | 339,0722 | -0,2949 | C15H16O9 | 6,7-Dihydroxycoumarin-6-glucoside | 0,270908 |

**Table S51** Statistical analyses of glucosinolates detected by LC-MS. LC-MS peak areas for glucosinolates were log-transformed and Pareto-scaled prior to analysis. Significant differences among groups were assed using ANOVA.

| Retention time (min) | Measured [M-H]- | Calculated [M-H]- | ppm error ([M-H]-) | Elemental composition | Compound identity | ANOVA p-value | GLS class |
|---|---|---|---|---|---|---|---|
| 1,25 | 436,0413 | 436,0411 | 0,291261953 | C12H23NO10S3 | **4-methylsulfinylbutyl_glucosinolate** | **0,002477705** | Aliphatic |
| 1,27 | 450,0579 | 450,0568 | 2,463995153 | C13H25NO10S3 | **5-methylsulfinylpentyl_glucosinolate** | **0,002086895** | Aliphatic |
| 1,83 | 464,0726 | 464,0724 | 0,361739375 | C14H27NO10S3 | **6-methylsulfinylhexyl_glucosinolate** | **0,007392043** | Aliphatic |
| 2,22 | 463,0486 | 463,0487 | -0,165708606 | C16H20N2O10S2 | 4-hydroxy-3-indolylmethyl-glucosinolate | 0,316195159 | Indolic |
| 3,38 | 478,0880 | 478,0881 | -0,146816435 | C15H29NO10S3 | **7-methylsulfinylheptyl_glucosinolate** | **0,018854876** | Aliphatic |
| 4,13 | 447,0540 | 447,0537 | 0,673047535 | C16H20N2O9S2 | 3-indolymethyl_glucosinolate | 0,853512646 | Indolic |
| 4,64 | 492,1036 | 492,1037 | -0,256563077 | C16H31NO10S3 | 8-Methylsulfinyloctyl_glucosinolate | 0,548299901 | Aliphatic |
| 5,36 | 477,0644 | 477,0643 | 0,107332263 | C17H22N2O10S2 | **4-methoxy-3-indolylmethyl-glucosinolate** | **0,002450611** | Indolic |
| 6,55 | 477,0643 | 477,0643 | -0,00225967 | C17H22N2O10S2 | 1-methoxy-3-indolylmethyl-glucosinolate | 0,679926521 | Indolic |
| 6,56 | 448,0790 | 448,0775 | 3,32864913 | C14H27NO9S3 | 6-methylthiohexylglucosinolate | 0,409626968 | Aliphatic |
| 7,57 | 494,0800 | 494,0796 | 0,729624422 | C18H25NO11S2 | 4-benzoyloxybutylglucosinolate | 0,346719727 | Aliphatic |
| 8,08 | 462,0933 | 462,0932 | 0,202185162 | C15H29NO9S3 | 7-methylthioheptyl_glucosinolate | 0,525417401 | Aliphatic |
| 8,24 | 416,1029 | 416,1054 | -6,109828108 | C14H27O9NS2 | heptylglucosinolate | 0,269423587 | Aliphatic |
| 9,54 | 476,1090 | 476,1088 | 0,462843605 | C16H31NO9S3 | 8-Methylthiooctyl_glucosinolate | 0,404941472 | Aliphatic |
| 10,61 | 492,0638 | 492,0640 | -0,271190301 | C18H23NO11S2 | **2-benzoyloxy-3-butenylglucosinolate** | **0,034825245** | Aliphatic |

**Table S52** Two-way analysis of variance (ANOVA) testing the effects of treatment (*Pseudomonas* WCS417 VOCs exposure), water regime (well-watered vs. drought), and their interaction on glucose, fructose, and sucrose concentrations in roots and shoots of axenically grown Arabidopsis thaliana seedlings.

| Sugar | Tissue | term | df | sumsq | meansq | statistic | p.value |
|---|---|---|---|---|---|---|---|
| **Glucose** | **Root** | **Treatment** | **1** | **0.4080** | **0.4080** | **10.08** | **0.0088** |
| | | **Water_regime** | **1** | **1.3749** | **1.3749** | **33.97** | **< 0.001** |
| | | Treatment:Water_regime | 1 | 0.1027 | 0.1027 | 2.53 | 0.1394 |
| | | Residuals | 11 | 0.4452 | 0.0404 | | |
| | **Shoot** | Treatment | 1 | 0.0882 | 0.0882 | 1.20 | 0.2947 |
| | | **Water_regime** | **1** | **0.8600** | **0.8600** | **11.69** | **0.0050** |
| | | Treatment:Water_regime | 1 | 0.0017 | 0.0017 | 0.02 | 0.8805 |
| | | Residuals | 12 | 0.8821 | 0.0735 | | |
| **Fructose** | **Root** | **Treatment** | **1** | **0.0662** | **0.0662** | **6.58** | **0.0262** |
| | | **Water_regime** | **1** | **1.2889** | **1.2889** | **128.24** | **< 0.001** |
| | | Treatment:Water_regime | 1 | 0.0079 | 0.0079 | 0.78 | 0.3936 |
| | | Residuals | 11 | 0.1105 | 0.0100 | | |
| | **Shoot** | Treatment | 1 | 0.0882 | 0.0882 | 1.20 | 0.2947 |
| | | **Water_regime** | **1** | **0.8600** | **0.8600** | **11.69** | **0.0050** |
| | | Treatment:Water_regime | 1 | 0.0017 | 0.0017 | 0.02 | 0.8805 |

| Sugar | Tissue | term | df | sumsq | meansq | statistic | p.value |
|---|---|---|---|---|---|---|---|
| | | Residuals | 12 | 0.8821 | 0.0735 | | |
| **Sucrose** | **Root** | **Treatment** | **1** | 0.1291 | 0.1291 | **14.70** | **0.0027** |
| | | **Water_regime** | **1** | 0.9926 | 0.9926 | **113.01** | **< 0.001** |
| | | **Treatment:Water_regime** | **1** | 0.0957 | 0.0957 | **10.89** | **0.0070** |
| | | Residuals | 11 | 0.0966 | 0.0087 | | |
| | **Shoot** | Treatment | 1 | 0.0053 | 0.0053 | 0.27 | 0.6077 |
| | | **Water_regime** | **1** | 2.9770 | 2.9770 | **154.38** | **< 0.001** |
| | | Treatment:Water_regime | 1 | 0.0187 | 0.0187 | 0.97 | 0.3433 |
| | | Residuals | 12 | 0.2313 | 0.0192 | | |

**Table S53** Tukey HSD post-hoc pairwise comparison following two-way ANOVA, showing pairwise comparisons for sucrose concentrations on roots for axenically grown Arabidopsis seedlings.

| diff | lwr | upr | p adj | Comparison |
|---|---|---|---|---|
| -0.0022 | -0.9994 | 0.9949 | 0.9999 | pseudomonas:control-mock:control |
| **3.3280** | **2.3307** | **4.3252** | **< 0.001** | **mock:drought-mock:control** |
| **1.7155** | **0.6384** | **2.7927** | **0.0026** | **pseudomonas:drought-mock:control** |
| **3.3302** | **2.3330** | **4.3274** | **< 0.001** | **mock:drought-pseudomonas:control** |
| **1.7178** | **0.6406** | **2.7949** | **0.0026** | **pseudomonas:drought-pseudomonas:control** |
| **-1.6124** | **-2.6895** | **-0.5352** | **0.0042** | **pseudomonas:drought-mock:drought** |

**Table S54** Kruskal-Wallis Test Result on Shoot Fresh Biomass of *Brassica oleracea***.**

| Kruskal-Wallis $\chi^2$ | df | p-value |
|---|---|---|
| 48.92 | 7 | < 0.001 |

**Table S55** Pairwise Wilcoxon Test (BH Adjusted) Result on Shoot Fresh Biomass of *Brassica oleracea*.

| Comparison | Group | p.value |
|---|---|---|
| Control-DR | Control-DR | 1.000 |
| **Control-DR** | **Control-WW** | **0.001** |
| **Control-DR** | **PsCHA0-DR** | **0.003** |
| **Control-DR** | **PsCHA0-WW** | **0.001** |
| **Control-DR** | **PsWCS315-DR** | **0.003** |
| **Control-DR** | **PsWCS315-WW** | **0.001** |
| **Control-DR** | **PsWCS417-DR** | **0.001** |
| **Control-DR** | **PsWCS417-WW** | **0.001** |
| **Control-WW** | **Control-DR** | **0.001** |
| Control-WW | Control-WW | 1.000 |
| **Control-WW** | **PsCHA0-DR** | **0.033** |
| **Control-WW** | **PsCHA0-WW** | **0.001** |
| Control-WW | PsWCS315-DR | 0.138 |
| **Control-WW** | **PsWCS315-WW** | **0.001** |
| **Control-WW** | **PsWCS417-DR** | **0.044** |

| Comparison | Group | p.value |
|---|---|---|
| **Control-WW** | **PsWCS417-WW** | **0.001** |
| **PsCHA0-DR** | **Control-DR** | **0.003** |
| **PsCHA0-DR** | **Control-WW** | **0.033** |
| PsCHA0-DR | PsCHA0-DR | 1.000 |
| **PsCHA0-DR** | **PsCHA0-WW** | **0.001** |
| PsCHA0-DR | PsWCS315-DR | 0.059 |
| **PsCHA0-DR** | **PsWCS315-WW** | **0.001** |
| PsCHA0-DR | PsWCS417-DR | 0.269 |
| **PsCHA0-DR** | **PsWCS417-WW** | **0.001** |
| **PsCHA0-WW** | **Control-DR** | **0.001** |
| **PsCHA0-WW** | **Control-WW** | **0.001** |
| **PsCHA0-WW** | **PsCHA0-DR** | **0.001** |
| PsCHA0-WW | PsCHA0-WW | 1.000 |
| **PsCHA0-WW** | **PsWCS315-DR** | **0.001** |
| **PsCHA0-WW** | **PsWCS315-WW** | **0.002** |
| **PsCHA0-WW** | **PsWCS417-DR** | **0.001** |
| **PsCHA0-WW** | **PsWCS417-WW** | **0.044** |
| **PsWCS315-DR** | **Control-DR** | **0.003** |
| PsWCS315-DR | Control-WW | 0.138 |
| PsWCS315-DR | PsCHA0-DR | 0.059 |
| **PsWCS315-DR** | **PsCHA0-WW** | **0.001** |
| PsWCS315-DR | PsWCS315-DR | 1.000 |
| **PsWCS315-DR** | **PsWCS315-WW** | **0.001** |
| PsWCS315-DR | PsWCS417-DR | 0.383 |
| **PsWCS315-DR** | **PsWCS417-WW** | **0.001** |
| **PsWCS315-WW** | **Control-DR** | **0.001** |
| **PsWCS315-WW** | **Control-WW** | **0.001** |
| **PsWCS315-WW** | **PsCHA0-DR** | **0.001** |
| **PsWCS315-WW** | **PsCHA0-WW** | **0.002** |
| **PsWCS315-WW** | **PsWCS315-DR** | **0.001** |
| PsWCS315-WW | PsWCS315-WW | 1.000 |
| **PsWCS315-WW** | **PsWCS417-DR** | **0.001** |
| **PsWCS315-WW** | **PsWCS417-WW** | **0.002** |
| **PsWCS417-DR** | **Control-DR** | **0.001** |
| **PsWCS417-DR** | **Control-WW** | **0.044** |
| PsWCS417-DR | PsCHA0-DR | 0.269 |
| **PsWCS417-DR** | **PsCHA0-WW** | **0.001** |
| PsWCS417-DR | PsWCS315-DR | 0.383 |
| **PsWCS417-DR** | **PsWCS315-WW** | **0.001** |
| PsWCS417-DR | PsWCS417-DR | 1.000 |
| **PsWCS417-DR** | **PsWCS417-WW** | **0.001** |
| **PsWCS417-WW** | **Control-DR** | **0.001** |
| **PsWCS417-WW** | **Control-WW** | **0.001** |

| Comparison | Group | p.value |
|---|---|---|
| **PsWCS417-WW** | **PsCHA0-DR** | **0.001** |
| **PsWCS417-WW** | **PsCHA0-WW** | **0.044** |
| **PsWCS417-WW** | **PsWCS315-DR** | **0.001** |
| **PsWCS417-WW** | **PsWCS315-WW** | **0.002** |
| **PsWCS417-WW** | **PsWCS417-DR** | **0.001** |
| PsWCS417-WW | PsWCS417-WW | 1.000 |

**Table S56** Compact Letter Display (CLD) on Shoot Fresh Biomass of *Brassica oleracea*.

| Group | Letters |
|---|---|
| Control-DR | a |
| Control-WW | b |
| PsCHA0-DR | c |
| PsCHA0-WW | d |
| PsWCS315-DR | bc |
| PsWCS315-WW | e |
| PsWCS417-DR | c |
| PsWCS417-WW | f |

**Table S57** Kruskal-Wallis Test Result on *Brassica oleracea* Shoot Water Content.

| Kruskal-Wallis $\chi^2$ | df | p-value |
|---|---|---|
| 23.61 | 7 | 0.001 |

**Table S58** Pairwise Wilcoxon Test (BH Adjusted) on *Brassica oleracea* Shoot Water Content.

| Comparison | Group | p.value |
|---|---|---|
| Control-DR | Control-DR | 1.000 |
| Control-DR | Control-WW | 0.525 |
| **Control-DR** | **PsCHA0-DR** | **0.019** |
| **Control-DR** | **PsCHA0-WW** | **0.016** |
| **Control-DR** | **PsWCS315-DR** | **0.019** |
| **Control-DR** | **PsWCS315-WW** | **0.019** |
| **Control-DR** | **PsWCS417-DR** | **0.019** |
| **Control-DR** | **PsWCS417-WW** | **0.016** |
| Control-WW | Control-DR | 0.525 |

| Comparison | Group | p.value |
|---|---|---|
| Control-WW | Control-WW | 1.000 |
| Control-WW | PsCHA0-DR | 0.069 |
| **Control-WW** | **PsCHA0-WW** | **0.019** |
| Control-WW | PsWCS315-DR | 0.105 |
| Control-WW | PsWCS315-WW | 0.120 |
| Control-WW | PsWCS417-DR | 0.098 |
| **Control-WW** | **PsWCS417-WW** | **0.029** |
| **PsCHA0-DR** | **Control-DR** | **0.019** |
| PsCHA0-DR | Control-WW | 0.069 |
| PsCHA0-DR | PsCHA0-DR | 1.000 |
| PsCHA0-DR | PsCHA0-WW | 0.468 |
| PsCHA0-DR | PsWCS315-DR | 0.900 |
| PsCHA0-DR | PsWCS315-WW | 0.755 |
| PsCHA0-DR | PsWCS417-DR | 0.902 |
| PsCHA0-DR | PsWCS417-WW | 0.468 |
| **PsCHA0-WW** | **Control-DR** | **0.016** |
| **PsCHA0-WW** | **Control-WW** | **0.019** |
| PsCHA0-WW | PsCHA0-DR | 0.468 |
| PsCHA0-WW | PsCHA0-WW | 1.000 |
| PsCHA0-WW | PsWCS315-DR | 0.468 |
| PsCHA0-WW | PsWCS315-WW | 0.449 |
| PsCHA0-WW | PsWCS417-DR | 0.468 |
| PsCHA0-WW | PsWCS417-WW | 0.902 |
| **PsWCS315-DR** | **Control-DR** | **0.019** |
| PsWCS315-DR | Control-WW | 0.105 |
| PsWCS315-DR | PsCHA0-DR | 0.900 |
| PsWCS315-DR | PsCHA0-WW | 0.468 |
| PsWCS315-DR | PsWCS315-DR | 1.000 |
| PsWCS315-DR | PsWCS315-WW | 0.818 |
| PsWCS315-DR | PsWCS417-DR | 0.818 |
| PsWCS315-DR | PsWCS417-WW | 0.468 |
| **PsWCS315-WW** | **Control-DR** | **0.019** |
| PsWCS315-WW | Control-WW | 0.120 |
| PsWCS315-WW | PsCHA0-DR | 0.755 |
| PsWCS315-WW | PsCHA0-WW | 0.449 |
| PsWCS315-WW | PsWCS315-DR | 0.818 |
| PsWCS315-WW | PsWCS315-WW | 1.000 |
| PsWCS315-WW | PsWCS417-DR | 0.755 |
| PsWCS315-WW | PsWCS417-WW | 0.468 |
| **PsWCS417-DR** | **Control-DR** | **0.019** |
| PsWCS417-DR | Control-WW | 0.098 |
| PsWCS417-DR | PsCHA0-DR | 0.902 |
| PsWCS417-DR | PsCHA0-WW | 0.468 |

| Comparison | Group | p.value |
|---|---|---|
| PsWCS417-DR | PsWCS315-DR | 0.818 |
| PsWCS417-DR | PsWCS315-WW | 0.755 |
| PsWCS417-DR | PsWCS417-DR | 1.000 |
| PsWCS417-DR | PsWCS417-WW | 0.525 |
| **PsWCS417-WW** | **Control-DR** | **0.016** |
| **PsWCS417-WW** | **Control-WW** | **0.029** |
| PsWCS417-WW | PsCHA0-DR | 0.468 |
| PsWCS417-WW | PsCHA0-WW | 0.902 |
| PsWCS417-WW | PsWCS315-DR | 0.468 |
| PsWCS417-WW | PsWCS315-WW | 0.468 |
| PsWCS417-WW | PsWCS417-DR | 0.525 |
| PsWCS417-WW | PsWCS417-WW | 1.000 |

**Table S59** Compact Letter Display (CLD) on *Brassica oleracea* Shoot Water Content.

| Group | Letters |
|---|---|
| Control-DR | a |
| Control-WW | ab |
| PsCHA0-DR | bc |
| PsCHA0-WW | c |
| PsWCS315-DR | bc |
| PsWCS315-WW | bc |
| PsWCS417-DR | bc |
| PsWCS417-WW | c |

**Table S60** PERMANOVA Results on root microbiome. Permutational multivariate analysis of variance (PERMANOVA) on Bray-Curtis dissimilarities of microbial community composition. The table summarizes the effects of *Pseudomonas* strain and water regime on bacterial community structure across samples. Significant p-values (p , 0.05) are highlighted in bold.

| Factor | DF | Sum Sq | $R^2$ | F-value | p-value |
|---|---|---|---|---|---|
| ***Pseudomonas* strain** | **3** | **0.19** | **0.084** | **2.05** | **0.005** |
| **Water regime** | **1** | **0.52** | **0.231** | **16.87** | **0.001** |
| Residual | 50 | 1.53 | 0.686 | | NA |
| Total | 54 | 2.23 | 1.000 | | NA |

**Table S61** ANOVA Results for Shannon Diversity on root microbiome**.**

| | | DF | Sum Sq | Mean Sq | F-value | p-value |
|---|---|---|---|---|---|---|
| Control | *Pseudomonas* strain | 3 | 0.15 | 0.05083305 | 1.84 | 0.166 |
| | Residuals | 24 | 0.66 | 0.02756219 | | NA |
| Drought | *Pseudomonas* strain | 3 | 0.05 | 0.01546561 | 0.88 | 0.467 |
| | Residuals | 23 | 0.41 | 0.01763261 | | NA |

**Table S62** Kruskal-Wallis Test Results Simpson Diversity on root microbiome**.**

| Test | Comparison | Statistic | p-value |
|---|---|---|---|
| **Kruskal-Wallis** | | **10.52** | **0.015** |

**Table S63** Dunn's Post-Hoc Test Results

| Comparison | Z-value | Adjusted p-value | p-value | Test |
|---|---|---|---|---|
| CHA0 - Control | 0.4548588 | 0.649210806 | 0.97381621 | Dunn's Test |
| CHA0 - WCS315 | 0.1624496 | 0.870951830 | 0.87095183 | Dunn's Test |
| Control - WCS315 | -0.2924092 | 0.769973740 | 0.92396849 | Dunn's Test |
| **CHA0 - WCS417** | **2.8266227** | **0.004704172** | **0.02822503** | **Dunn's Test** |
| **Control - WCS417** | **2.3717639** | **0.017703400** | **0.03540680** | **Dunn's Test** |
| **WCS315 - WCS417** | **2.6641731** | **0.007717783** | **0.02315335** | **Dunn's Test** |

**Table S64** ANOVA Results for Simpson Diversity

| DF | Sum Sq | Mean Sq | F-value | p-value |
|---|---|---|---|---|
| 3 | 0 | 0.0000003157686 | 0.69 | 0.566 |
| 23 | 0 | 0.0000004561493 | | NA |

**Table S65** ANOVA Results for Observed Diversity

| | | DF | Sum Sq | Mean Sq | F-value | p-value |
|---|---|---|---|---|---|---|
| Control | *Pseudomonas* strain | 3 | 1,658,911 | 552,970.4 | 0.71 | 0.555 |
| | Residuals | 24 | 18,648,739 | 777,030.8 | | NA |

| | | DF | Sum Sq | Mean Sq | F-value | p-value |
|---|---|---|---|---|---|---|
| Drought | *Pseudomonas* strain | 3 | 1,239,276 | 413,092.1 | 1.06 | 0.387 |
| | Residuals | 23 | 9,000,667 | 391,333.4 | | NA |

**Table S66** List of ASVs retrieved from ANCOM-BC differential abundance analyses performed on *Brassica oleracea* root microbiome 16S amplicon sequencing data. This tabe is provided as an excel file.
